## Supplementary material for "On the interplay between lipids and asymmetric dynamics of an NBS degenerate ABC transporter": Electronic Supplementary information

*Inserm U1248 Pharmacology & Transplantation, Univ. Limoges, 2 rue du Prof. Descottes, 87000 F-Limoges, France*

**Contact information**

Dr. Florent Di Meo

INSERM U1248 Pharmacology & Transplantation, Univ. Limoges,

2 rue du Prof. Descottes,

87000 F-Limoges, France

Supplementary Figure 16. Representative NBD communities from Network Analyses calculated from MD simulations performed in POPC:POPE:Chol (2:1:1) simulations. a) IF apo *b*MRP1, b) *b*MRP1-$(ATP)_2$, c) *b*MRP1-LTX, d) *b*MRP1-LTX-$(ATP)_2$ and e) OF *b*MRP1-$(ATP)_2$. NBD1 and NBD2 are coloured yellow and cyan, respectively. Orange and blue colours show the TMH parts involved in the communities. The Walker A community is purple, while the signature sequence community silver. NBD dimers are shown from upper view. ........ 55

Supplementary Figure 17. Non-covalent interaction energies between LTX and *b*MRP1 extracted from Coulomb (top) and van der Waals (bottom) potentials obtained from MD simulations performed in a) POPC:POPE:Chol (2:1:1), b) POPC:Chol (3:1) and c) POPC lipid bilayers. ........ 56

Supplementary Figure 18. Calculated per-residue betweennesses in the allosteric pathway from substrate binding pocket to NBS1 (left) and NBS2 (right). a) IF apo *b*MRP1, b) *b*MRP1-$(ATP)_2$, c) *b*MRP1-LTX, d) *b*MRP1-LTX-$(ATP)_2$ and e) OF *b*MRP1-$(ATP)_2$ systems embedded in POPC. ........ 57

Supplementary Figure 19. Calculated per-residue betweennesses in the allosteric pathway from substrate binding pocket to NBS1 (left) and NBS2 (right). a) IF apo *b*MRP1, b) *b*MRP1-$(ATP)_2$, c) *b*MRP1-LTX, d) *b*MRP1-LTX-$(ATP)_2$ and e) OF *b*MRP1-$(ATP)_2$ systems embedded in POPC:Chol (3:1). ........ 58

Supplementary Figure 20. Calculated per-residue betweennesses in the allosteric pathway from substrate binding pocket to NBS1 (left) and NBS2 (right). a) IF apo *b*MRP1, b) *b*MRP1-$(ATP)_2$, c) *b*MRP1-LTX, d) *b*MRP1-LTX-$(ATP)_2$ and e) OF *b*MRP1-$(ATP)_2$ systems embedded in POPC:POPE:Chol (2:1:1). ........ 59

Supplementary Figure 21. Average normalized information flow through *b*MRP1 protein regarding the allosteric communication from substrate binding pocket to NBS1 and NBS2 for IF apo *b*MRP1 and *b*MRP1-LTX system in POPC:POPE:Chol (2:1:1) ........ 60

Supplementary Figure 22. ABC conformational space of trajectories in POPC compared to resolved ABC transporters. ........ 61

Supplementary Figure 23. ABC conformational space of trajectories in POPC:Chol (3:1) compared to resolved ABC transporters. ........ 62

Supplementary Figure 24. Calculated z-dependent pore radii for all systems investigated in the present study. Replica 1, 2 and 3 are coloured red, blue and yellow, respectively. ........ 43

Supplementary Figure 25. Free energy landscapes according to NBD distance and NBD twist. obtained in different lipid bilayers using the InfleCS approach for IF systems, embedded in different lipid bilayers. a) IF apo *b*MRP1, b) *b*MRP1-$(ATP)_2$, c) *b*MRP1-LTX, d) *b*MRP1-LTX-$(ATP)_2$ and e) OF *b*MRP1-$(ATP)_2$........ 63

Supplementary Figure 26. Free energy landscapes according to NBD twist and EC angle obtained in different lipid bilayers using the InfleCS approach for OF *b*MRP1-$(ATP)_2$ systems........ 64

Supplementary Figure 27. Tilt angles with respect to membrane normal axis of different transmembrane helices. Since $TMD_0$ was not modelled in the present study, the conventional TMH1 to TMH12 labelling for type I ABC transporter was used. ..65

Supplementary Figure 28. Tilt angles of bundles A, B, C and D which consist in (TMH1, 2, 10 and 11), (TMH4, 5, 7 and 8), (TMH 3 and 6) and (TMH9 and 12), respectively (see main text). ........ 66

Supplementary Figure 29. Putative substrate access to *b*MRP1 binding pocket. We here propose different possible substrate access to *b*MRP1 binding pocket depending on substrate lipid bilayer partitioning. Amphiphilic compounds including charged molecules were shown to possibly partition within the high-density polar head region of lipid bilayer membrane. Our MD simulations suggest that access to TMD binding pocket may be possible between TMH4 and TMH6 while access between TMH10 and TMH12 might less likely. ........ 71

Supplementary Figure 30. Root-mean-square deviation (RMSD) along MD simulations calculated for all the systems and for all different membrane compositions. Replica 1, 2 and 3 are coloured red, blue and yellow, respectively...... 74

Supplementary Figure 31. Atom selections for co-factor nodes for allosteric pathway calculations. Each cofactor as split into fragments (depicted in blue, green, red and pink shades, accordingly) that were used to defined nodes for allosteric pathway calculations. PC and PE lipids were split into three nodes corresponding to polar head and lipid tails. Cholesterol and $Mg^{2+}$ were considered as single node each. ATP molecule was split into three fragments (purine and sugar moieties as well as phosphate tail). LTX was split into three fragments (namely, glutathione moiety, aliphatic chain and 6-ketohexanoate moiety). ........ 78

Supplementary Figure 32. Number of H-bonds calculated between nucleotides and *b*MRP1 protein over MD simulations. H-bond interactions were counted between nucleotides and protein residues over MD simulations performed on *b*MRP1-$(ATP)_2$, *b*MRP1-LTX-$(ATP)_2$ and OF *b*MRP1-$(ATP)_2$ systems embedded in a) POPC:POPE:Chol (2:1:1), b) POPC:Chol (3:1) and c) POPC. Distance and angle cutoffs were set 3.5 Å and 120°, respectively........ 79

Supplementary Figure 33. Number of H-bonds calculated between LTX and *b*MRP1 protein over MD simulations. H-bond interactions were counted between LTX and proteins over MD simulations performed on *b*MRP1-LTX and *b*MRP1-LTX-$(ATP)_2$ systems embedded in a) POPC:POPE:Chol (2:1:1), b) POPC:Chol (3:1) and c) POPC. Distance and angle cutoffs were set 3.5 Å and 120°, respectively........ 80

Supplementary Figure 34. Overall dynamic cross-correlation matrices from MD simulations performed in POPC. a) IF apo *b*MRP1, b) *b*MRP1-$(ATP)_2$, c) *b*MRP1-LTX, d) *b*MRP1-LTX-$(ATP)_2$ and e) OF *b*MRP1-$(ATP)_2$. ........................................ 81

Supplementary Figure 35. Overall dynamic cross-correlation matrices from MD simulations performed in POPC:Chol (3:1). a) IF apo *b*MRP1, b) *b*MRP1-$(ATP)_2$, c) *b*MRP1-LTX, d) *b*MRP1-LTX-$(ATP)_2$ and e) OF *b*MRP1-$(ATP)_2$............................. 82

Supplementary Figure 36. Overall dynamic cross-correlation matrices from MD simulations performed in POPC:POPE (3:1). a) IF apo *b*MRP1 and b) OF *b*MRP1-$(ATP)_2$. ........................................................................................................ 83

Supplementary Figure 37. Overall dynamic cross-correlation matrices from MD simulations performed in POPE. a) IF apo *b*MRP1 and b) OF *b*MRP1-$(ATP)_2$.......... 84

**Supplementary Movies ........................................................................................ 85**

Supplementary Movie 1. First principal component obtained from MD simulations performed on IF apo *b*MRP1 embedded in POPC:POPE:Chol (2:1:1). PCA were performed considering only the “so-called” ABC core, i.e., TMHs and NBDs. ......... 85

Supplementary Movie 2. First principal component obtained from MD simulations performed on IF *b*MRP1-$(ATP)_2$ embedded in POPC:POPE:Chol (2:1:1). PCA were performed considering only the “so-called” ABC core, i.e., TMHs and NBDs. ......... 85

Supplementary Movie 3. First principal component obtained from MD simulations performed on IF *b*MRP1-LTX embedded in POPC:POPE:Chol (2:1:1). PCA were performed considering only the “so-called” ABC core, i.e., TMHs and NBDs. ......... 85

Supplementary Movie 4. First principal component obtained from MD simulations performed on IF *b*MRP1-LTX-$(ATP)_2$ embedded in POPC:POPE:Chol (2:1:1). PCA were performed considering only the “so-called” ABC core, i.e., TMHs and NBDs.. 85

Supplementary Movie 5. First principal component obtained from MD simulations performed on OF *b*MRP1-$(ATP)_2$ embedded in POPC:POPE:Chol (2:1:1). PCA were performed considering only the “so-called” ABC core, i.e., TMHs and NBDs. ......... 85

### Supplementary Tables

**Supplementary Table 1. Residues selected for the calculations of ABC structural parameters of ABC transporters used for mapping the ABC conformational space, namely a) EC and IC angles and b) NBD distance and twist.** Selected ABC transporters are reported according to their PBD ID. Residues were selected as explained in the Method section.

**a)** EC and IC angles

| PDB ID | EC angle | | IC angle | |
|---|---|---|---|---|
| | **Vector1 direction** | **Vector2 direction** | **Vector1 direction** | **Vector2 direction** |
| 4PL0 | Chain A 273-274,285-286<br>Chain B 48-49,64-67 | Chain A 47-52,60-67<br>Chain B 273-275,284-286 | Chain A 24-25,90-109,125-141,309-323<br>Chain B 188-212,229-250 | Chain A 188-213,228-250<br>Chain B 21-25,90-109,124-141,308-322 |
| 5TTP | Chain A 161,167,270-273,284-287<br>Chain B 47-52,63-67 | Chain A 46-52,61-67<br>Chain B 270-272,283-286 | Chain A 91-110,123-138,309-322<br>Chain B 190-211,226-244 | Chain A 190-211,227-245<br>Chain B 90-108,123-139,309-321 |
| 4RY2 | Chain A 412<br>Chain B 192-194,204-206 | Chain A 192-195,203-206<br>Chain B 412,423 | Chain A 232-252,265-282,447-462<br>Chain B 327-351,366-388 | Chain A 327-352,366-388<br>Chain B 232-249,264-282,447-461 |
| 4S0F | | Chain A 194-195,203-206<br>Chain B 412,423 | Chain A 231-252,265-284,447-462<br>Chain B 327-351,366-388 | |
| 5C73 | Chain C 278<br>Chain K 42-43,72 | Chain C 40-44,69-74<br>Chain K 277-278,286-287 | Chain C 16-17,98-118,131-148,311-327<br>Chain K 195-218,232-253 | Chain A 195-218,232-255<br>Chain B 15-17,96-117,131-148,311-324 |
| 2ONJ | Chain A 265-267,279-282<br>Chain B 37-42,58-62 | Chain A 38-43,56-59<br>Chain B 267,278-282 | Chain A 15,83,86-105,120-135,304-317<br>Chain B 182-205,221-244 | Chain A 183-208,221-242<br>Chain B 86-103,119-135,303-316 |
| 4Q4A | Chain A 256-259,271-272<br>Chain B 61-65,76-79 | Chain A 37-42,49-56<br>Chain B 280-283,294-296 | Chain A 78-98,111-129,294-309 | Chain A 174-200,213-235<br>Chain B 100-121,135-155,316-332 |

| | | | | |
|---|---|---|---|---|
| | | | Chain B 199-223,237-258,260-260 | |
| 5MKK | Chain A 276-280,292-293<br>Chain B 47-50,58-61 | Chain A 51-55,67-76<br>Chain B 264-267,276-277 | Chain A 99-116,133-151,314-330<br>Chain B 181-204,221-240,242 | Chain A 197-222,234-254<br>Chain B 22-24,83-103,117-137,297-314 |
| 6RAF | Chain A 277-280,292-293<br>Chain B 46-50,58-61 | Chain A 51-55,67-76<br>Chain B 263-267,276-278 | Chain A 98-116,133-152,314-330<br>Chain B 181-204,221-239 | Chain A 196-222,234-255<br>Chain B 23,84-103,117-136,300-314 |
| 6RAG | Chain A 277-280,292-293<br>Chain B 47-50,58-61 | Chain A 51-55,67-76<br>Chain B 264-267,276-277 | Chain A 98-116,133-152,314-330<br>Chain B 180-204,221-242 | Chain A 196-222,234-255<br>Chain B 22-24,83-103,117-137,297-314 |
| 6RAH | Chain A 279-280,292-295<br>Chain B 46-50,58 | Chain A 48,50-55,67-75<br>Chain B 262-267,276-280 | Chain A 99-116,133-149,315-330<br>Chain B 181-204,221-240 | Chain A 195-222,234-256<br>Chain B 23,84-103,117-134,301-314 |
| 6RAI | Chain A 279-280,292<br>Chain B 46-50,58-60 | Chain A 52-55,67-74<br>Chain B 264-267,276-277 | Chain A 27-29,96-116,133-153,313-330<br>Chain B 178-204,221-243 | Chain A 195-222,234-257<br>Chain B 22-24,81-103,117-137,297-297,299-314 |
| 6RAJ | Chain A 276-280,292-295<br>Chain B 46-50,58-61 | Chain A 48,50-55,67-74<br>Chain B 263-267,276-280 | Chain A 99-116,133-149,316-330<br>Chain B 181-204,221-240 | Chain A 195-222,234-255<br>Chain B 84-103,117-134,301-314 |
| 6RAK | Chain A 279-280,292<br>Chain B 46-50,58-60 | Chain A 52-55,67-75<br>Chain B 265-267,276-277 | Chain A 27-29,96-116,133-152,313-330<br>Chain B 178-204,221-243 | Chain A 195-222,234-257<br>Chain B 22-24,81-103,117-137,299-314 |
| 6RAL | Chain A 276-280,292-294<br>Chain B 46-50,58-61 | Chain A 51-55,67-76<br>Chain B 263-267,276-278 | Chain A 99-116,133-150,315-330<br>Chain B 180-204,221-241 | Chain A 196-222,234-255<br>Chain B 22-23,84-103,117-134,136-136,300-314 |
| 6RAM | Chain A 277-280,292-293<br>Chain B 46-50,58-61 | Chain A 51-55,67-75<br>Chain B 263-267,276-278 | Chain A 27-29,96-116,133-152,313-330<br>Chain B 180-204,221-242 | Chain A 195-222,234-257<br>Chain B 22-23,82-103,117-136,299-314 |
| 6RAN | Chain A 277-280,292-293<br>Chain B 46-50,58-61 | Chain A 51-55,67-76<br>Chain B 262-267,276-278 | Chain A 27-28,98-116,133-152,312-330<br>Chain B 180-204,221-240,242 | Chain A 195-222,234-257<br>Chain B 22-24,82-103,117-137,299-314 |

| | | | | |
|---|---|---|---|---|
| **5UJ9** | Chain A 568-569,581,996-999,1011-1017 | Chain A 348-352,360-363,1217-1218,1226-1228 | Chain A 324-325,386-409,422-439,441,603-619,1130-1158,1172-1195 | Chain A 484-510,523-542,1039-1056,1070-1089,1250-1264 |
| **5UJA** | Chain A 460,565-569,581-582,995-999,1011-1017 | Chain A 348-352,360-363,1226-1227 | Chain A 324-325,386-409,422-439,605-619,1130-1158,1172-1195,1197-1197 | Chain A 487-510,523-540,1039-1056,1070-1087,1089-1089,1248-1248,1250-1264 |
| **6BHU** | Chain A 460,565-569,581-585,995-999,1011-1016 | Chain A 348-352,360-363,1218,1226-1230 | Chain A 324-325,386-409,422-438,605-619,1130-1158,1172-1195 | Chain A 487-510,523-542,971,1037-1056,1070-1087,1251-1264 |
| **6UY0** | Chain A 460,565-569,581-584,996-999,1011-1015 | Chain A 348,350-352,360-363,1218,1226-1230 | Chain A 324-325,386-409,422-438,605-619,1131-1158,1172-1195 | Chain A 487-510,523-542,1038-1056,1070-1087,1251-1264 |
| 6LR0 | Chain U 342-345,358-359,778-781,793-798 | Chain U 84-89,135-141,1001-1006,1015-1017 | Chain U 164-184,197-214,381-396,920-943,960-977 | Chain U 261-286,299-319,820-838,856-875,1037-1053 |
| 6S7P | Chain A 319-321,332-334,733-737,748-752 | Chain A 77-80,109-116,957-960,971-974 | Chain A 52-53,140-158,173-188,358-370,877-900,914-933 | Chain A 236-261,275-294,777-793,812-829,996-1009 |
| 6QEX | Chain A 317-318,330-330,736-737,751-753 | Chain A 74-82,104-114,961-962,971-972 | Chain A 137-153,171-187,353-368,880-900,916-935 | Chain A 235,237-259,272-295,709-712,775-796,813-833,995-1009 |
| 6C0V | Chain A 209-209,314-318,330-334,732-737,751-755 | Chain A 70-78,107-116,957-962,971-976 | Chain A 139-156,171-185,356-368,878-900,916-934 | Chain A 235-259,272-292,779-796,813-830,997-1009 |

**b)** NBD distance and NBD twist angle

| **PDB ID** | **NBD distance** | | **NBD twist angle** | | | |
|---|---|---|---|---|---|---|
| | ***NBD1*** | ***NBD2*** | ***Vector1 origin*** | ***Vector1 direction*** | ***Vector2 origin*** | ***Vector2 direction*** |
| 4PL0 | Chain A 344-554 | Chain B 345-559 | Chain A 344-425,500-506,530-554 | Chain A 432-497,512-527 | Chain B 345-425,500-506,530-559 | Chain B 432-497,512-527 |
| 5TTP | Chain A 341-553 | Chain B 342-556 | Chain A 341-423,500- | Chain A 432-497,512-527 | Chain B 342-423,500-505,530-556 | Chain B 432-497,512-527 |

| | | | | | | |
|---|---|---|---|---|---|---|
| | | | 505,530-553 | | | |
| 4RY2<br>4S0F | Chain A 485-695 | Chain B 486-700 | Chain A 485-566,643-648,673-695 | Chain A 573-640,654-669 | Chain B 486-566,643-647,673-700 | Chain B 573-640,654-669 |
| 5C73 | Chain C 348-558 | Chain K 348-562 | Chain C 348-429,505-511,535-558 | Chain C 434-501,517-532 | Chain K 348-429,505-511,535-562 | Chain K 434-501,517-532 |
| 2ONJ | Chain A 339-550 | Chain B 340-554 | Chain A 339-421,498-503,528-550 | Chain A 427-495,509-525 | Chain B 340-421,498-503,528-554 | Chain B 427-495,509-525 |
| 4Q4A | Chain A 331-541 | Chain B 355-569 | Chain A 331-413,490-495,520-541 | Chain A 419-487,501-515 | Chain B 355-435,512-516,542-569 | Chain B 440-509,523-539 |
| 5MKK<br>6RAF<br>6RAG<br>6RAH<br>6RAI<br>6RAJ<br>6RAK<br>6RAL<br>6RAM<br>6RAN | Chain A 352-569 | Chain B 339-553 | Chain A 352-440,518-523,548-569 | Chain A 447-514,530-545 | Chain B 339-418,495-500,525-553 | Chain B 422-492,507-520 |
| **5UJ9**<br>**5UJA**<br>**6BHU**<br>**6UY0** | Chain A 643-842 | Chain A 1291-1505 | Chain A 643-711,788-791,821-850 | Chain A 722-784,800-810 | Chain A 1290-1372,1449-1453,1479-1507 | Chain A 1381-1445,1461-1474 |
| 6LR0 | Chain U 419-630 | Chain U 1080-1294 | Chain U 419-502,578-584,609-630 | Chain U 506-574,591-605 | Chain U 1080-1160,1239-1243,1269-1294 | Chain U 1166-1235,1252-1263 |
| 6S7P | Chain A 393-605 | Chain A 1034-1248 | Chain A 393-476,553-558,583-612 | Chain A 481-549,565-579 | Chain A 1034-1116,1195-1200,1225-1248 | Chain A 1121-1191,1207-1222 |
| 6QEX | Chain A 391-602 | Chain A 1035-1249 | Chain A 391-474,551-556,581-602 | Chain A 479-547,563-576 | Chain A 1035-1116,1196-1201,1226-1248 | Chain A 1121-1191,1208-1222 |

| | | | | | | |
|---|---|---|---|---|---|---|
| 6C0V | Chain A 392-602 | Chain A 1035-1249 | Chain A 392-473,551-556,581-602 | Chain A 480-547,563-577 | Chain A 1035-1117,1196-1201,1225-1249 | Chain A 1123-1193,1208-1221 |

**Supplementary Table 2. EC angles (°) calculated from each MD simulation replica performed for the different *b*MRP1 systems in different lipid bilayer membranes considered in the present study. Standard deviations are reported in brackets.**

| | | IF apo *b*MRP1 | *b*MRP1-(ATP)$_2$ | *b*MRP1-LTX | *b*MRP1-LTX-(ATP)$_2$ | OF *b*MRP1-(ATP)$_2$ |
|---|---|---|---|---|---|---|
| ***POPC*** | *rep1* | 13.43 (0.26) | 13.71 (0.28) | 14.10 (0.27) | 14.10 (0.27) | 13.93 (0.66) |
| | *rep2* | 13.54 (0.31) | 13.90 (0.36) | 13.54 (0.29) | 13.54 (0.29) | 20.58 (1.57) |
| | *rep3* | 13.37 (0.31) | 13.12 (0.37) | 13.33 (0.29) | 13.33 (0.29) | 16.98 (1.43) |
| ***POPE*** | *rep1* | 13.34 (0.24) | | | | 17.53 (0.89) |
| | *rep2* | 13.65 (0.28) | - | - | - | 15.66 (0.52) |
| | *rep3* | 13.32 (0.24) | | | | 16.74 (1.79) |
| ***POPC:POPE (3:1)*** | *rep1* | 13.31 (0.33) | | | | 17.41 (2.46) |
| | *rep2* | 13.48 (0.40) | - | - | - | 13.32 (0.46) |
| | *rep3* | 13.70 (0.41) | | | | 13.57 (0.34) |
| ***POPC:Chol (3:1)*** | *rep1* | 13.52 (0.27) | 13.44 (0.29) | 13.33 (0.27) | 13.33 (0.27) | 18.87 (2.54) |
| | *rep2* | 13.47 (0.28) | 13.22 (0.24) | 13.48 (0.3) | 13.48 (0.3) | 14.5 (0.79) |
| | *rep3* | 13.69 (0.27) | 13.12 (0.28) | 13.34 (0.26) | 13.34 (0.26) | 14.63 (1.00) |
| ***POPC:POPE:Chol (2:1:1)*** | *rep1* | 13.31 (0.26) | 13.34 (0.25) | 13.28 (0.25) | 13.28 (0.25) | 18.92 (1.42) |
| | *rep2* | 13.55 (0.29) | 13.36 (0.25) | 13.47 (0.24) | 13.47 (0.24) | 18.67 (0.96) |
| | *rep3* | 13.58 (0.34) | 13.46 (0.23) | 13.41 (0.26) | 13.41 (0.26) | 16.28 (0.86) |

**Supplementary Table 3. IC angles (°) calculated from each MD simulation replica performed for the different *b*MRP1 systems in different lipid bilayer membranes considered in the present study. Standard deviations are reported in brackets.**

| | | **IF apo *b*MRP1** | ***b*MRP1-(ATP)$_2$** | ***b*MRP1-LTX** | ***b*MRP1-LTX-(ATP)$_2$** | **OF *b*MRP1-(ATP)$_2$** |
|---|---|---|---|---|---|---|
| ***POPC*** | *rep1* | 31.53 (2.60) | 35.37 (1.77) | 29.92 (2.86) | 29.92 (2.86) | 20.77 (0.31) |
| | *rep2* | 33.22 (2.04) | 33.86 (1.00) | 27.58 (0.55) | 27.58 (0.55) | 20.79 (0.21) |
| | *rep3* | 28.69 (2.56) | 26.89 (1.29) | 26.43 (0.61) | 26.43 (0.61) | 21.06 (0.24) |
| ***POPE*** | *rep1* | 27.77 (0.91) | | | | 20.69 (0.28) |
| | *rep2* | 28.38 (2.49) | - | - | - | 20.56 (0.63) |
| | *rep3* | 28.88 (1.22) | | | | 20.9 (0.42) |
| ***POPC:POPE (3:1)*** | *rep1* | 26.94 (0.96) | | | | 20.64 (0.43) |
| | *rep2* | 27.76 (2.27) | - | - | - | 20.74 (0.29) |
| | *rep3* | 27.85 (2.35) | | | | 21.09 (0.42) |
| ***POPC:Chol (3:1)*** | *rep1* | 28.63 (1.57) | 28.75 (1.70) | 26.04 (0.53) | 26.04 (0.53) | 20.42 (0.25) |
| | *rep2* | 28.36 (1.48) | 27.15 (1.55) | 26.78 (0.61) | 26.78 (0.61) | 20.64 (0.31) |
| | *rep3* | 26.28 (1.62) | 26.45 (1.59) | 26.46 (0.44) | 26.46 (0.44) | 20.82 (0.21) |
| ***POPC:POPE:Chol (2:1:1)*** | *rep1* | 27.69 (1.12) | 27.50 (1.1) | 27.60 (1.03) | 27.60 (1.03) | 21.51 (0.37) |
| | *rep2* | 26.64 (1.66) | 27.34 (1.22) | 26.05 (0.73) | 26.05 (0.73) | 20.92 (0.24) |
| | *rep3* | 31.12 (3.90) | 30.98 (1.02) | 26.25 (0.54) | 26.25 (0.54) | 20.75 (0.34) |

**Supplementary Table 4. NBD distances (Å) calculated from each MD simulation replica performed for the different *b*MRP1 systems in different lipid bilayer membranes considered in the present study. Standard deviations are reported in brackets.**

| | | IF apo *b*MRP1 | *b*MRP1-$(ATP)_2$ | *b*MRP1-LTX | *b*MRP1-LTX-$(ATP)_2$ | OF *b*MRP1-$(ATP)_2$ |
|---|---|---|---|---|---|---|
| ***POPC*** | *rep1* | 39.04 (7.33) | 51.80 (4.80) | 43.52 (9.74) | 43.52 (9.74) | 27.05 (0.21) |
| | *rep2* | 52.62 (6.75) | 55.01 (2.83) | 33.89 (1.04) | 33.89 (1.04) | 27.46 (0.41) |
| | *rep3* | 34.42 (7.64) | 33.25 (3.10) | 32.13 (2.35) | 32.13 (2.35) | 27.25 (0.21) |
| ***POPE*** | *rep1* | 31.92 (3.01) | | | | 27.12 (0.19) |
| | *rep2* | 34.11 (7.18) | - | - | - | 27.29 (0.18) |
| | *rep3* | 34.19 (4.84) | | | | 27.29 (0.21) |
| ***POPC:POPE (3:1)*** | *rep1* | 30.36 (2.63) | | | | 27.27 (0.16) |
| | *rep2* | 32.81 (5.55) | - | - | - | 27.11 (0.18) |
| | *rep3* | 33.64 (6.71) | | | | 27.38 (0.19) |
| ***POPC:Chol (3:1)*** | *rep1* | 31.68 (4.25) | 33.48 (6.34) | 28.93 (1.46) | 28.93 (1.46) | 27.59 (0.25) |
| | *rep2* | 33.30 (4.12) | 33.61 (4.11) | 31.55 (1.86) | 31.55 (1.86) | 27.57 (0.36) |
| | *rep3* | 35.82 (3.99) | 32.05 (4.99) | 31.09 (0.57) | 31.09 (0.57) | 27.88 (0.27) |
| ***POPC:POPE:Chol (2:1:1)*** | *rep1* | 32.68 (4.48) | 31.33 (2.74) | 34.52 (3.36) | 34.52 (3.36) | 27.33 (0.21) |
| | *rep2* | 34.91 (5.51) | 31.48 (4.49) | 30.25 (1.92) | 30.25 (1.92) | 27.36 (0.22) |
| | *rep3* | 40.33 (10.09) | 36.97 (1.52) | 30.56 (1.6) | 30.56 (1.60) | 27.06 (0.17) |

**Supplementary Table 5. NBD twists (°) calculated from each MD simulation replica performed for the different *b*MRP1 systems in different lipid bilayer membranes considered in the present study. Standard deviations are reported in brackets.**

| | | **IF apo *b*MRP1** | ***b*MRP1-(ATP)₂** | ***b*MRP1-LTX** | ***b*MRP1-LTX-(ATP)₂** | **OF *b*MRP1-(ATP)₂** |
|---|---|---|---|---|---|---|
| ***POPC*** | *rep1* | -141.82 (5.55) | -156.78 (26.72) | -151.58 (8.55) | -151.58 (8.55) | -143.28 (1.88) |
| | *rep2* | -140.39 (8.67) | -150.38 (44.18) | -126.67 (6.09) | -126.67 (6.09) | -143.66 (2.26) |
| | *rep3* | -147.39 (5.87) | -145.18 (32.55) | -140.15 (4.47) | -140.15 (4.47) | -144.72 (1.93) |
| ***POPE*** | *rep1* | -126.04 (5.12) | - | - | - | -143.67 (1.65) |
| | *rep2* | -136.34 (7.63) | | | | -143.58 (1.39) |
| | *rep3* | -143.84 (5.92) | | | | -143.15 (1.80) |
| ***POPC:POPE (3:1)*** | *rep1* | -139.79 (4.03) | - | - | - | -140.56 (1.73) |
| | *rep2* | -143.57 (7.43) | | | | -142.32 (2.01) |
| | *rep3* | -146.51 (11.38) | | | | -143.98 (1.75) |
| ***POPC:Chol (3:1)*** | *rep1* | -148.02 (12.69) | -148.89 (17.76) | -131.54 (4.38) | -131.54 (4.38) | -142.01 (1.80) |
| | *rep2* | -155.97 (8.17) | -142.70 (6.88) | -137.66 (4.27) | -137.66 (4.27) | -144.01 (2.29) |
| | *rep3* | -136.86 (62.01) | -142.80 (5.18) | -139.52 (4.81) | -139.52 (4.81) | -147.90 (2.41) |
| ***POPC:POPE:Chol (2:1:1)*** | *rep1* | -146.03 (9.40) | -136.68 (5.57) | -137.92 (5.62) | -137.92 (5.62) | -141.05 (1.80) |
| | *rep2* | -150.91 (43.34) | -140.95 (9.11) | -134.44 (6.66) | -134.44 (6.66) | -146.40 (1.99) |
| | *rep3* | -146.99 (8.90) | -143.58 (4.61) | -131.17 (3.77) | -131.17 (3.77) | -145.81 (1.79) |

**Supplementary Table 6. Overlaps of calculated network communities between the different IF systems and OF *b*MRP1-(ATP)$_2$. Analyses were performed in POPC:POPE:Chol (2:1:1) lipid bilayer.** For a given community, overlaps were obtained by calculating the proportion of shared residues between target system and OF *b*MRP1-(ATP)$_2$.

| Community name | **IF apo *b*MRP1** | ***b*MRP1-(ATP)$_2$** | ***b*MRP1-LTX** | ***b*MRP1-LTX-(ATP)$_2$** |
|---|---|---|---|---|
| **NBD1 Walker A** | 81.94% | 87.39% | 88.37% | 87.79% |
| **NBD1 signature** | 82.35% | 83.08% | 83.33% | 84.85 % |
| **NBD2 Walker A** | 71.32% | 75.70% | 72.32% | 69.66% |
| **NBD2 signature** | 71.64% | 73.24% | 64.63% | 71.01% |

**Supplementary Table 7. The number of calculated raw NBD communities obtained from MD simulations performed in POPC:POPE:Chol (2:1:1).** It is worth mentioning that for IF apo *b*MRP1 and OF *bMRP1*-$(ATP)_2$, more than 2 communities per NBD were sometimes observed. They were however systematically corresponding to subcommunities of either Walker A or signature community described in Supplementary Table 6.

| | **IF apo *b*MRP1** | | ***b*MRP1-$(ATP)_2$** | | ***b*MRP1-LTX** | | ***b*MRP1-LTX-$(ATP)_2$** | | **OF *b*MRP1-$(ATP)_2$** | |
|---|---|---|---|---|---|---|---|---|---|---|
| | **NBD1** | **NBD2** | **NBD1** | **NBD2** | **NBD1** | **NBD2** | **NBD1** | **NBD2** | **NBD1** | **NBD2** |
| **Rep1** | **4** | **3** | 2 | 2 | 2 | 2 | 2 | **3** | 2 | **3** |
| **Rep2** | 2 | 2 | 2 | 2 | 2 | 2 | 2 | 2 | 2 | **4** |
| **Rep3** | **3** | **3** | 2 | 2 | 2 | 2 | 2 | 2 | 2 | 2 |

**Supplementary Table 8. Key distances between ATP and *b*MRP1 NBD or IC-loop for a) OF *b*MRP1-(ATP)$_2$, b) *b*MRP1-LTX-(ATP)$_2$ and c) *b*MRP1-(ATP)$_2$.** Values were averaged over each replica; standard deviations are reported in brackets.

**a)** OF *b*MRP1-(ATP)$_2$

| Membrane | Replica | ATP1 | | | | | | | | | | |
|---|---|---|---|---|---|---|---|---|---|---|---|---|
| | | | | | | | PG | | | | W653 (A-loop) | 6 amino group |
| | | W653 (A-loop) | K684 (Walker A) | S685 (Walker A) | W653 (A-loop) | S1430 (Signature) | D793 (Walker B) | G770 (Signature) | Q713 (Q-loop) | H827 (H-loop) | G770 Signature) | N412 |
| POPC | 1 | 4.48 (0.35) | 4.34 (0.09) | 5.03 (0.07) | 7.06 (0.09) | 4.72 (0.15) | 9.33 (0.32) | 17.29 (0.36) | 7.44 (0.24) | 8.61 (0.41) | 20.88 (0.34) | 5.41 (0.74) |
| | 2 | 4.38 (0.31) | 4.38 (0.13) | 5.02 (0.07) | 7.05 (0.09) | 4.72 (0.14) | 9.32 (0.43) | 17.23 (0.54) | 7.36 (0.31) | 8.8 (0.4) | 20.71 (0.63) | 5.4 (0.77) |
| | 3 | 4.99 (0.52) | 4.35 (0.08) | 5.04 (0.07) | 7.03 (0.09) | 5.57 (0.52) | 9.09 (0.37) | 16.86 (0.38) | 7.52 (0.28) | 8.64 (0.43) | 19.76 (0.53) | 6.2 (0.91) |
| POPC:Chol (3:1) | 1 | 4.68 (0.45) | 4.42 (0.11) | 5.03 (0.07) | 7.08 (0.1) | 4.7 (0.13) | 9.27 (0.35) | 16.46 (0.41) | 6.76 (0.16) | 8.94 (0.39) | 19.76 (0.48) | 6.35 (1.2) |
| | 2 | 5.16 (0.3) | 4.37 (0.08) | 5.03 (0.07) | 7.02 (0.09) | 4.86 (0.31) | 9.14 (0.37) | 15.86 (0.59) | 7.21 (0.19) | 8.66 (0.51) | 18.6 (0.73) | 6.59 (0.64) |
| | 3 | 5.38 (0.42) | 4.38 (0.08) | 5.07 (0.07) | 7.07 (0.1) | 5.71 (0.22) | 8.58 (0.37) | 16.1 (0.73) | 7.09 (0.19) | 9.62 (0.66) | 18.24 (0.6) | 7.71 (0.93) |
| POPC:POPE: Chol (2:1:1) | 1 | 4.59 (0.35) | 4.36 (0.08) | 5.02 (0.07) | 7.04 (0.09) | 4.71 (0.12) | 11.43 (1.12) | 17.43 (0.49) | 6.68 (0.22) | 8.92 (0.39) | 20.93 (0.44) | 5.02 (0.64) |
| | 2 | 4.72 (0.57) | 4.41 (0.11) | 5.02 (0.07) | 7.05 (0.1) | 4.74 (0.2) | 9.3 (0.33) | 17.61 (0.48) | 6.79 (0.17) | 8.92 (0.41) | 20.74 (0.67) | 5.68 (1.36) |
| | 3 | 4.64 (0.37) | 4.44 (0.15) | 5.02 (0.07) | 7.06 (0.1) | 4.68 (0.12) | 9.17 (0.39) | 17.64 (0.44) | 6.94 (0.26) | 8.85 (0.42) | 21.12 (0.36) | 7.35 (1.34) |

| Membrane | Replica | ATP2 | | | | | | | | | | |
|---|---|---|---|---|---|---|---|---|---|---|---|---|
| | | | | | | | PG | | | | Ribose ring | 6 amino group |
| | | Y1301 (A-loop) | K1332 (Walker A) | S1333 (Walker A) | S1334 (Walker A) | G771 (Signature) | E1454 (Walker B) | V1431 (Signature) | Q1374 (Q-loop) | H1485 (H-loop)) | V1431 (Signature) | E1064 |
| POPC | 1 | 4.62 (0.35) | 4.49 (0.09) | 5.07 (0.07) | 7.09 (0.1) | 6.94 (0.17) | 8.3 (0.31) | 16.55 (0.29) | 6.25 (0.15) | 8.19 (0.68) | 20.09 (0.29) | 6.1 (0.79) |
| | 2 | 4.67 (0.74) | 4.49 (0.1) | 5.09 (0.08) | 7.1 (0.1) | 7.72 (1.28) | 8.29 (0.76) | 17.03 (0.73) | 6.49 (0.37) | 8.88 (0.88) | 20.34 (0.41) | 6.3 (1.29) |
| | 3 | 4.45 (0.33) | 4.55 (0.11) | 5.06 (0.07) | 7.07 (0.1) | 6.97 (0.18) | 8.2 (0.57) | 14.05 (1.27) | 6.34 (0.16) | 9.26 (1.19) | 17.55 (1.4) | 5.65 (0.74) |

| | | | | | | | | | | | | |
|---|---|---|---|---|---|---|---|---|---|---|---|---|
| **POPC:Chol (3:1)** | **1** | 4.36 (0.33) | 4.62 (0.18) | 5.06 (0.09) | 7.01 (0.12) | 8.26 (0.46) | 6.8 (0.22) | 17.61 (0.39) | 7.29 (0.2) | 9.6 (1.19) | 20.1 (0.35) | 5.53 (1.0) |
| | **2** | 4.38 (0.3) | 4.65 (0.22) | 5.07 (0.09) | 7.02 (0.12) | 8.81 (0.75) | 6.55 (0.22) | 15.68 (0.7) | 7.08 (0.27) | 10.28 (0.64) | 18.24 (0.61) | 4.92 (0.8) |
| | **3** | 5.07 (0.84) | 4.8 (0.16) | 5.1 (0.08) | 7.01 (0.11) | 9.52 (0.66) | 6.62 (0.17) | 14.22 (0.81) | 7.03 (0.26) | 7.82 (0.65) | 16.6 (0.86) | 6.48 (1.88) |
| **POPC:POPE: Chol (2:1:1)** | **1** | 4.56 (0.38) | 4.75 (0.15) | 5.04 (0.07) | 7.12 (0.12) | 6.89 (0.23) | 7.92 (0.46) | 16.34 (0.49) | 6.22 (0.13) | 10.33 (1.27) | 20.08 (0.49) | 5.18 (0.55) |
| | **2** | 4.51 (0.3) | 4.51 (0.12) | 5.03 (0.07) | 7.1 (0.11) | 7.05 (0.17) | 9.23 (0.42) | 16.5 (0.61) | 6.16 (0.16) | 10.0 (0.83) | 20.24 (0.6) | 4.99 (0.44) |
| | **3** | 4.64 (0.36) | 4.62 (0.15) | 5.01 (0.07) | 7.09 (0.1) | 6.9 (0.18) | 8.23 (0.98) | 16.66 (0.37) | 6.13 (0.11) | 10.74 (0.74) | 20.37 (0.37) | 6.45 (0.8) |

**b)** *b*MRP1-LTX-(ATP)$_2$

| **Membrane** | **Replica** | **ATP1** | | | | | | | | | | |
|---|---|---|---|---|---|---|---|---|---|---|---|---|
| | | | | | | | **PG** | | | | **Ribose ring** | **6 amino group** |
| | | **W653 (A-loop)** | **K684 (Walker A)** | **S685 (Walker A)** | **S686 (Walker A)** | **S1430 (Signature)** | **D793 (Walker B)** | **G770 (Signature)** | **Q713 (Q-loop)** | **H827** | **G770** | **N412** |
| **POPC** | **1** | 4.94 (0.52) | 4.64 (0.36) | 5.12 (0.11) | 7.07 (0.12) | 8.33 (2.9) | 9.63 (0.63) | 16.58 (0.89) | 7.54 (1.02) | 10.57 (1.47) | 19.13 (0.68) | 5.56 (1.3) |
| | **2** | 5.97 (1.26) | 4.41 (0.1) | 5.11 (0.08) | 7.09 (0.1) | 18.46 (8.73) | 9.08 (0.54) | 16.04 (0.7) | 6.92 (0.21) | 9.4 (1.05) | 18.06 (0.88) | 6.68 (1.79) |
| | **3** | 5.43 (0.7) | 4.49 (0.11) | 5.17 (0.09) | 6.94 (0.1) | 7.29 (4.33) | 12.0 (0.48) | 16.21 (0.67) | 7.01 (0.31) | 8.96 (0.94) | 18.8 (0.77) | 8.64 (1.91) |
| **POPC:Chol (3:1)** | **1** | 9.14 (3.24) | 4.42 (0.11) | 5.08 (0.08) | 7.07 (0.12) | 10.23 (4.32) | 9.24 (0.55) | 16.07 (0.64) | 7.06 (0.27) | 9.75 (1.0) | 18.78 (0.9) | 12.84 (7.44) |
| | **2** | 5.47 (1.56) | 4.46 (0.14) | 5.12 (0.08) | 7.06 (0.1) | 20.6 (1.74) | 8.6 (0.45) | 15.67 (0.61) | 7.15 (0.28) | 8.64 (0.65) | 17.16 (0.8) | 7.22 (3.78) |
| | **3** | 6.76 (2.73) | 4.43 (0.12) | 5.09 (0.09) | 7.09 (0.15) | 17.95 (3.19) | 8.97 (0.54) | 16.32 (0.74) | 7.02 (0.26) | 9.85 (0.92) | 18.62 (0.96) | 9.35 (5.62) |
| **POPC:POPE:C hol (2:1:1)** | **1** | 6.81 (2.4) | 4.46 (0.16) | 5.11 (0.1) | 7.08 (0.11) | 9.89 (2.03) | 8.71 (0.56) | 15.6 (0.71) | 7.02 (0.24) | 8.74 (0.89) | 17.48 (0.96) | 9.08 (5.41) |
| | **2** | 5.92 (1.61) | 4.41 (0.11) | 5.1 (0.08) | 7.07 (0.1) | 9.58 (1.35) | 8.76 (0.62) | 16.63 (0.95) | 7.03 (0.27) | 9.58 (1.17) | 18.38 (1.28) | 8.45 (2.75) |
| | **3** | 7.12 (2.42) | 4.41 (0.13) | 5.1 (0.08) | 7.11 (0.16) | 10.25 (1.53) | 9.04 (0.61) | 16.99 (0.62) | 6.87 (0.24) | 9.41 (1.27) | 18.5 (0.94) | 7.56 (3.06) |

| Membrane | Replica | ATP2 | | | | | | | | | | |
|---|---|---|---|---|---|---|---|---|---|---|---|---|
| | | | | | | | PG | | | | Ribose ring | 6 amino group |
| | | Y1301 (A-loop) | K1332 (Walker A) | S1333 (Walker A) | S1334 (Walker A) | G771 (Signature) | E1454 (Walker B) | V1431 (Signature) | Q1374 (Q-loop) | H1485 | V1431 | E1064 |
| POPC | 1 | 9.39 (0.61) | 4.53 (0.09) | 5.15 (0.09) | 6.91 (0.1) | 11.04 (2.19) | 6.4 (0.31) | 16.97 (0.49) | 6.75 (0.25) | 6.72 (0.62) | 19.92 (0.95) | 11.91 (1.25) |
| | 2 | 10.01 (1.31) | 4.53 (0.11) | 5.15 (0.09) | 6.92 (0.09) | 23.93 (8.47) | 6.02 (0.43) | 17.05 (0.84) | 6.92 (0.28) | 9.45 (1.61) | 19.45 (0.96) | 7.63 (2.85) |
| | 3 | 12.68 (1.49) | 4.53 (0.09) | 5.09 (0.09) | 6.92 (0.1) | 13.67 (2.73) | 6.49 (0.32) | 17.53 (0.94) | 6.46 (0.29) | 6.4 (0.53) | 20.62 (1.04) | 11.21 (2.63) |
| POPC:Chol (3:1) | 1 | 10.79 (1.85) | 4.53 (0.1) | 5.17 (0.09) | 6.94 (0.1) | 14.25 (4.12) | 7.17 (0.53) | 14.83 (1.06) | 6.84 (0.29) | 11.27 (0.93) | 17.51 (1.27) | 13.76 (6.93) |
| | 2 | 10.46 (2.99) | 4.48 (0.11) | 5.11 (0.08) | 7.07 (0.1) | 22.66 (1.79) | 6.56 (0.67) | 15.38 (1.35) | 6.7 (0.2) | 8.51 (0.99) | 17.84 (1.47) | 7.11 (1.8) |
| | 3 | 10.85 (0.98) | 4.48 (0.11) | 5.08 (0.09) | 7.06 (0.12) | 13.91 (1.17) | 6.87 (0.54) | 16.7 (0.78) | 6.74 (0.26) | 9.53 (1.33) | 19.78 (0.87) | 9.97 (2.61) |
| POPC:POPE:Chol (2:1:1) | 1 | 8.59 (0.85) | 4.51 (0.09) | 5.15 (0.09) | 6.94 (0.1) | 13.93 (1.63) | 12.35 (1.11) | 17.02 (1.01) | 6.7 (0.31) | 9.8 (0.98) | 20.24 (1.23) | 5.36 (1.12) |
| | 2 | 8.4 (1.3) | 4.63 (0.16) | 5.14 (0.09) | 6.93 (0.11) | 13.27 (1.29) | 6.85 (0.36) | 14.12 (1.04) | 6.63 (0.24) | 12.58 (1.17) | 17.52 (1.51) | 13.45 (5.28) |
| | 3 | 6.19 (2.0) | 4.48 (0.12) | 5.11 (0.09) | 7.06 (0.1) | 15.85 (1.71) | 7.65 (0.55) | 17.13 (0.6) | 6.97 (0.25) | 10.55 (0.54) | 19.25 (0.8) | 10.96 (6.62) |

**c)** *b*MRP1-$(ATP)_2$

| Membrane | Replica | ATP1 | | | | | | | | | | |
|---|---|---|---|---|---|---|---|---|---|---|---|---|
| | | | | | | | PG | | | | Ribose ring | 6 amino group |
| | | W653 (A-loop) | K684 (Walker A) | S685 (Walker A) | S686 (Walker A) | S1430 (Signature) | D793 (Walker B) | G770 (Signature) | Q713 (Q-loop) | H827 | G770 | N412 |
| POPC | 1 | 5.32 (0.47) | 4.48 (0.1) | 5.08 (0.09) | 6.9 (0.09) | 33.15 (5.18) | 12.42 (0.6) | 16.06 (0.53) | 6.08 (0.17) | 9.18 (1.15) | 19.45 (0.59) | 4.82 (1.04) |
| | 2 | 6.27 (2.03) | 4.87 (0.23) | 5.09 (0.1) | 7.07 (0.12) | 34.74 (3.66) | 9.52 (0.8) | 15.82 (0.65) | 6.24 (0.13) | 10.99 (1.17) | 19.15 (0.83) | 9.21 (4.7) |
| | 3 | 5.07 (0.73) | 4.5 (0.14) | 5.09 (0.08) | 6.89 (0.09) | 15.59 (3.87) | 12.2 (0.62) | 16.01 (0.51) | 6.13 (0.18) | 9.09 (0.86) | 19.4 (0.61) | 5.85 (1.62) |

| | | | | | | | | | | | | |
|---|---|---|---|---|---|---|---|---|---|---|---|---|
| POPC:Chol (3:1) | 1 | 5.15 (0.45) | 4.49 (0.14) | 5.08 (0.08) | 6.89 (0.09) | 12.38 (5.73) | 12.01 (0.74) | 15.95 (0.54) | 6.07 (0.18) | 8.98 (0.73) | 19.49 (0.6) | 6.07 (1.35) |
| | 2 | 5.37 (0.52) | 4.46 (0.1) | 5.09 (0.08) | 6.91 (0.09) | 12.23 (3.96) | 12.39 (0.5) | 16.21 (0.52) | 6.04 (0.17) | 9.51 (1.01) | 19.6 (0.62) | 5.26 (1.01) |
| | 3 | 5.47 (0.57) | 4.47 (0.09) | 5.09 (0.08) | 6.91 (0.09) | 12.65 (5.29) | 12.17 (0.49) | 15.98 (0.51) | 6.11 (0.23) | 9.4 (0.92) | 19.26 (0.6) | 5.76 (1.25) |
| POPC:POPE:Chol (2:1:1) | 1 | 5.17 (0.5) | 4.53 (0.16) | 5.09 (0.08) | 6.89 (0.09) | 10.93 (3.2) | 11.89 (0.6) | 15.75 (0.61) | 6.18 (0.21) | 8.68 (1.17) | 19.05 (0.62) | 5.51 (1.0) |
| | 2 | 5.6 (1.16) | 6.44 (1.42) | 5.59 (0.45) | 7.15 (0.5) | 11.18 (4.85) | 10.95 (0.85) | 15.37 (0.86) | 6.23 (0.17) | 13.21 (1.25) | 18.53 (0.96) | 8.12 (3.14) |
| | 3 | 5.1 (0.5) | 4.52 (0.17) | 5.1 (0.09) | 6.9 (0.1) | 23.86 (2.34) | 12.32 (0.83) | 16.04 (0.8) | 6.13 (0.19) | 10.01 (1.53) | 19.52 (0.81) | 5.73 (1.19) |

| Membrane | Replica | ATP2 | | | | | | | | | | |
|---|---|---|---|---|---|---|---|---|---|---|---|---|
| | | | | | | | PG | | | | Ribose ring | 6 amino group |
| | | Y1301 (A-loop) | K1332 (Walker A) | S1333 (Walker A) | S1334 (Walker A) | G771 (Signature) | E1454 (Walker B) | V1431 (Signature) | Q1374 (Q-loop) | H1485 | V1431 | E1064 |
| POPC | 1 | 9.46 (1.03) | 4.52 (0.1) | 5.12 (0.09) | 6.94 (0.1) | 31.35 (5.06) | 8.57 (0.63) | 16.49 (0.69) | 6.1 (0.17) | 11.12 (1.33) | 19.54 (0.77) | 6.63 (1.56) |
| | 2 | 9.42 (1.19) | 4.56 (0.12) | 5.12 (0.09) | 6.91 (0.1) | 37.96 (3.45) | 10.08 (0.56) | 15.25 (0.71) | 6.14 (0.15) | 12.31 (0.99) | 18.91 (0.71) | 9.45 (1.9) |
| | 3 | 12.15 (3.24) | 4.55 (0.17) | 5.13 (0.09) | 7.04 (0.13) | 17.68 (3.05) | 7.66 (1.24) | 15.79 (1.3) | 6.81 (0.2) | 8.76 (1.21) | 17.37 (1.2) | 11.61 (4.82) |
| POPC:Chol (3:1) | 1 | 9.52 (1.85) | 4.47 (0.12) | 5.15 (0.09) | 7.06 (0.1) | 18.29 (5.85) | 8.28 (0.64) | 17.0 (0.89) | 7.02 (0.29) | 8.27 (1.12) | 19.08 (1.3) | 13.52 (4.88) |
| | 2 | 8.77 (2.21) | 4.47 (0.11) | 5.11 (0.08) | 7.08 (0.12) | 18.74 (3.46) | 6.59 (0.56) | 16.5 (1.15) | 6.8 (0.18) | 8.24 (1.92) | 19.76 (1.47) | 12.59 (5.28) |
| | 3 | 9.25 (0.92) | 4.52 (0.09) | 5.16 (0.09) | 6.95 (0.1) | 15.32 (4.64) | 7.5 (1.09) | 15.94 (0.51) | 6.68 (0.27) | 7.81 (1.33) | 18.85 (0.62) | 6.2 (1.2) |
| POPC:POPE:Chol (2:1:1) | 1 | 8.98 (1.11) | 4.54 (0.14) | 5.15 (0.1) | 6.94 (0.1) | 14.11 (2.12) | 8.49 (1.05) | 15.89 (1.27) | 6.56 (0.28) | 9.46 (1.0) | 19.0 (1.21) | 5.63 (1.43) |
| | 2 | 8.55 (1.61) | 4.51 (0.09) | 5.14 (0.09) | 6.94 (0.1) | 15.02 (3.44) | 9.88 (1.05) | 16.66 (0.62) | 5.94 (0.15) | 11.18 (0.95) | 19.77 (0.73) | 6.29 (1.1) |
| | 3 | 12.25 (1.67) | 4.49 (0.15) | 5.07 (0.1) | 7.05 (0.12) | 14.68 (2.04) | 6.46 (0.57) | 17.22 (0.52) | 6.63 (0.18) | 6.78 (0.67) | 20.39 (0.59) | 11.24 (0.76) |

**Supplementary Table 9. Selected residues for allosteric pathway calculations from substrate-binding site to NBS1 and NBS2.**

| Sink | Sources | |
|---|---|---|
| ***LTX-binding site*** | ***NBS1*** | ***NBS2*** |
| K332<br>Y440<br>R593<br>R1196<br>E1203<br>T1241<br>N1244<br>W1245<br>R1248 | Walker A$^{NBD1}$:<br>G681, G683, K684, S686 | Walker A$^{NBD2}$:<br>R1328, T1329, A1331, G1332, S1334 |
| | Q-loop$^{NBD1}$:<br>Q713 | Q-loop$^{NBD2}$:<br>Q1374 |
| | A-loop$^{NBD1}$:<br>W653 | A-loop$^{NBD2}$:<br>Y1301 |
| | ABC Signature$^{NBD2}$:<br>S1430, V1431, G1432, Q1433 | ABC Signature$^{NBD1}$:<br>S769, G770, G771, Q772 |

**Supplementary Table 10. Membrane free energy deformations. Bilayer Thicknesses at the outer edge of the simulation cells in POPC:POPE:Chol (2:1:1).**

| | | **IF apo *b*MRP1** | ***b*MRP1-$(ATP)_2$** | ***b*MRP1-LTX** | ***b*MRP1-LTX-$(ATP)_2$** | **OF *b*MRP1-$(ATP)_2$** |
|---|---|---|---|---|---|---|
| $d_0$ (Å) | *rep1*<br>*rep2*<br>*rep3* | 42.4<br>42.0<br>42.2 | 44.6<br>44.0<br>44.3 | 45.0<br>44.6<br>44.6 | 44.8<br>44.5<br>44.7 | 44.6<br>44.6<br>44.5 |
| Free energy cost of deformation due to membrane protein (kT) | *rep1*<br>*rep2*<br>*rep3* | 60.2<br>54.3<br>66.3 | 42.9<br>53.4<br>46.6 | 47.7<br>56.3<br>49.3 | 43.7<br>50.6<br>47.8 | 42.5<br>48.9<br>46.7 |
| Reference free energy cost for flat membrane (kT) | *rep1*<br>*rep2*<br>*rep3* | 65.9<br>65.9<br>67.7 | 58.6<br>59.1<br>59.7 | 60.5<br>62.9<br>61.7 | 61.1<br>59.5<br>60.6 | 58.6<br>59.1<br>59.7 |
| Number of iterations | *rep1*<br>*rep2*<br>*rep3* | 150<br>150<br>150 | 150<br>150<br>150 | 150<br>150<br>150 | 130<br>130<br>150 | 150<br>150<br>150 |
| Assumed ∆G (Deformation - flat) (kT) | *rep1*<br>*rep2*<br>*rep3* | -5.7<br>11.6<br>-1.4 | -15.7<br>-5.7<br>-13.1 | -14.0<br>-5.4<br>-12.4 | -17.4<br>-8.9<br>-12.8 | -16.1<br>-10.2<br>-13.0 |

* compared to average flat energy cost, since calculations has crashed for flat membrane.

**Supplementary Table 11. Membrane free energy deformations. Bilayer Thicknesses at the outer edge of the simulation cells in POPC:Chol (3:1).**

| | | **IF apo *b*MRP1** | ***b*MRP1-(ATP)$_2$** | ***b*MRP1-LTX** | ***b*MRP1-LTX-(ATP)$_2$** | **OF *b*MRP1-(ATP)$_2$** |
|---|---|---|---|---|---|---|
| $d_0$ (Å) | *rep1*<br>*rep2*<br>*rep3* | 40.0<br>40.7<br>40.7 | 42.7<br>43.4<br>43.3 | 42.8<br>43.1<br>43.8 | 42.9<br>43.8<br>43.2 | 42.9<br>41.8<br>42.3 |
| Free energy cost of deformation due to membrane protein (kT) | *rep1*<br>*rep2*<br>*rep3* | 54.9<br>43.3<br>56.3 | 32.8<br>32.1<br>30.7 | 26.2<br>25.7<br>33.7 | 35.5<br>31.5<br>30.5 | 30.1<br>23.6<br>32.1 |
| Reference free energy cost for flat membrane (kT) | *rep1*<br>*rep2*<br>*rep3* | 30.2<br>31.4<br>29.9 | 28.6<br>28.4<br>28.6 | 26.7<br>26.9<br>25.8 | 27.6<br>26.1<br>26.0 | 29.3<br>27.7<br>30.4 |
| Number of iterations | *rep1*<br>*rep2*<br>*rep3* | 150<br>150<br>150 | 150<br>150<br>150 | 150<br>150<br>150 | 150<br>150<br>150 | 150<br>150<br>150 |
| Assumed ΔG (Deformation - flat) (kT) | *rep1*<br>*rep2*<br>*rep3* | 14.7<br>10.9<br>16.4 | 4.2<br>3.7<br>2.1 | -0.5<br>-1.2<br>7.9 | 7.9<br>5.3<br>4.5 | 0.8<br>-4.1<br>1.7 |

**Supplementary Table 12. Membrane free energy deformations. Bilayer Thicknesses at the outer edge of the simulation cells in pure POPC.**

| | | IF apo *b*MRP1 | *b*MRP1-(ATP)$_2$ | *b*MRP1-LTX | *b*MRP1-LTX-(ATP)$_2$ | OF *b*MRP1-(ATP)$_2$ |
|---|---|---|---|---|---|---|
| $d_0$ (Å) | *rep1* | 36.6 | 38.0 | 38.5 | 38.7 | 35.1 |
| | *rep2* | 36.7 | 38.4 | 37.7 | 38.2 | 33.4 |
| | *rep3* | 36.8 | 38.3 | 38.9 | 37.9 | 35.8 |
| Free energy cost of deformation due to membrane protein (kT) | *rep1* | 15.2 | 20.9 | 12.3 | 11.9 | 18.6 |
| | *rep2* | 19.9 | 16.7 | 16.7 | 12.2 | 22.5 |
| | *rep3* | 18.0 | - | 9.9 | 14.4 | 18.0 |
| Reference free energy cost for flat membrane (kT) | *rep1* | 1.0 | 1.0 | 0.8 | 0.9 | 1.0 |
| | *rep2* | 0.9 | 0.9 | 0.9 | 0.9 | 1.1 |
| | *rep3* | 1.0 | - | 0.9 | 0.9 | 1.1 |
| Number of iterations | *rep1* | 150 | 150 | 150 | 150 | 150 |
| | *rep2* | 150 | 150 | 150 | 150 | 150 |
| | *rep3* | 150 | - | 150 | 150 | 150 |
| Assumed ΔG (Deformation - flat) (kT) | *rep1* | 14.2 | 19.9 | 11.5 | 11.0 | 17.6 |
| | *rep2* | 19.0 | 16.8 | 16.8 | 11.3 | 21.4 |
| | *rep3* | 17.0 | - | 9.0 | 13.5 | 16.9 |

**Supplementary Table 13. $L_0$ modelling of OF state.**

| | |
|---|---|
| 269-RKQPVKIV-276 | $L_0$ of IF system |
| 277-YSSKDPAKPKGSSKVDV-293 | sequence |
| 294-NEEAEALIVKCPQKERD-310 | $L_0$ of IF system |

**Supplementary Table 14. Box sizes (Å) of each system considered in the present study.**

| | IF apo *b*MRP1 | *b*MRP1-$(ATP)_2$ | *b*MRP1-LTX | *b*MRP1-LTX-$(ATP)_2$ | OF *b*MRP1-$(ATP)_2$ |
|---|---|---|---|---|---|
| ***POPC*** | 121.5x121.2 x179.8 | 121.6x121.2 x179.8 | 121.6x121.5 x179.2 | 121.6x121.5 x179.2 | 121.5x121.6 x179.5 |
| ***POPE*** | 119.6x119.6 x178.3 | - | - | - | 121.5x121.6 x179.5 |
| ***POPC:POPE (3:1)*** | 121.5x121.6 x179.7 | - | - | - | 121.5x121.5 x179.5 |
| ***POPC:Chol (3:1)*** | 123.0x123.7 x179.7 | 123.0x123.7 x179.8 | 121.9x122.8 x179.2 | 121.9x122.8 x179.2 | 123.1x126.3 x179.5 |
| ***POPC:POPE:Chol (2:1:1)*** | 121.2x122.4 x179.7 | 121.2x122.4 x179.8 | 122.8x122.3 x179.2 | 122.8x122.3 x179.2 | 123.1x126.3 x179.5 |

**Supplementary Table 15. Number of lipids of the different lipid bilayers investigated in the present study.**

| | *Leaflet* | IF apo *b*MRP1 | *b*MRP1-(ATP)$_2$ | *b*MRP1-LTX | *b*MRP1-LTX-(ATP)$_2$ | OF *b*MRP1-(ATP)$_2$ |
|---|---|---|---|---|---|---|
| ***POPC*** | *upper* | 190 | 190 | 191 | 191 | 189 |
| | *lower* | 174 | 174 | 171 | 171 | 174 |
| ***POPE*** | *upper* | 220 | - | - | - | 220 |
| | *lower* | 200 | | | | 202 |
| ***POPC:Chol (3:1)*** | *upper* | 165 | 165 | 160 | 160 | 159 |
| | *lower* | 151 | 151 | 144 | 144 | 146 |
| | *upper* | 55 | 55 | 54 | 54 | 53 |
| | *lower* | 50 | 50 | 48 | 48 | 49 |
| ***POPC:POPE (3:1)*** | *upper* | 149 | - | - | - | 147 |
| | *lower* | 135 | | | | 135 |
| | *upper* | 49 | | | | 49 |
| | *lower* | 45 | | | | 45 |
| ***POPC:POPE:Chol (2:1:1)*** | *upper* | 110 | 110 | 112 | 112 | 110 |
| | *lower* | 100 | 100 | 100 | 100 | 102 |
| | *upper* | 55 | 55 | 56 | 56 | 55 |
| | *lower* | 50 | 50 | 50 | 50 | 50 |
| | *upper* | 55 | 55 | 56 | 56 | 55 |
| | *lower* | 50 | 49 | 50 | 50 | 50 |

**Supplementary Table 16. Number of atoms for each system investigated in the present study.**

| | IF apo *b*MRP1 | *b*MRP1-(ATP)$_2$ | *b*MRP1-LTX | *b*MRP1-LTX-(ATP)$_2$ | OF *b*MRP1-(ATP)$_2$ |
|---|---|---|---|---|---|
| ***POPC*** | 240 956 | 241 044 | 241 408 | 241 492 | 242 975 |
| ***POPE*** | 244 324 | - | - | - | 247 676 |
| ***POPC:POPE (3:1)*** | 242 943 | - | - | - | 243 828 |
| ***POPC:Chol (3:1)*** | 248 527 | 248 615 | 242 640 | 242 724 | 244 008 |
| ***POPC:POPE:Chol (2:1:1)*** | 241 020 | 241 034 | 244 089 | 244 173 | 245 145 |

**Supplementary Table 17. Distances used to originally restrain Mg2+-ATP-NBD arrangement in MD simulations.** Distances were measured on the minimized structure (OF *b*MRP1-$(ATP)_2$) and on the starting structure (*b*MRP1-$(ATP)_2$ and *b*MRP1-LTX-$(ATP)_2$)

| | **OF *b*MRP1-$(ATP)_2$** | | | | | ***b*MRP1-$(ATP)_2$ *b*MRP1-LTX-$(ATP)_2$** |
|---|---|---|---|---|---|---|
| | POPC | POPE | POPC:POPE (3:1) | POPC:Chol (3:1) | POPC:POPE:Chol (2:1:1) | POPC POPC:Chol (3:1) POPC:POPE:Chol (2:1:1) |
| $ATP_{NBD1}$ – Trp653 | 3.22 | 3.46 | 3.45 | 3.56 | 3.57 | 3.50 |
| $ATP_{NBD1}(P\alpha)$ – Ser686 | 2.74 | 2.80 | 2.80 | 2.78 | 2.78 | 2.71 |
| $ATP_{NBD1}(P\alpha)$ – Ser685 | 3.43 | 3.16 | 3.16 | 3.14 | 3.16 | 3.07 |
| $ATP_{NBD1}(P\beta)$ – Ser685 | 2.76 | 2.86 | 2.85 | 2.77 | 2.76 | 2.83 |
| $ATP_{NBD1}(P\beta)$ – Lys684 | 2.73 | 2.79 | 2.78 | 2.75 | 2.75 | 2.79 |
| $ATP_{NBD1}(P\gamma)$ – Ser1430 | 2.49 | 2.68 | 2.67 | 2.65 | 2.66 | - |
| $ATP_{NBD1}(P\gamma)$ – Lys684 | 2.50 | 2.64 | 2.61 | 2.68 | 2.68 | 2.71 |
| $ATP_{NBD2}$ – Tyr1301 | 3.55 | 3.52 | 3.55 | 3.52 | 3.52 | 3.60 |
| $ATP_{NBD2}(P\alpha)$ – Ser1334 | 2.61 | 2.62 | 2.62 | 2.63 | 2.63 | 2.70 |
| $ATP_{NBD2}(P\alpha)$ – Ser1334 | 3.5 | 2.99 | 2.97 | 2.96 | 3.00 | 2.62 |

| | | | | | | |
|---|---|---|---|---|---|---|
| $ATP_{NBD2}$(Pβ) – Ser1333 | 2.61 | 2.75 | 2.75 | 2.74 | 2.73 | 2.76 |
| $ATP_{NBD2}$(Pβ) – Lys1332 | 3.01 | 2.71 | 2.71 | 2.70 | 2.70 | 2.71 |
| $ATP_{NBD2}$(Pγ) – Gly771 | 3.61 | 3.00 | 3.01 | 3.00 | 3.04 | - |
| $ATP_{NBD2}$(Pγ) – Lys1332 | 2.63 | 2.74 | 2.73 | 2.77 | 2.77 | 2.74 |
| $Mg^{2+}_{NBD1}$ – $ATP_{NBD1}$(Pα) | 1.88 | 1.86 | 1.90 | 1.90 | 1.90 | 1.88 |
| $Mg^{2+}_{NBD1}$ – $ATP_{NBD1}$(Pβ) | 1.93 | 1.94 | 1.94 | 1.93 | 1.93 | 1.91 |
| $Mg^{2+}_{NBD1}$ – $ATP_{NBD1}$(Pγ) | 1.81 | 1.83 | 1.83 | 1.83 | 1.83 | 1.83 |
| $Mg^{2+}_{NBD1}$ – Ser685 | 1.95 | 1.98 | 1.98 | 1.98 | 1.98 | 2.06 |
| $Mg^{2+}_{NBD1}$ – Gln713 | 2.59 | 2.06 | 2.06 | 2.01 | 2.01 | 2.09 |
| $Mg^{2+}_{NBD2}$ – $ATP_{NBD2}$(Pα) | 1.91 | 1.95 | 1.94 | 1.95 | 1.96 | 1.93 |
| $Mg^{2+}_{NBD2}$ – $ATP_{NBD2}$(Pβ) | 1.87 | 1.89 | 1.90 | 1.90 | 1.90 | 1.90 |
| $Mg^{2+}_{NBD2}$ – $ATP_{NBD2}$(Pγ) | 2.32 | 1.86 | 1.86 | 1.87 | 1.87 | 1.87 |
| $Mg^{2+}_{NBD2}$ – Ser1333 | 2.06 | 2.00 | 1.99 | 2.01 | 2.02 | 2.05 |
| $Mg^{2+}_{NBD2}$ – Gln1374 | 2.52 | 2.03 | 2.03 | 2.03 | 2.04 | 1.97 |

**Supplementary Table 18. Force constants (in kcal.mol$^{-1}$.Å$^{-2}$) used to restrain Mg2+-ATP-NBD arrangement.**

| **Distance between ATP and the protein** | |
|---|---|
| **< 2.9 Å** | 61.6 kcal.mol$^{-1}$.Å$^{-2}$ |
| **2.9 – 3.1 Å** | 30.80 kcal.mol$^{-1}$.Å$^{-2}$ |
| **> 3.1 Å** | 12.32 kcal.mol$^{-1}$.Å$^{-2}$ |
| **Any distance between $Mg^{2+}$ and ATP or protein** | 250 kcal.mol$^{-1}$.Å$^{-2}$ |

**Supplementary Table 19. Total length of the MD simulations for each replica in nanoseconds.** In total, 112.37 μs MD simulations were conducted in the present study.

| | | IF apo *b*MRP1 | *b*MRP1-(ATP)$_2$ | *b*MRP1-LTX | *b*MRP1-LTX-(ATP)$_2$ | OF *b*MRP1-(ATP)$_2$ |
|---|---|---|---|---|---|---|
| ***POPC*** | *rep1* | 2510 | 2010 | 2010 | 2010 | 1520 |
| | *rep2* | 2510 | 2010 | 2010 | 2010 | 1520 |
| | *rep3* | 2510 | 2010 | 2010 | 2010 | 1520 |
| ***POPE*** | *rep1* | 2010 | | | | 1500 |
| | *rep2* | 2010 | - | - | - | 1520 |
| | *rep3* | 2010 | | | | 1520 |
| ***POPC:POPE (3:1)*** | *rep1* | 2510 | | | | 1520 |
| | *rep2* | 2510 | - | - | - | 1520 |
| | *rep3* | 2510 | | | | 1520 |
| ***POPC:Chol (3:1)*** | *rep1* | 2090 | 2010 | 2010 | 2010 | 1520 |
| | *rep2* | 2060 | 2010 | 2010 | 2010 | 1520 |
| | *rep3* | 2060 | 2010 | 2010 | 2010 | 1520 |
| ***POPC:POPE:Chol (2:1:1)*** | *rep1* | 2170 | 2010 | 2010 | 2010 | 2010 |
| | *rep2* | 2200 | 2010 | 2010 | 2010 | 2010 |
| | *rep3* | 2180 | 2010 | 2010 | 2010 | 2010 |

**Supplementary Table 20. Parameters used for Lipid Bilayer Deformation calculation**

| Lipid Bilayer | Compressibility Elastic modulus Ka ($1x10^{-11}$N/Ang) | Bending Modulus Kc ($1x10^{-10}$ N*Ang) | Spontaneous monolayer curvature $c_0$ $AA^{-1}$ | Coefficient of Surface Tension $\alpha$, ($1x10^{-13}$ N/Ang) |
|---|---|---|---|---|
| **POPC** | 2.55 | 9.93 | -0.0026 | 3 |
| **POPC:Chol (3:1)** | 6.73 | 11.9 | -0.0144 | 3 |
| **POPC:POPE:Chol (2:1:1)** | 6.73 | 11.9 | -0.0217 | 3 |

### Supplementary Figures

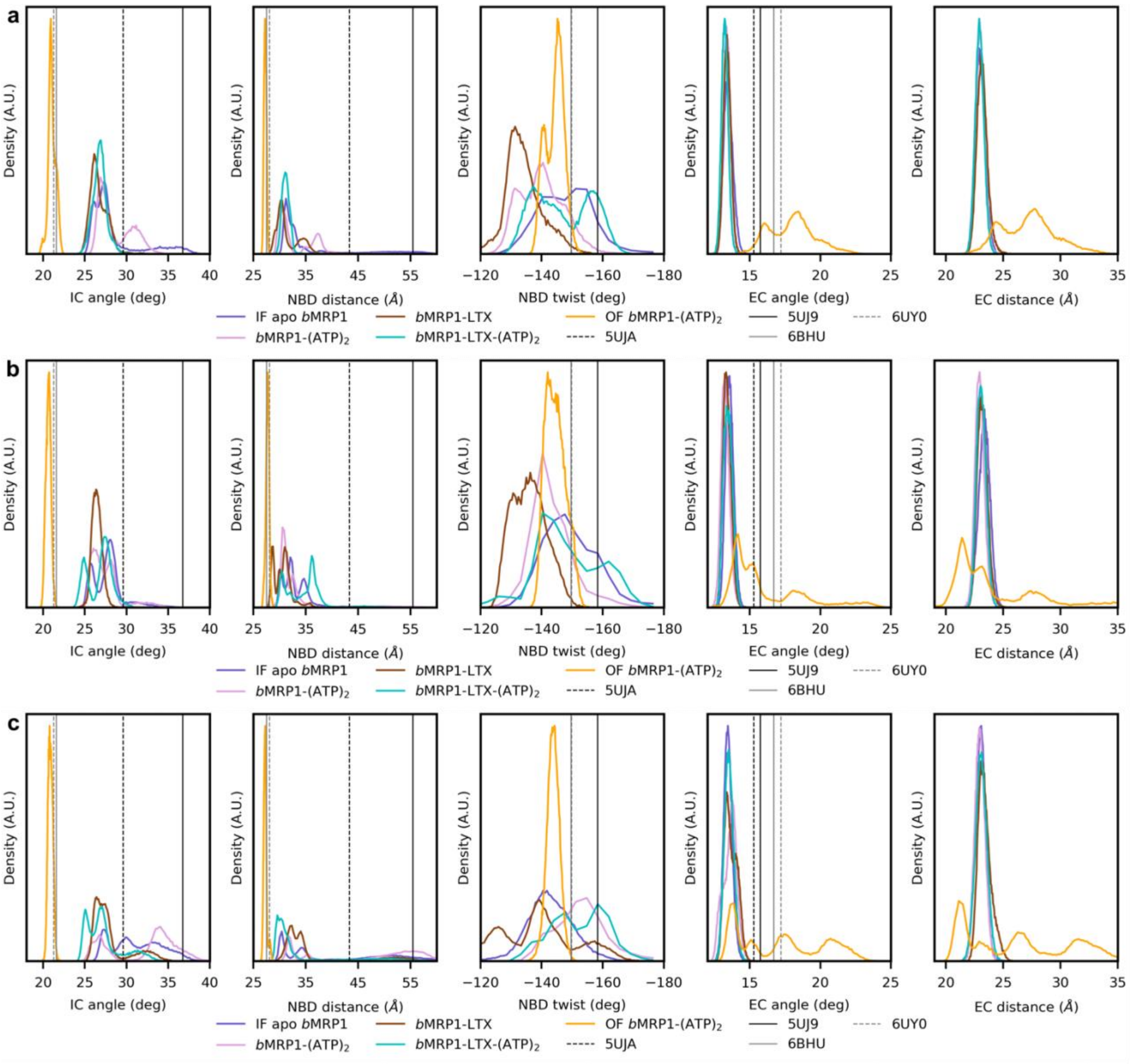

**Supplementary Figure 1. Distribution of ABC structural parameters obtained in different lipid bilayer compositions. a)** POPC:POPE:Chol (2:1:1) **b)** POPC:Chol (3:1) and **c)** POPC.

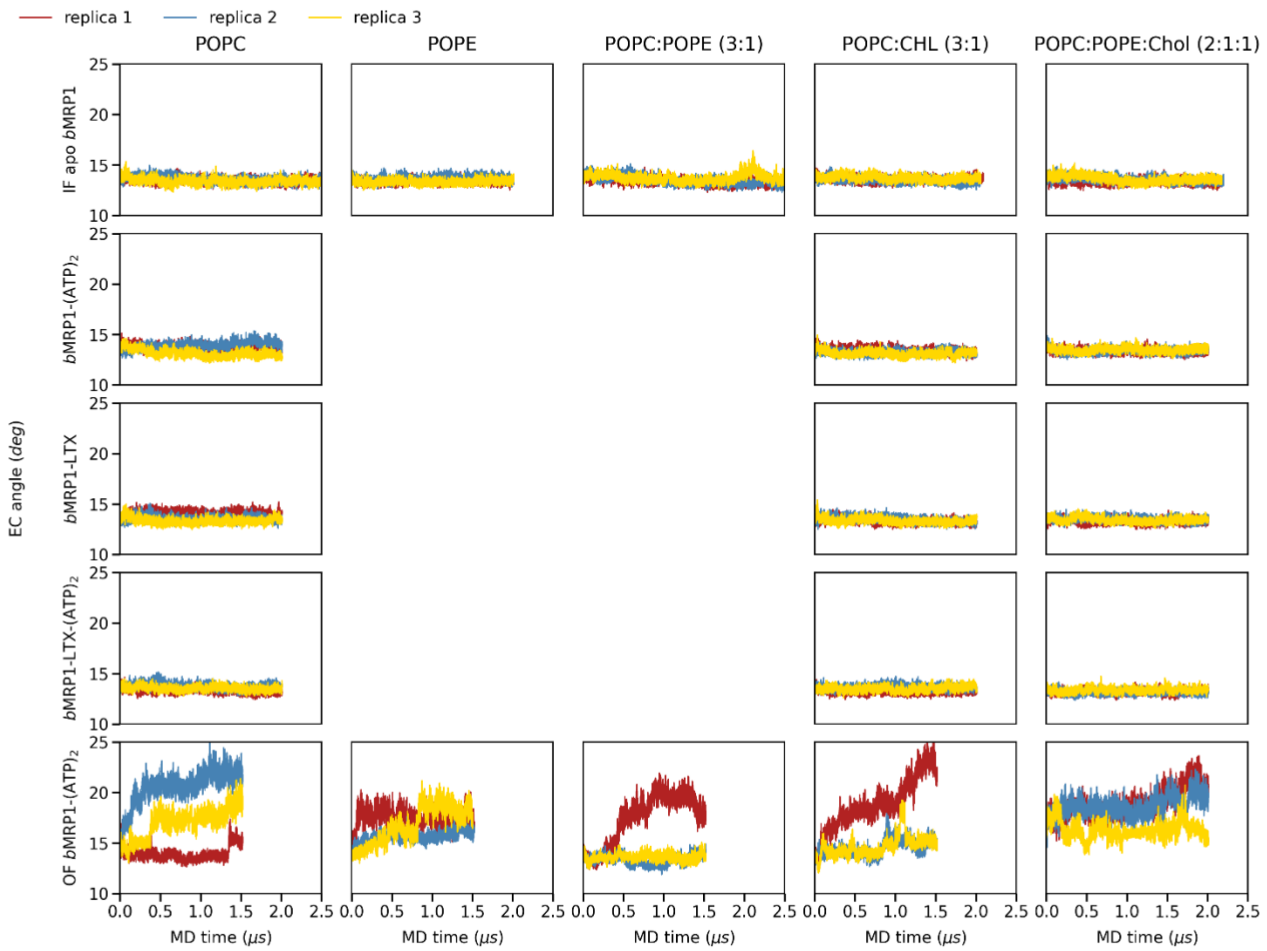

**Supplementary Figure 2. Evolution of EC angle (°) along MD simulations calculated for all the systems and for all different membrane compositions.** Replica 1, 2 and 3 are coloured red, blue and yellow, respectively.

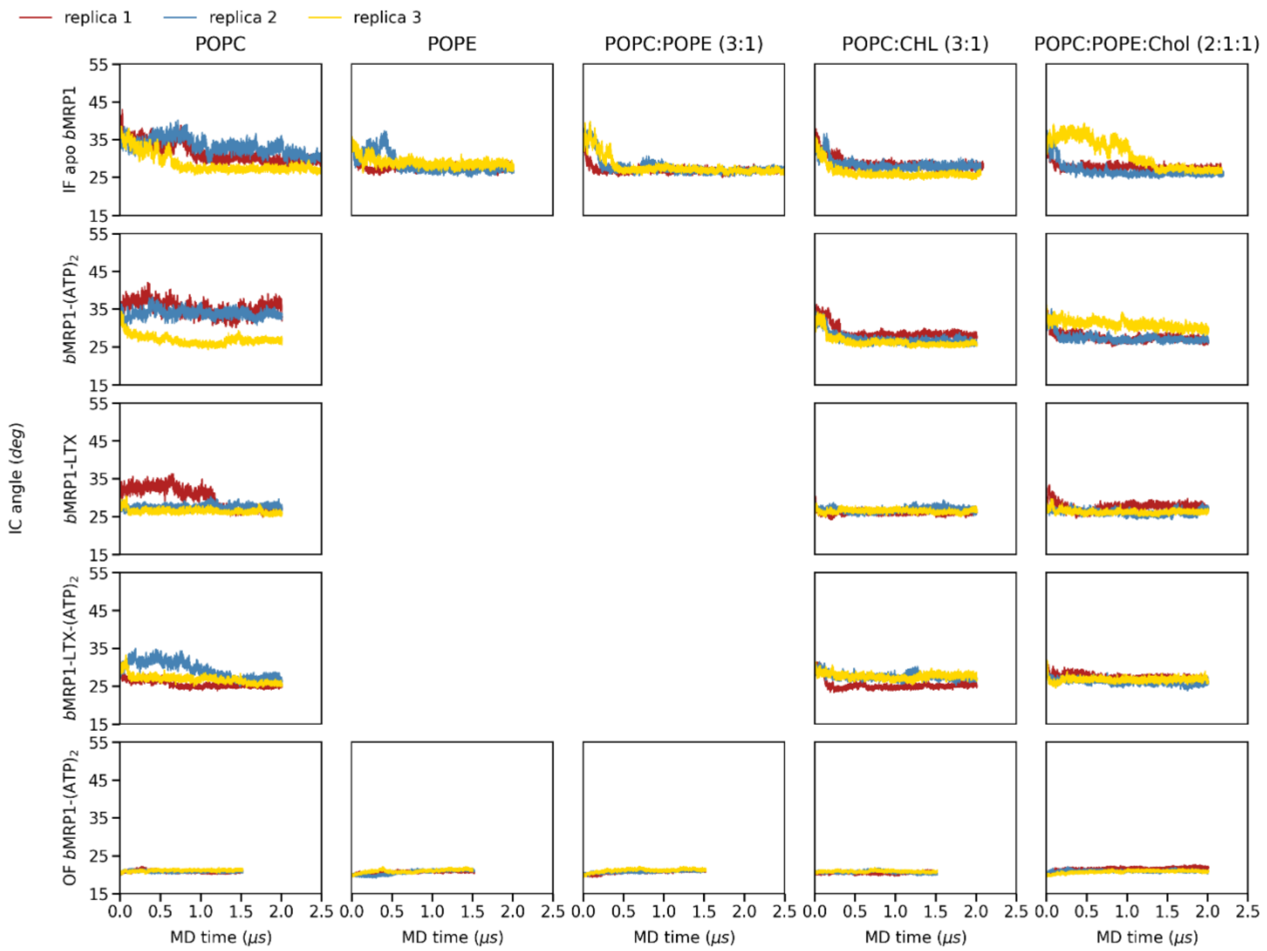

**Supplementary Figure 3. IC angle (°) during the whole simulations calculated for all the systems and for all different membrane compositions.** Replica 1, 2 and 3 are coloured red, blue and yellow, respectively.

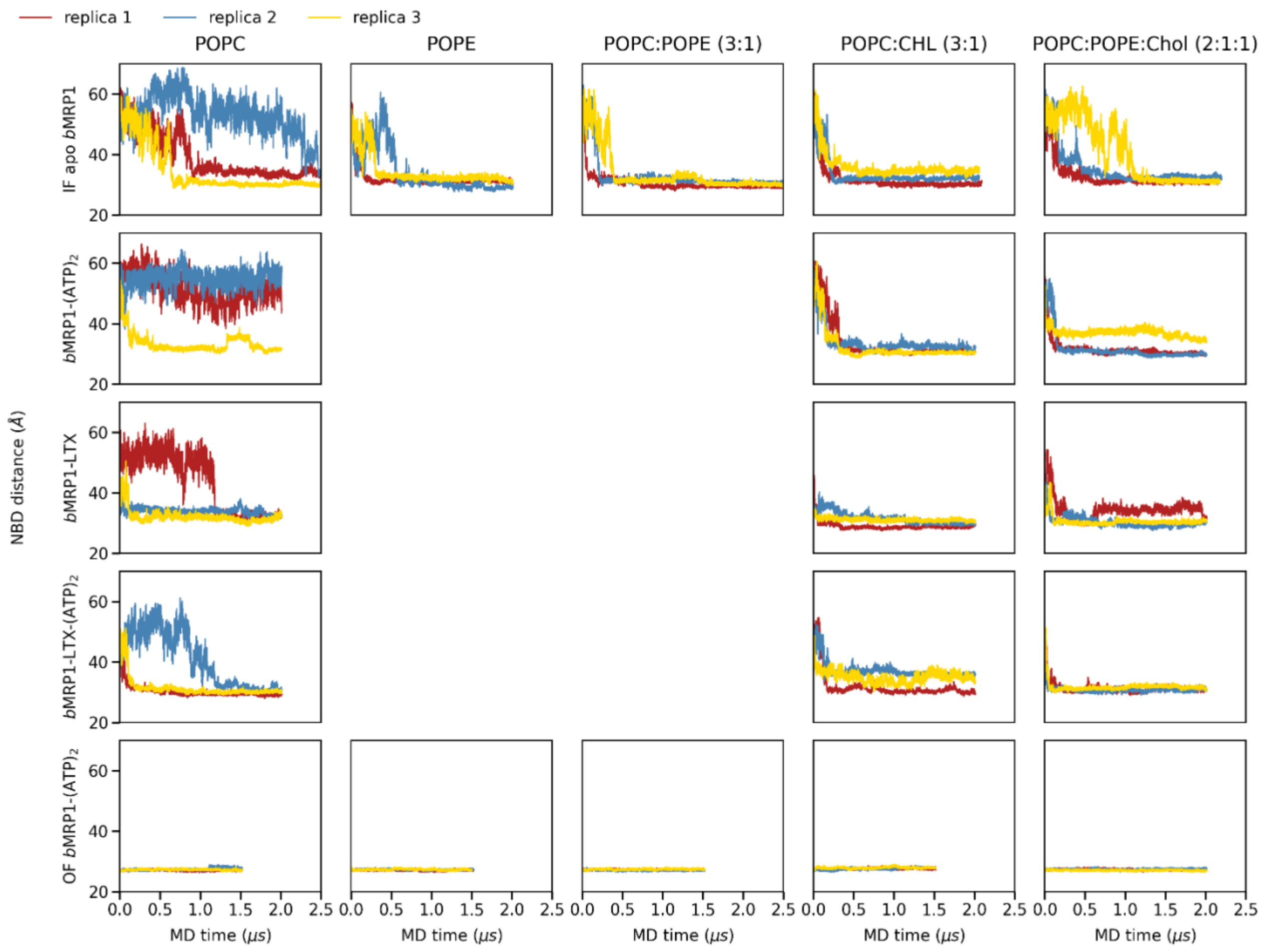

**Supplementary Figure 4. NBD distance (Å) during the whole simulations calculated for all the systems and for all different membrane compositions.** Replica 1, 2 and 3 are coloured red, blue and yellow, respectively.

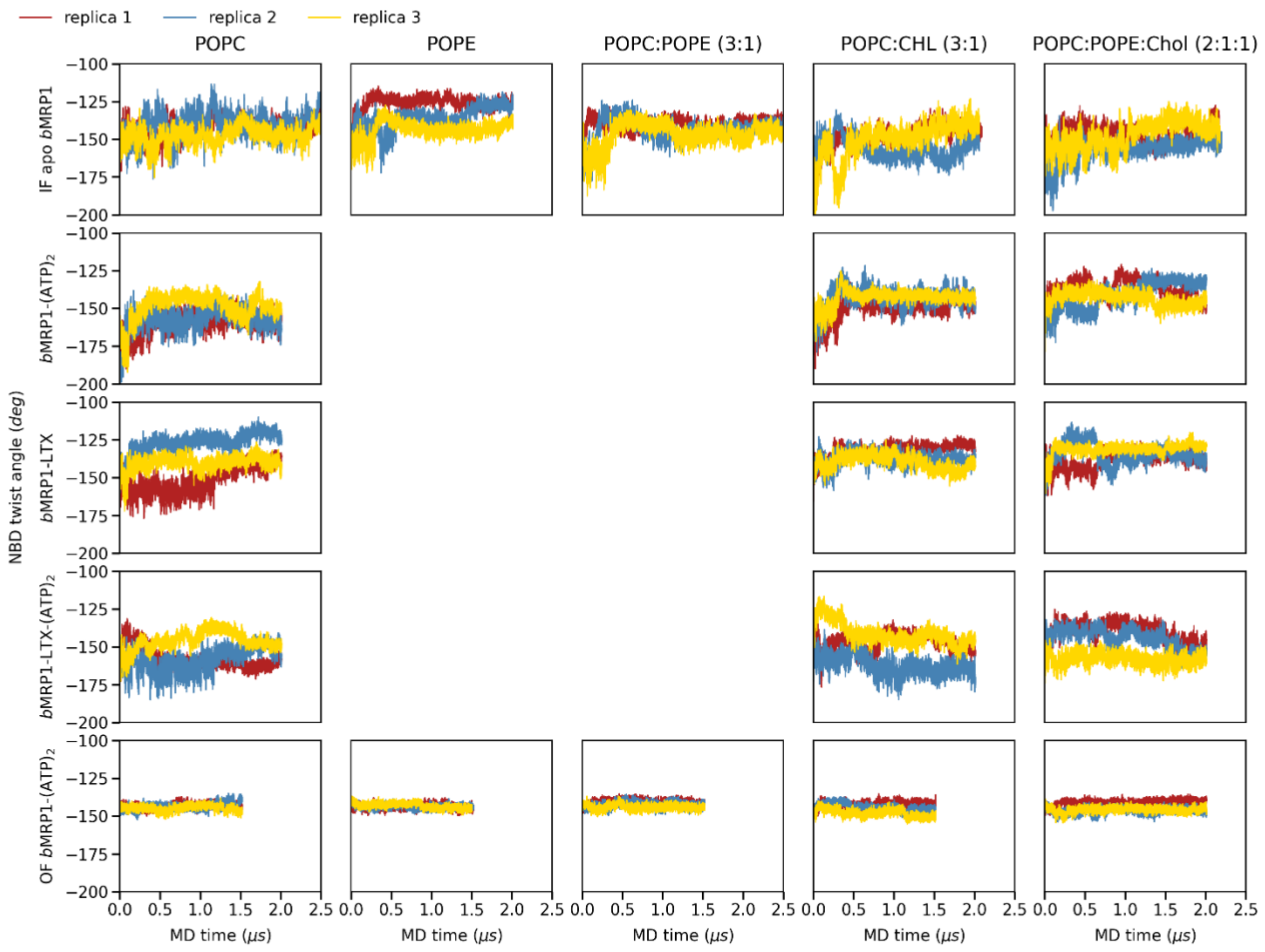

**Supplementary Figure 5. NBD twist angle (°) during the whole simulations calculated for all the systems and for all different membrane compositions.** Replica 1, 2 and 3 are coloured red, blue and yellow, respectively.

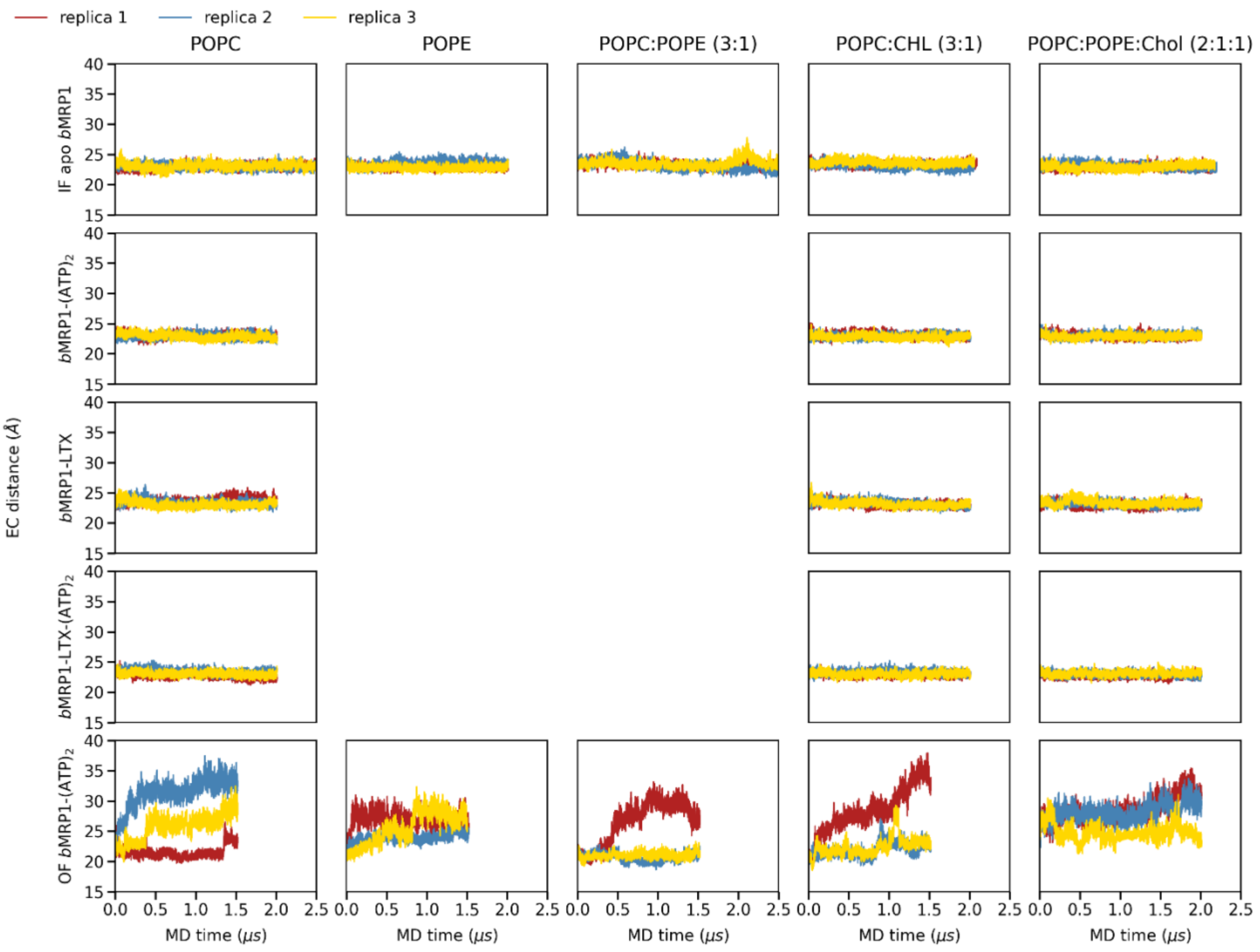

**Supplementary Figure 6. EC distance (Å) during the whole simulations calculated for all the systems and for all different membrane compositions**. Replica 1, 2 and 3 are coloured red, blue and yellow, respectively.

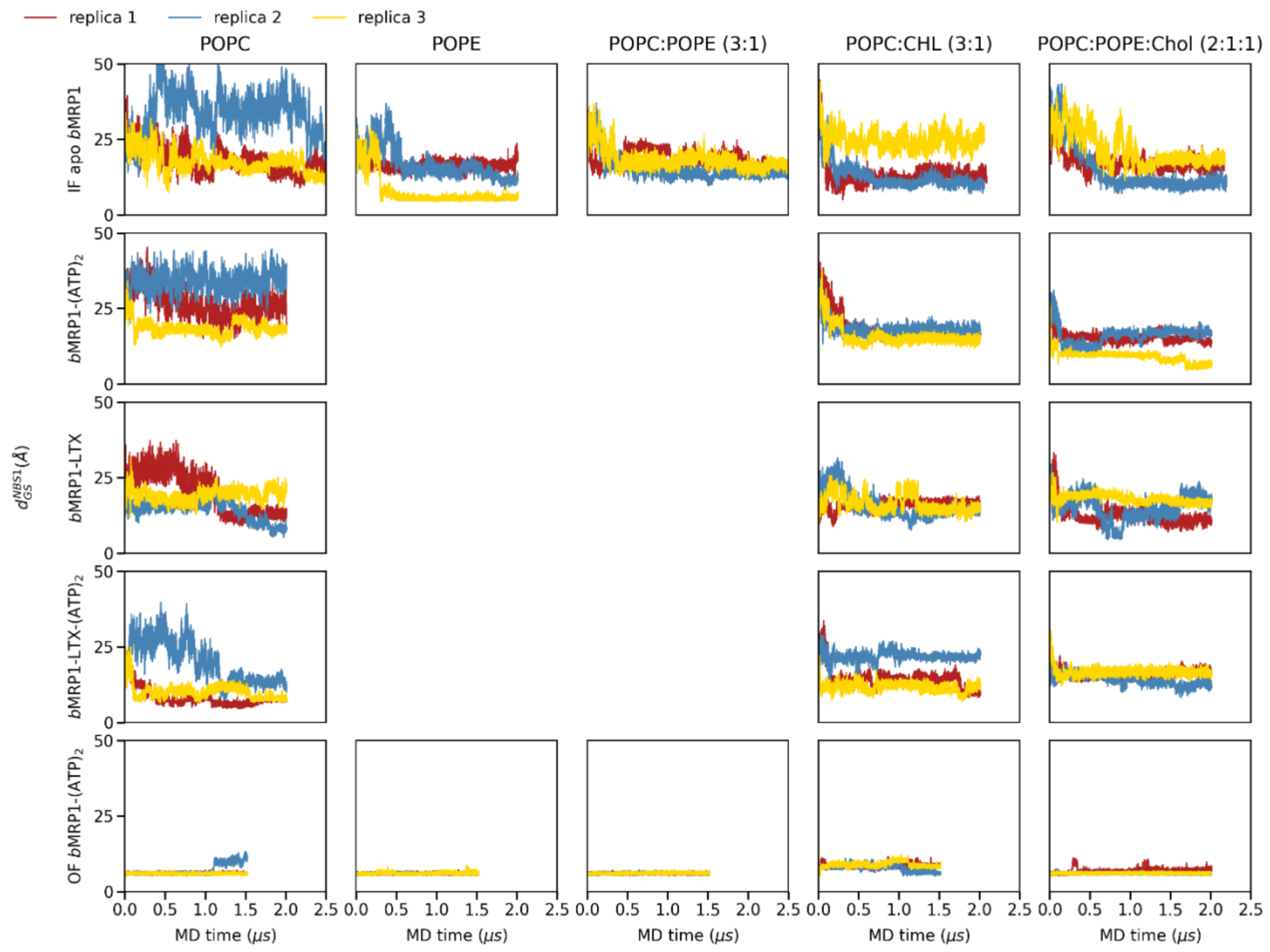

**Supplementary Figure 7. Gly681-Ser1430 ($d_{GS}^{NBS1}$) distances (Å) during the whole simulations calculated for all the systems and for all different membrane compositions**. Replica 1, 2 and 3 are coloured red, blue and yellow, respectively.

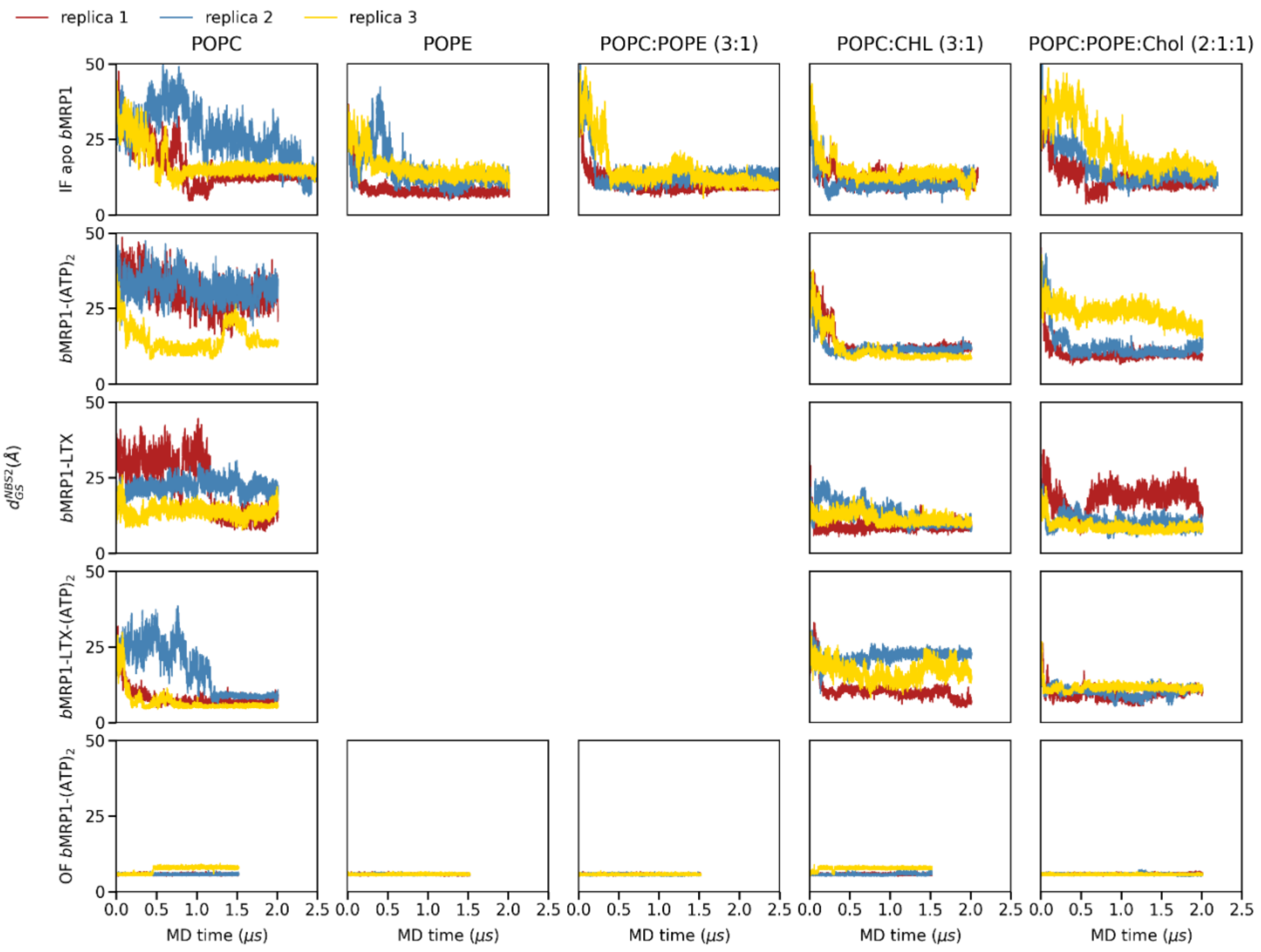

**Supplementary Figure 8. Ser769-Gly1329 ($d_{GS}^{NBS2}$) distances (Å) during the whole simulations calculated for all the systems and for all different membrane compositions.** Replica 1, 2 and 3 are coloured red, blue and yellow, respectively.

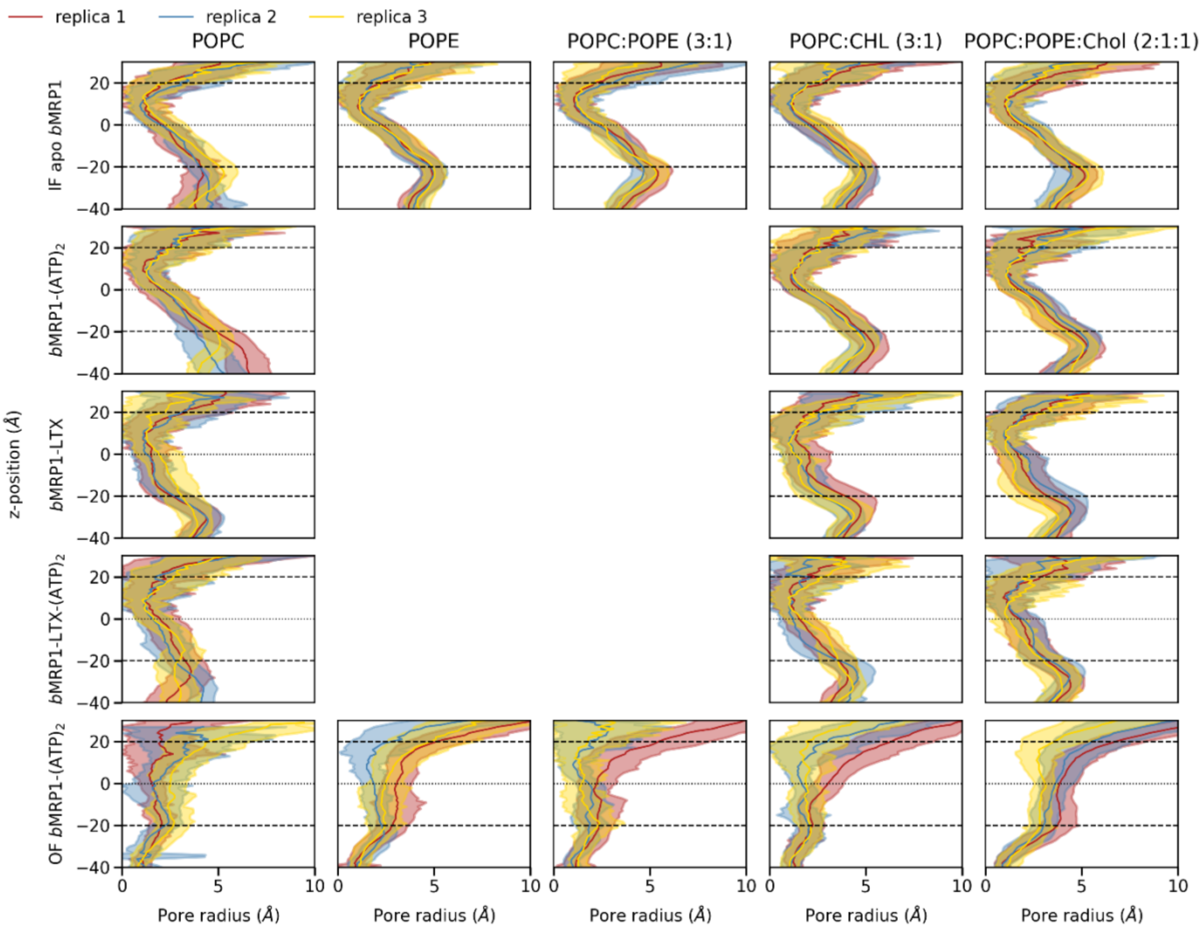

**Supplementary Figure 9. Calculated z-dependent pore radii for all systems investigated in the present study.** Replica 1, 2 and 3 are coloured red, blue and yellow, respectively.

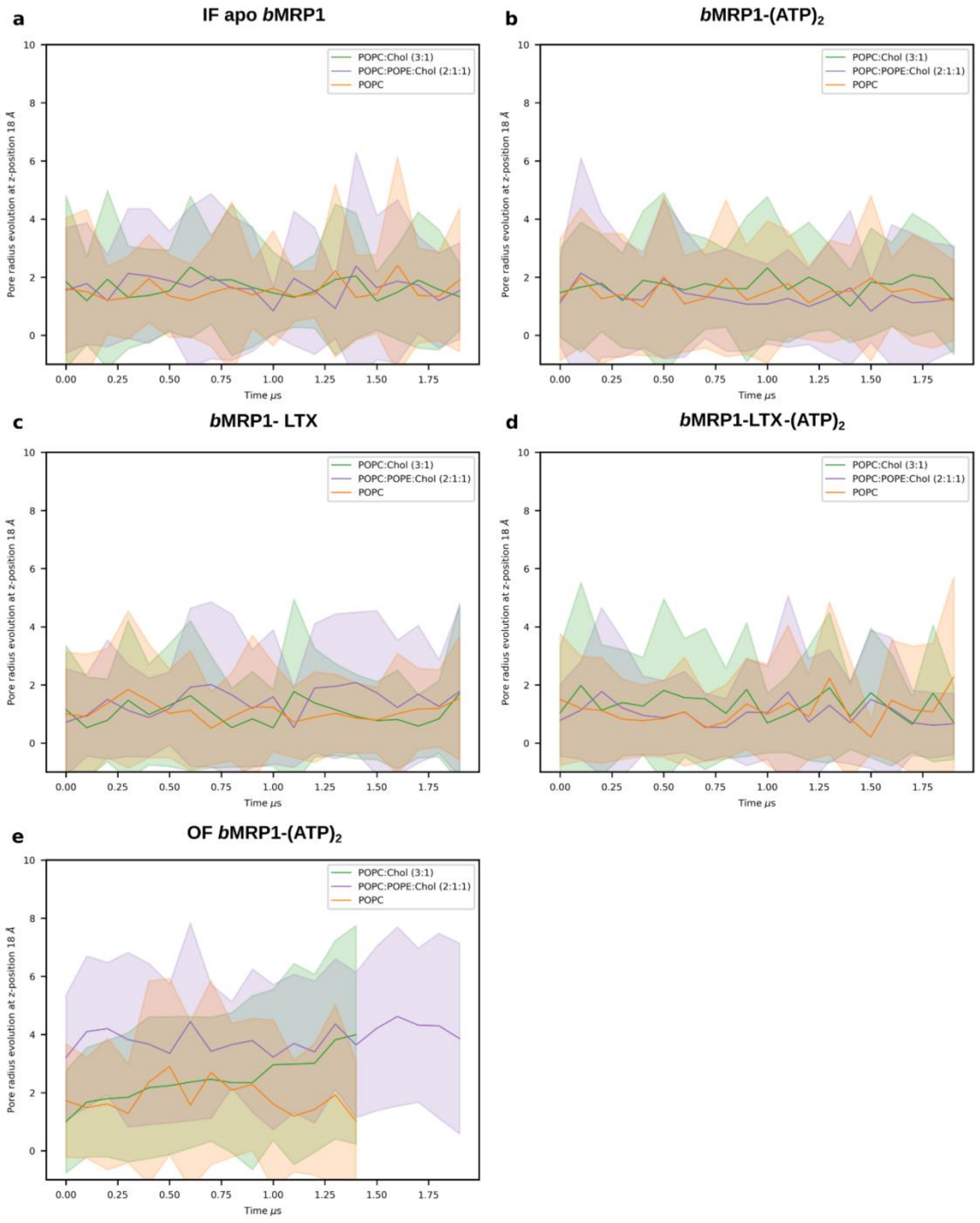

**Supplementary Figure 10. Time-dependent pore radius profiles at z = 18 Å. a)** IF apo *b*MRP1, **b)** *b*MRP1-$(ATP)_2$, **c)** *b*MRP1-LTX, **d)** *b*MRP1-LTX-$(ATP)_2$ and **e)** OF *b*MRP1-$(ATP)_2$.

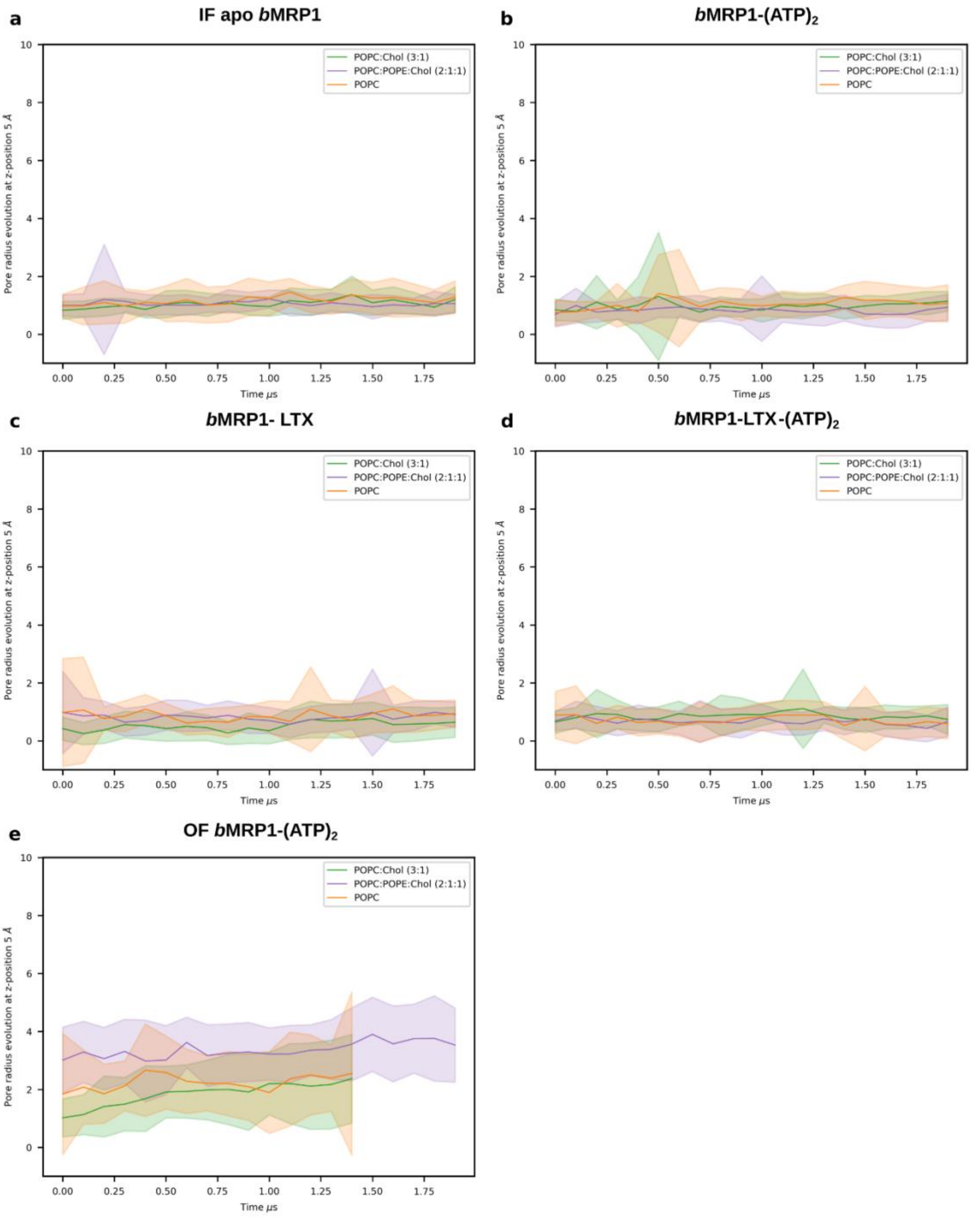

**Supplementary Figure 11. Time-dependent pore radius profiles at z = 5 Å. a)** IF apo *b*MRP1, **b)** *b*MRP1-$(ATP)_2$, **c)** *b*MRP1-LTX, **d)** *b*MRP1-LTX-$(ATP)_2$ and **e)** OF *b*MRP1-$(ATP)_2$.

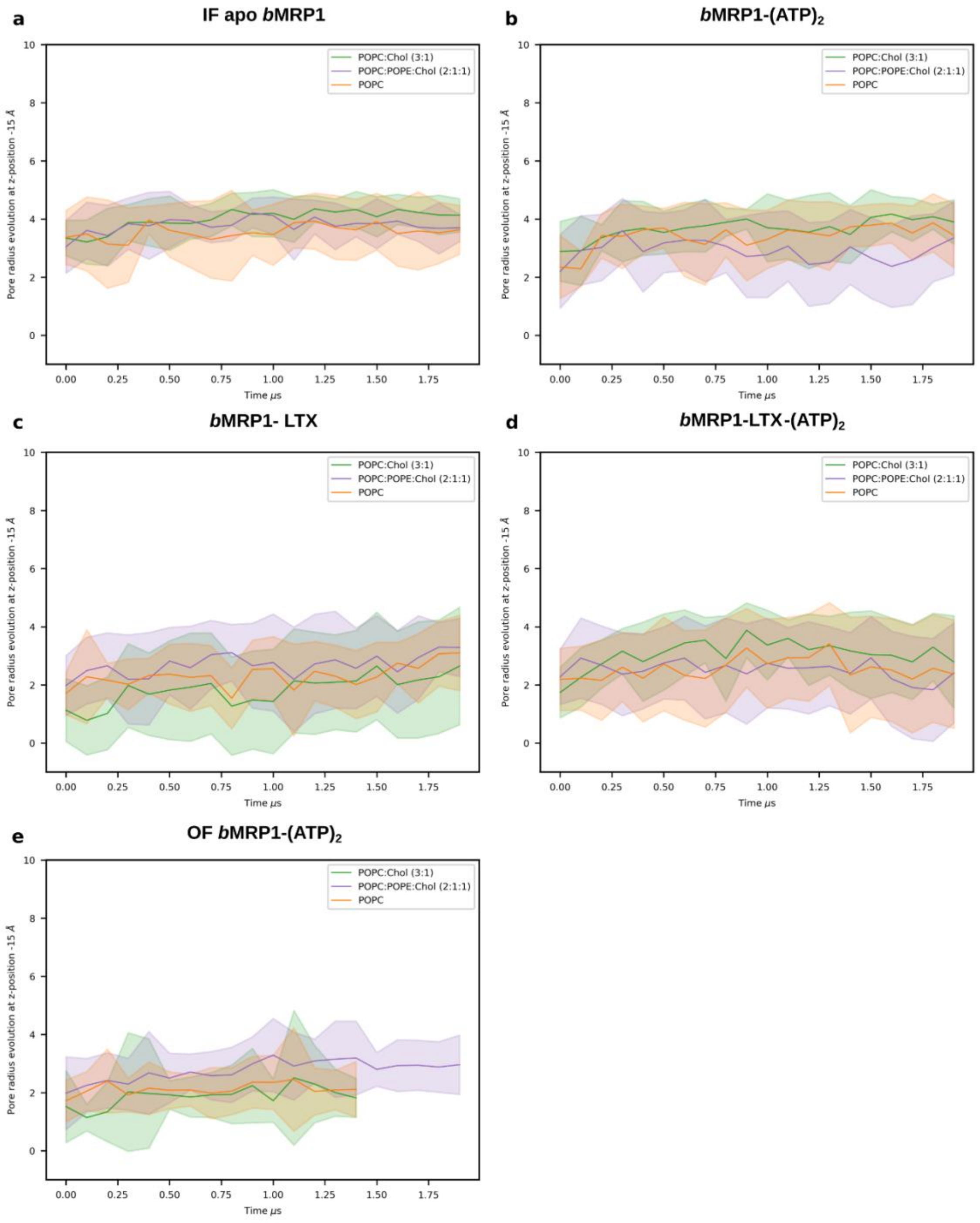

**Supplementary Figure 12. Time-dependent pore radius profiles at z = -15 Å. a)** IF apo *b*MRP1, **b)** *b*MRP1-$(ATP)_2$, **c)** *b*MRP1-LTX, **d)** *b*MRP1-LTX-$(ATP)_2$ and **e)** OF *b*MRP1-$(ATP)_2$.

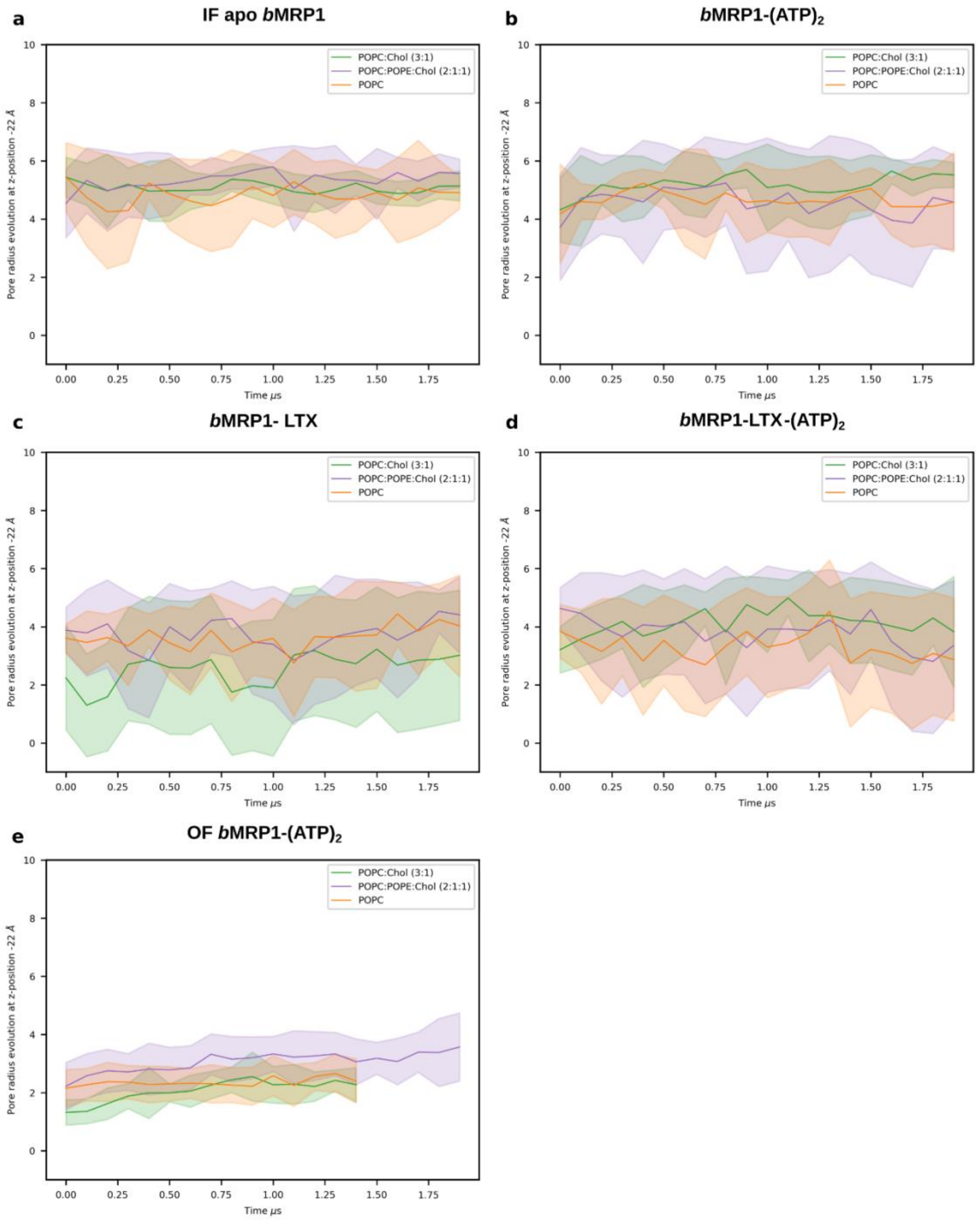

**Supplementary Figure 13. Time-dependent pore radius profiles at z = -22 Å. a)** IF apo *b*MRP1, **b)** *b*MRP1-$(ATP)_2$, **c)** *b*MRP1-LTX, **d)** *b*MRP1-LTX-$(ATP)_2$ and **e)** OF *b*MRP1-$(ATP)_2$.

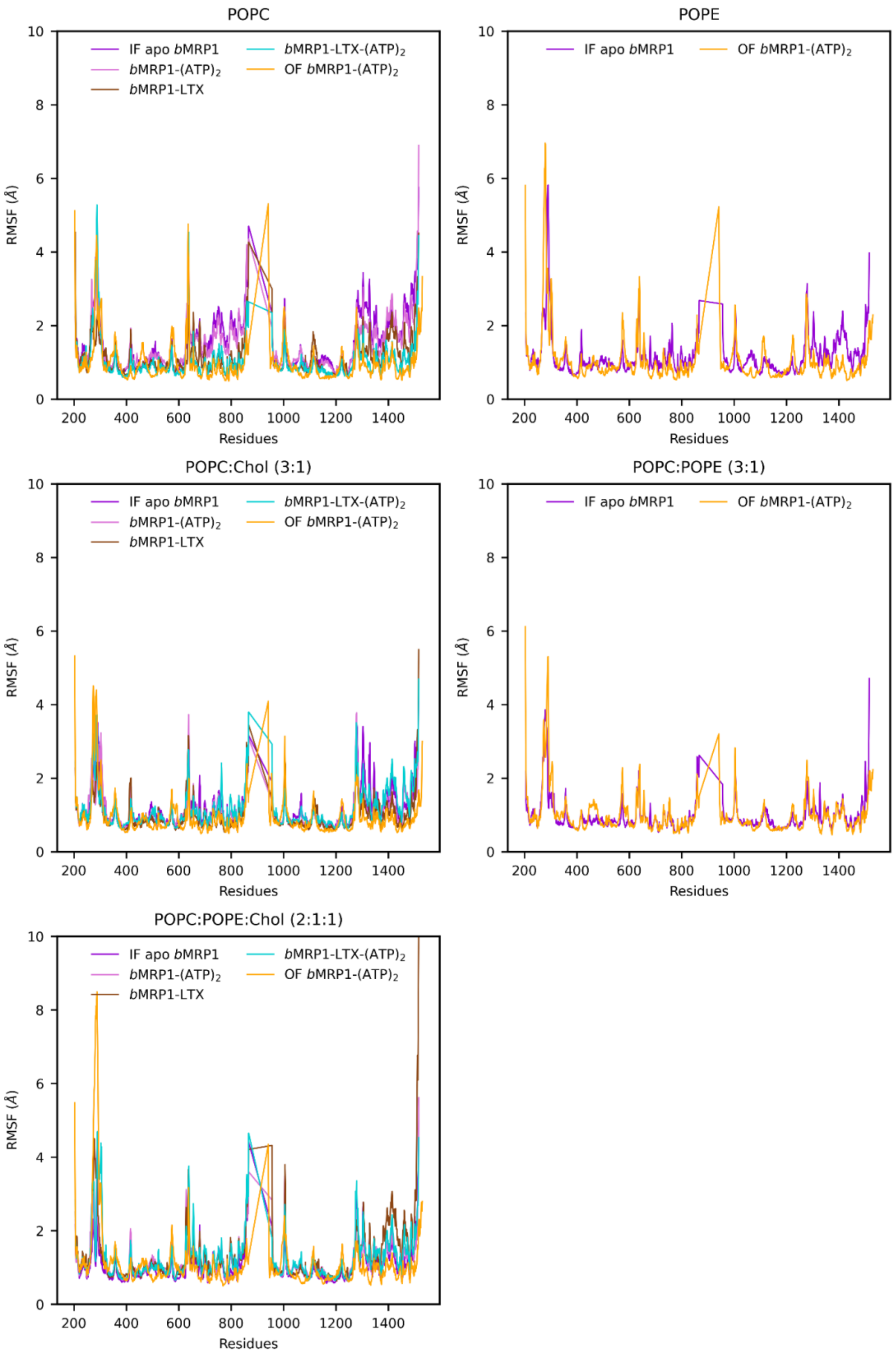

**Supplementary Figure 14. Per-residue averaged Root-mean-square fluctuations (RMSF), obtained after equilibration of the different systems in different lipid bilayer composition.**

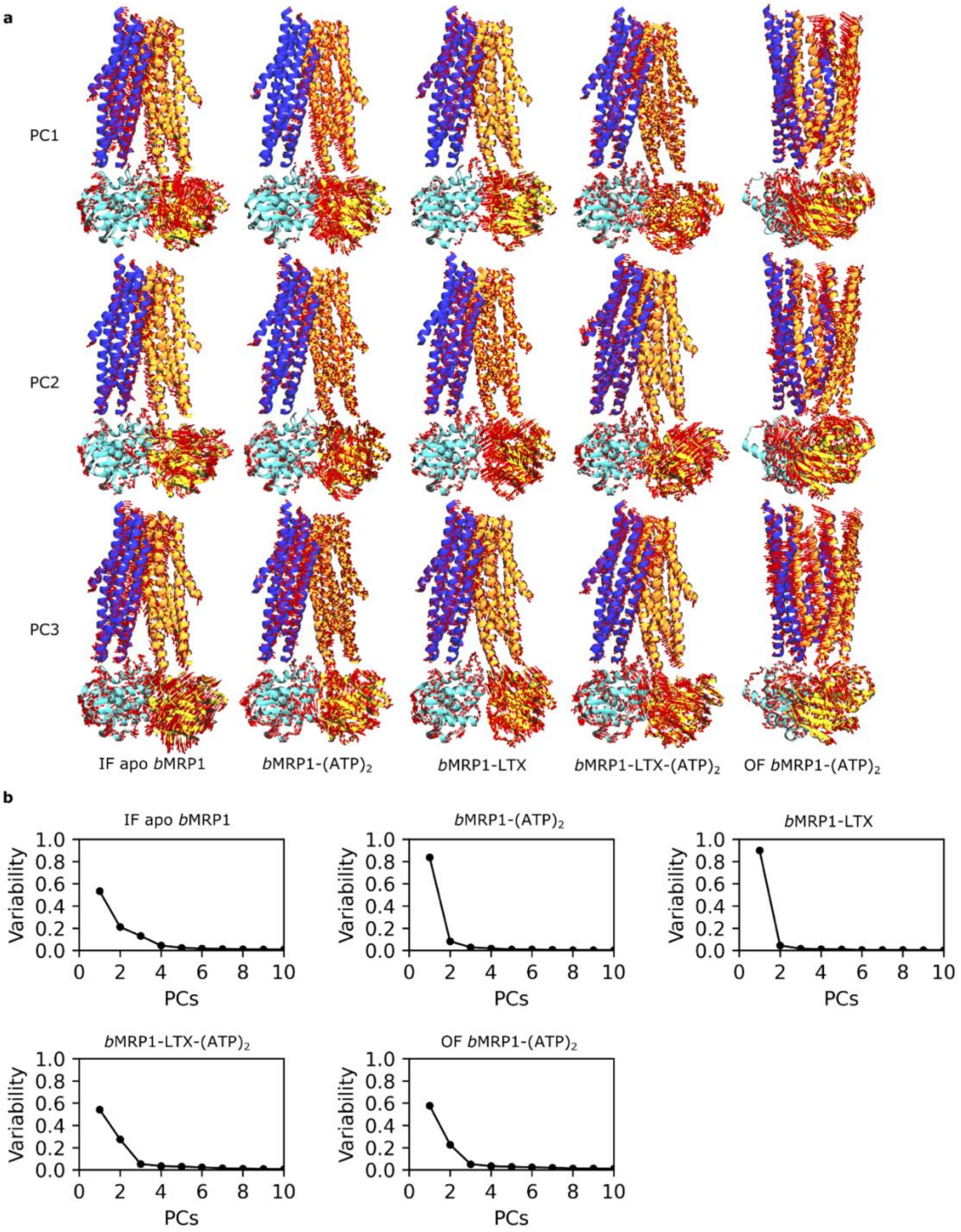

**Supplementary Figure 15. Principal component analysis (PCA) of the different systems**. For a given system, trajectories were aligned to an average structure obtained from the whole set of simulations. **a)** Arrows show the main movements in PC1-PC3, while on panel **b)** the variability of 10 PCs is shown.

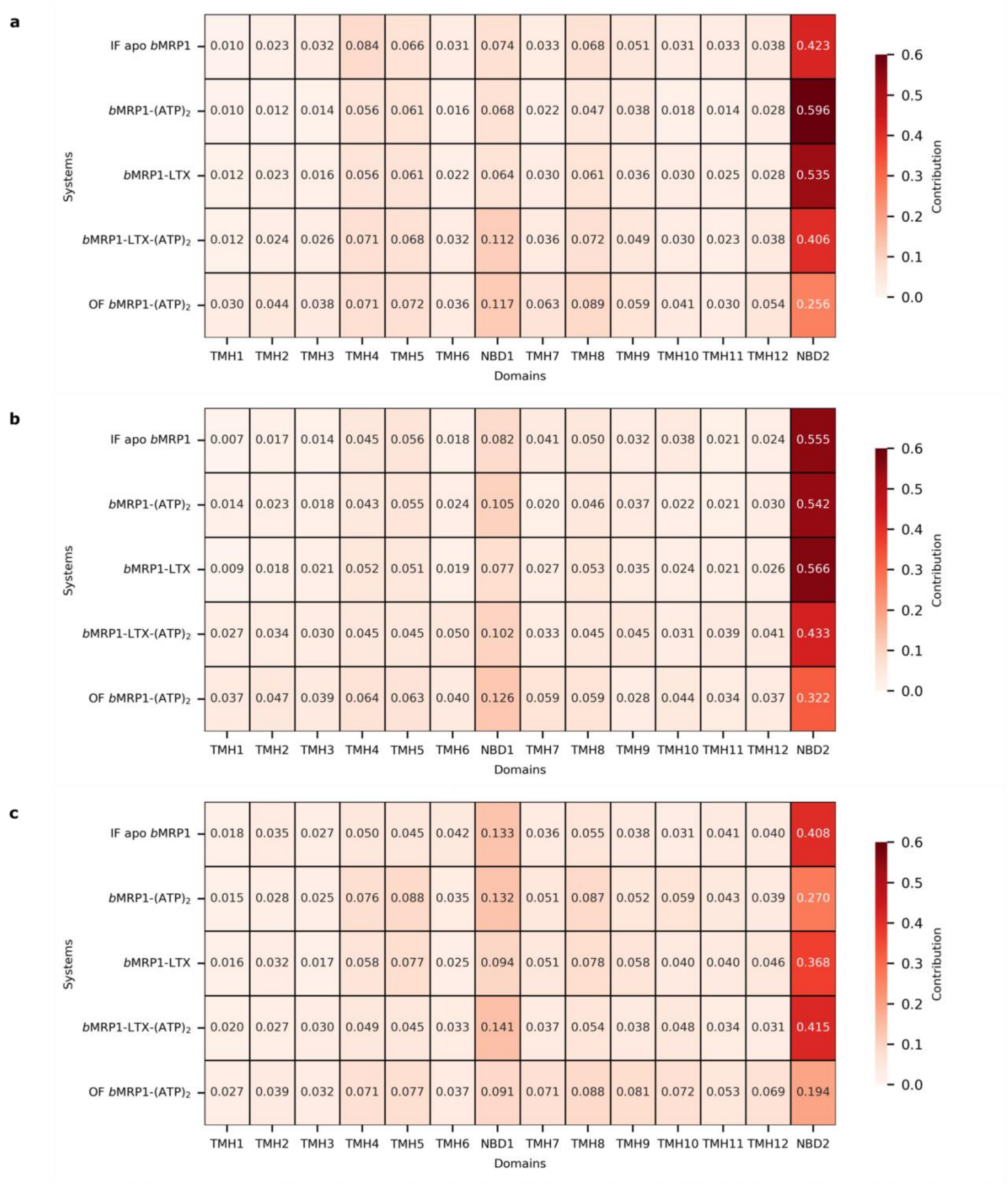

**Supplementary Figure 16. Residue contributions to the three first components. a)** PC1, **b)** PC2 and **c)** PC3 highlighting the major contribution of NBD2.

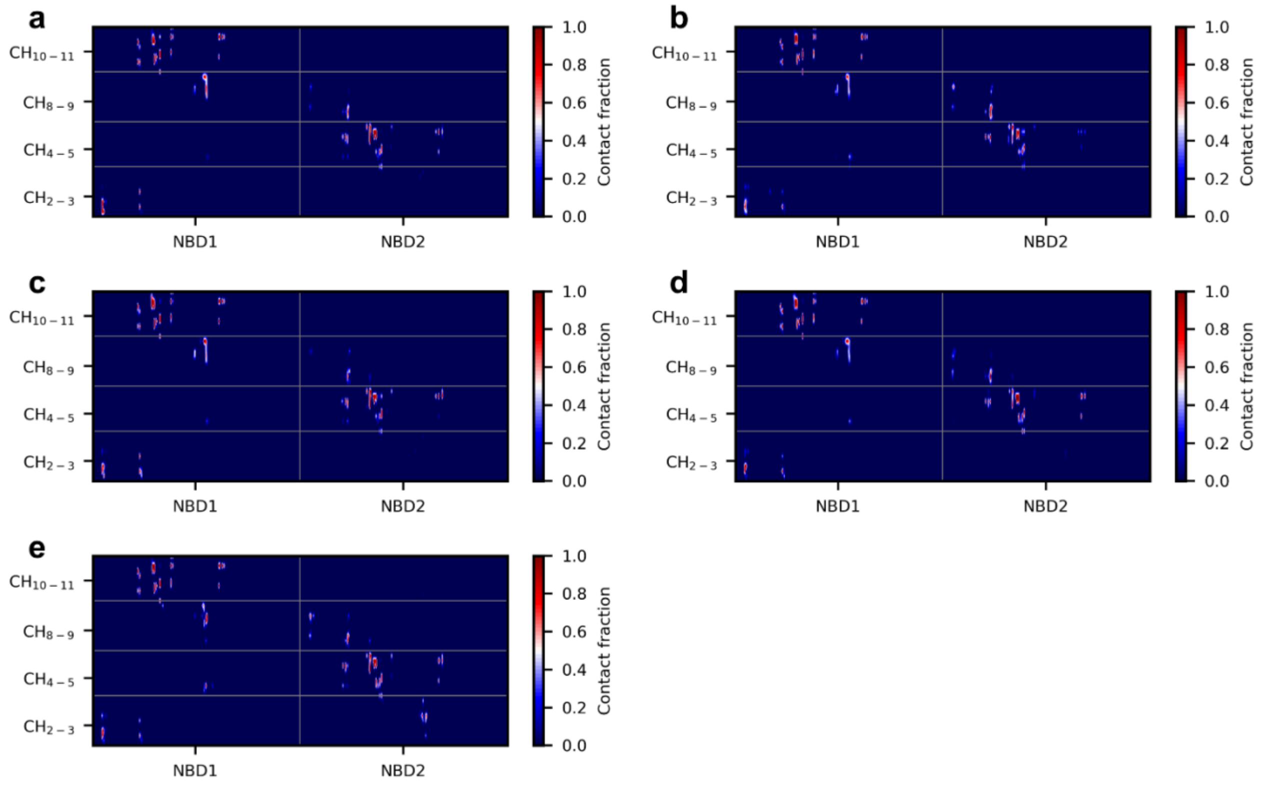

**Supplementary Figure 17. Contact maps between NBDs and Coupling Helices (CHs) obtained from MD simulations performed in POPC:POPE:Chol (2:1:1). a)** IF apo *b*MRP1, **b)** *b*MRP1-$(ATP)_2$, **c)** *b*MRP1-LTX, **d)** *b*MRP1-LTX-$(ATP)_2$ and **e)** OF *b*MRP1-$(ATP)_2$.

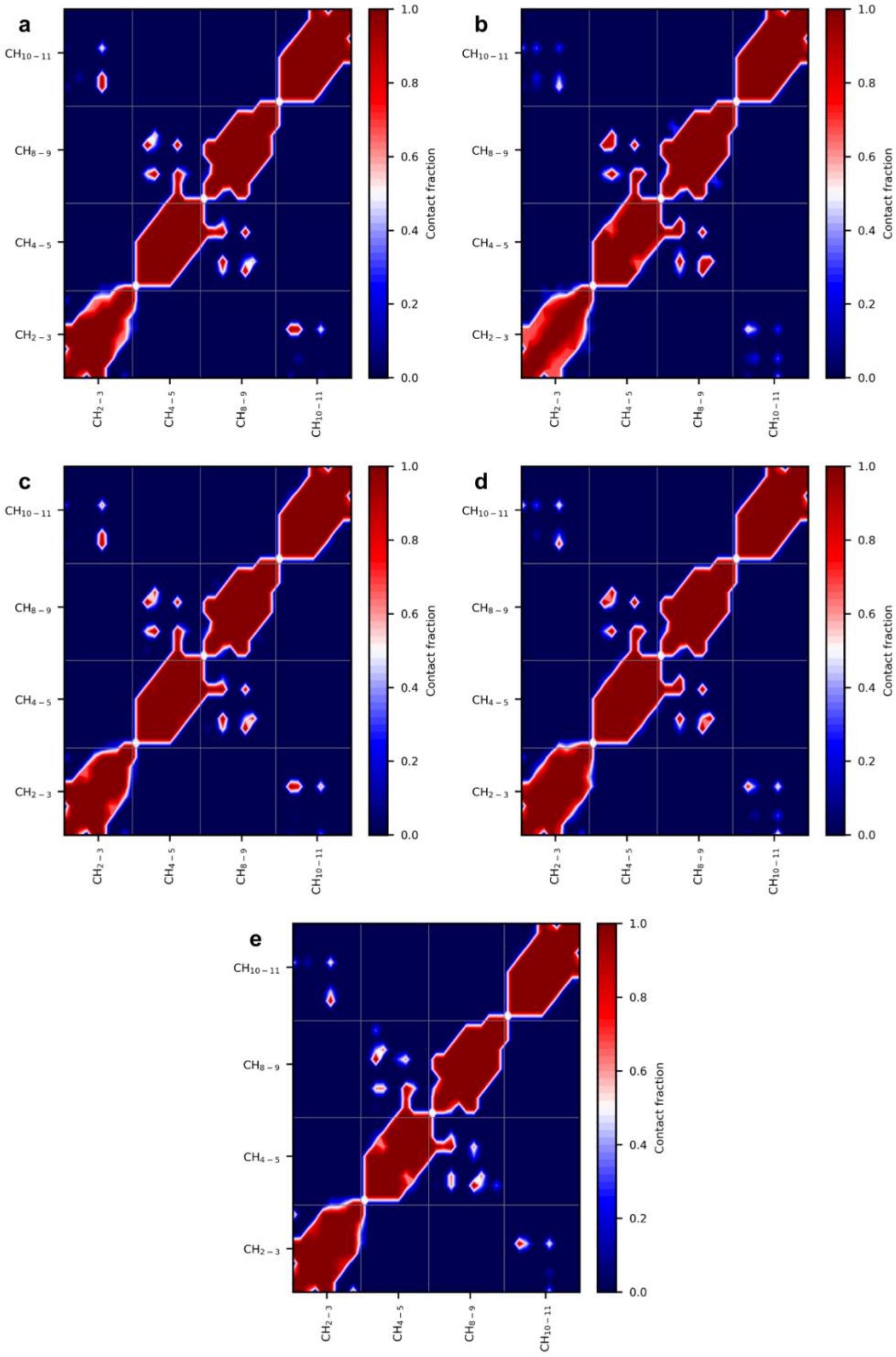

**Supplementary Figure 18. Contact maps between Coupling Helices (CHs) obtained from MD simulations performed in POPC:POPE:Chol (2:1:1). a)** IF apo *b*MRP1, **b)** *b*MRP1-$(ATP)_2$, **c)** *b*MRP1-LTX, **d)** *b*MRP1-LTX-$(ATP)_2$ and **e)** OF *b*MRP1-$(ATP)_2$.

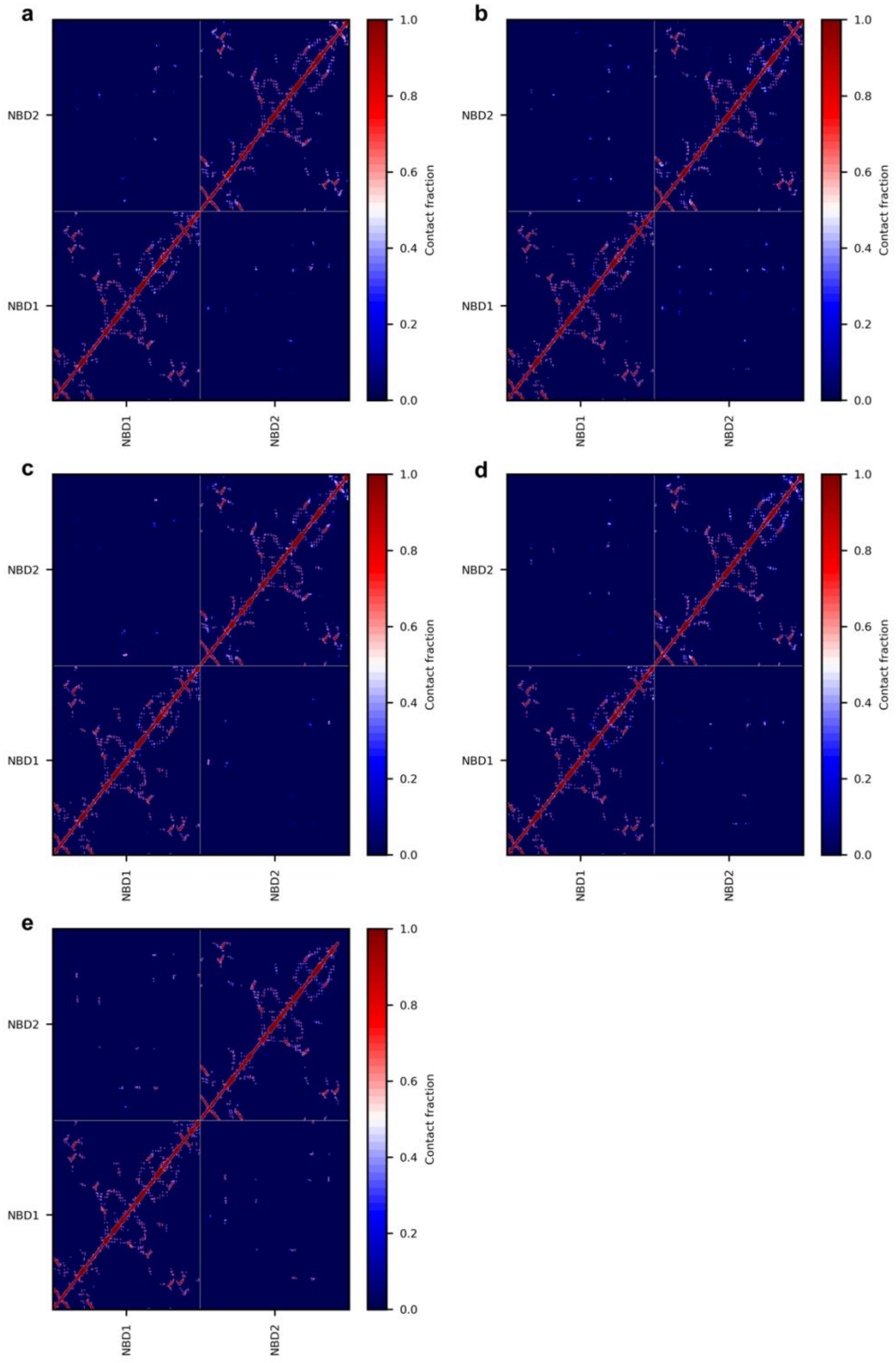

**Supplementary Figure 19. Contact maps between NBDs obtained from MD simulations performed in POPC:POPE:Chol (2:1:1). a)** IF apo *b*MRP1, **b)** *b*MRP1-$(ATP)_2$, **c)** *b*MRP1-LTX, **d)** *b*MRP1-LTX-$(ATP)_2$ and **e)** OF *b*MRP1-$(ATP)_2$.

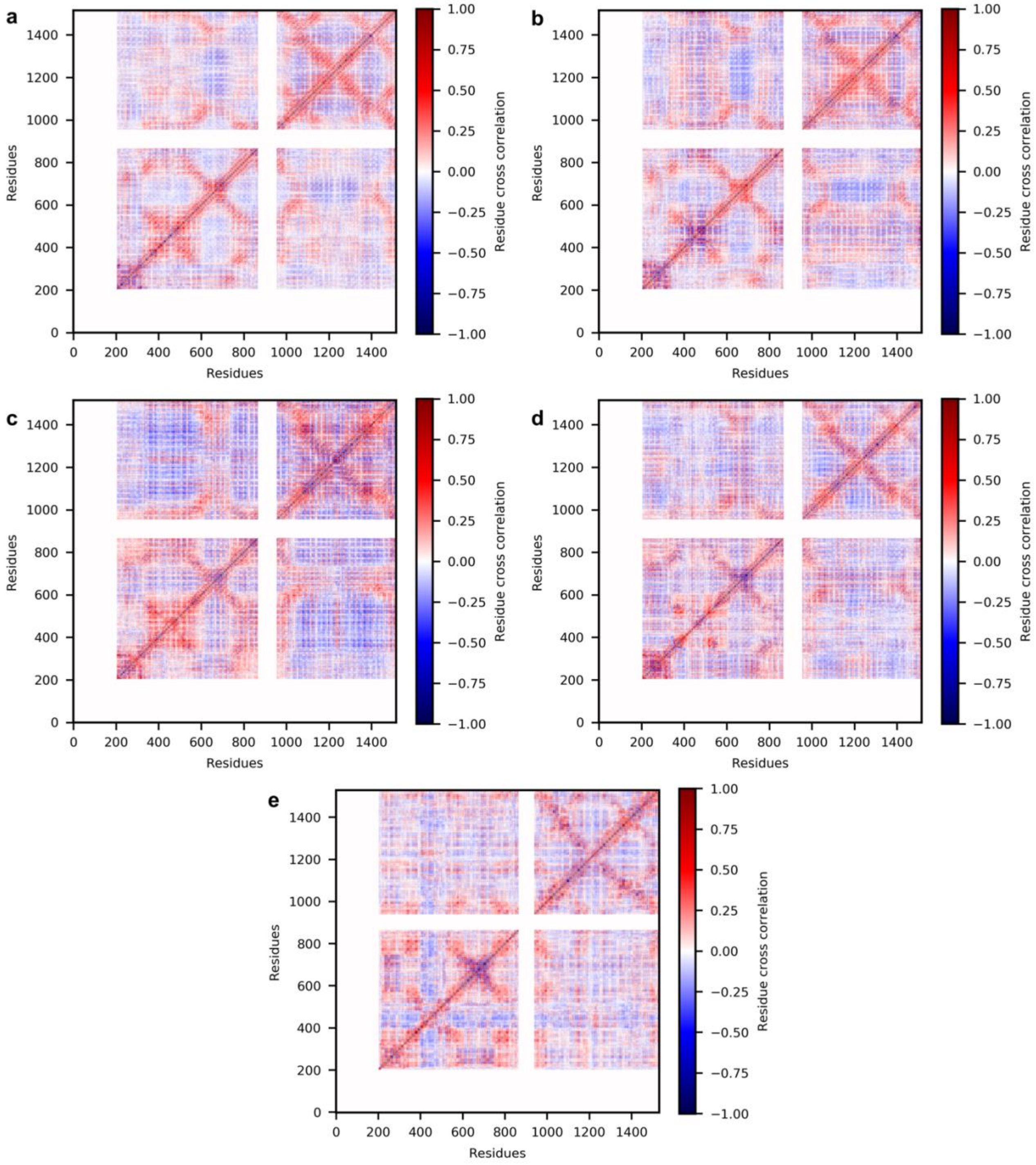

**Supplementary Figure 20. Overall dynamic cross-correlation matrices from MD simulations performed POPC:POPE:Chol (2:1:1). a)** IF apo *b*MRP1, **b)** *b*MRP1-$(ATP)_2$, **c)** *b*MRP1-LTX, **d)** *b*MRP1-LTX-$(ATP)_2$ and **e)** OF *b*MRP1-$(ATP)_2$.

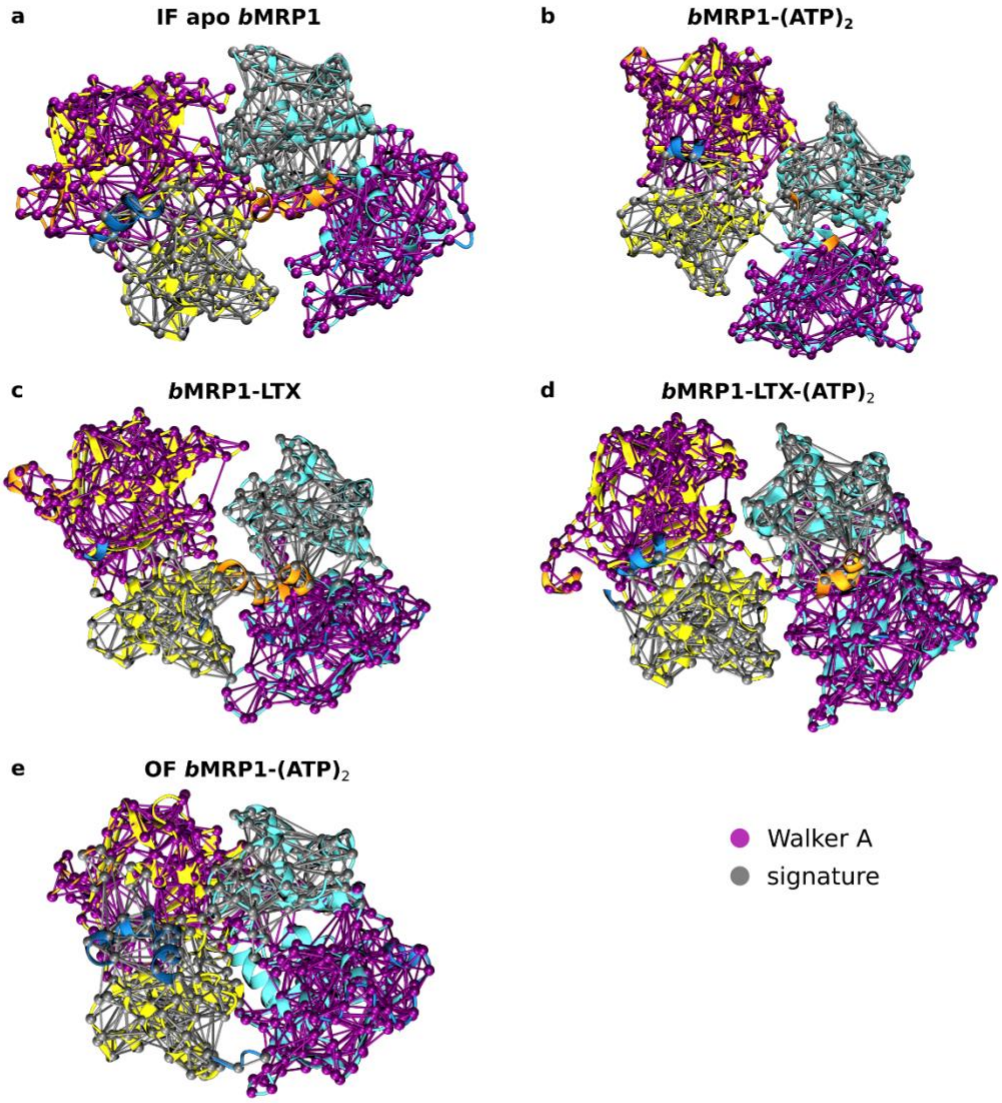

**Supplementary Figure 21. Representative NBD communities from Network Analyses calculated from MD simulations performed in POPC:POPE:Chol (2:1:1) simulations. a)** IF apo *b*MRP1, **b)** *b*MRP1-$(ATP)_2$, **c)** *b*MRP1-LTX, **d)** *b*MRP1-LTX-$(ATP)_2$ and **e)** OF *b*MRP1-$(ATP)_2$. NBD1 and NBD2 are coloured yellow and cyan, respectively. Orange and blue colours show the TMH parts involved in the communities. The Walker A community is purple, while the signature sequence community silver. NBD dimers are shown from upper view.

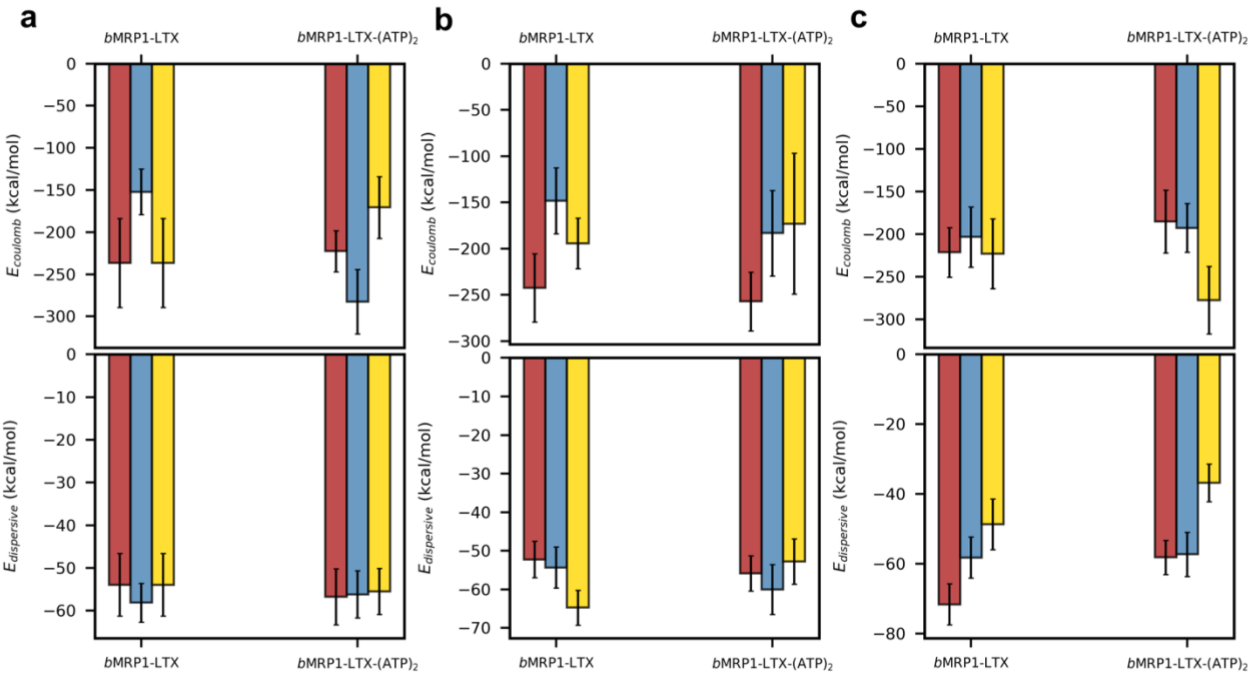

**Supplementary Figure 22. Non-covalent interaction energies between LTX and *b*MRP1 extracted from Coulomb (top) and van der Waals (bottom) potentials obtained from MD simulations performed in a)** POPC:POPE:Chol (2:1:1), **b)** POPC:Chol (3:1) and **c)** POPC lipid bilayers.

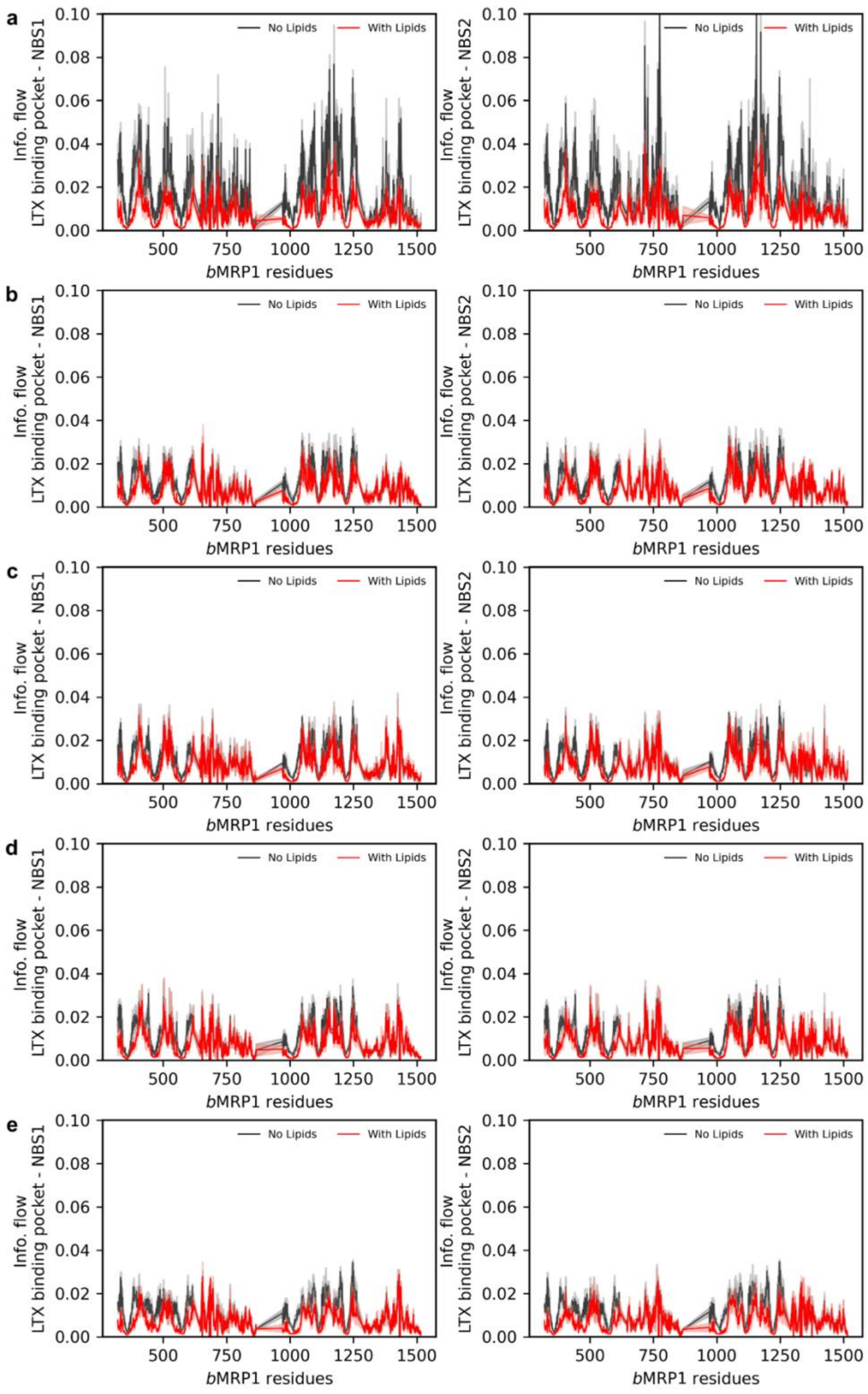

**Supplementary Figure 23. Calculated per-residue betweennesses in the allosteric pathway from substrate binding pocket to NBS1 (left) and NBS2 (right). a)** IF apo *b*MRP1**, b)** *b*MRP1-$(ATP)_2$, **c)** *b*MRP1-LTX, **d)** *b*MRP1-LTX-$(ATP)_2$ and **e)** OF *b*MRP1-$(ATP)_2$ systems embedded in POPC.

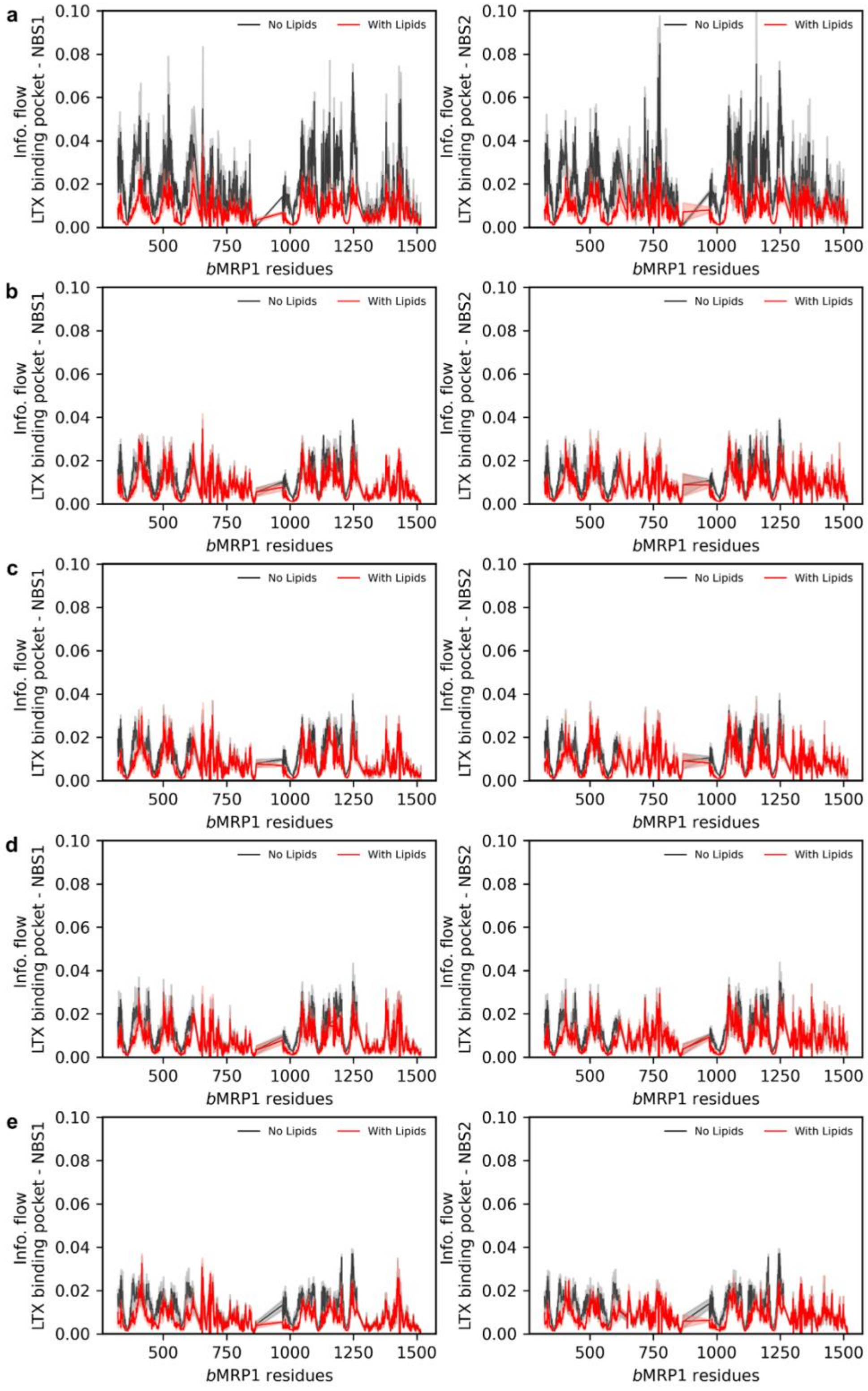

**Supplementary Figure 24. Calculated per-residue betweennesses in the allosteric pathway from substrate binding pocket to NBS1 (left) and NBS2 (right). a)** IF apo *b*MRP1, **b)** *b*MRP1-$(ATP)_2$, **c)** *b*MRP1-LTX, **d)** *b*MRP1-LTX-$(ATP)_2$ and **e)** OF *b*MRP1-$(ATP)_2$ systems embedded in POPC:Chol (3:1).

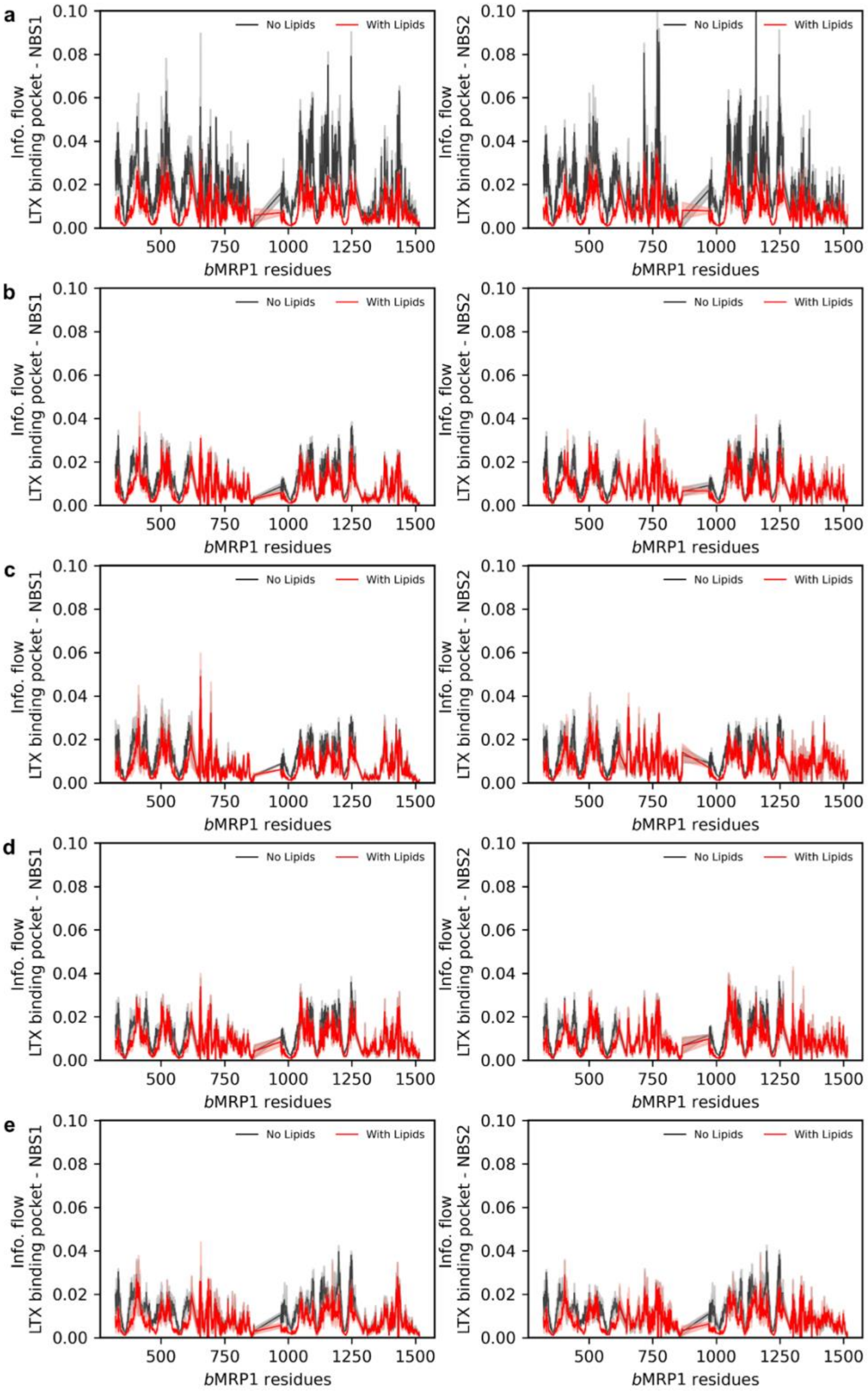

**Supplementary Figure 25. Calculated per-residue betweennesses in the allosteric pathway from substrate binding pocket to NBS1 (left) and NBS2 (right). a)** IF apo *b*MRP1, **b)** *b*MRP1-$(ATP)_2$, **c)** *b*MRP1-LTX, **d)** *b*MRP1-LTX-$(ATP)_2$ and **e)** OF *b*MRP1-$(ATP)_2$ systems embedded in POPC:POPE:Chol (2:1:1).

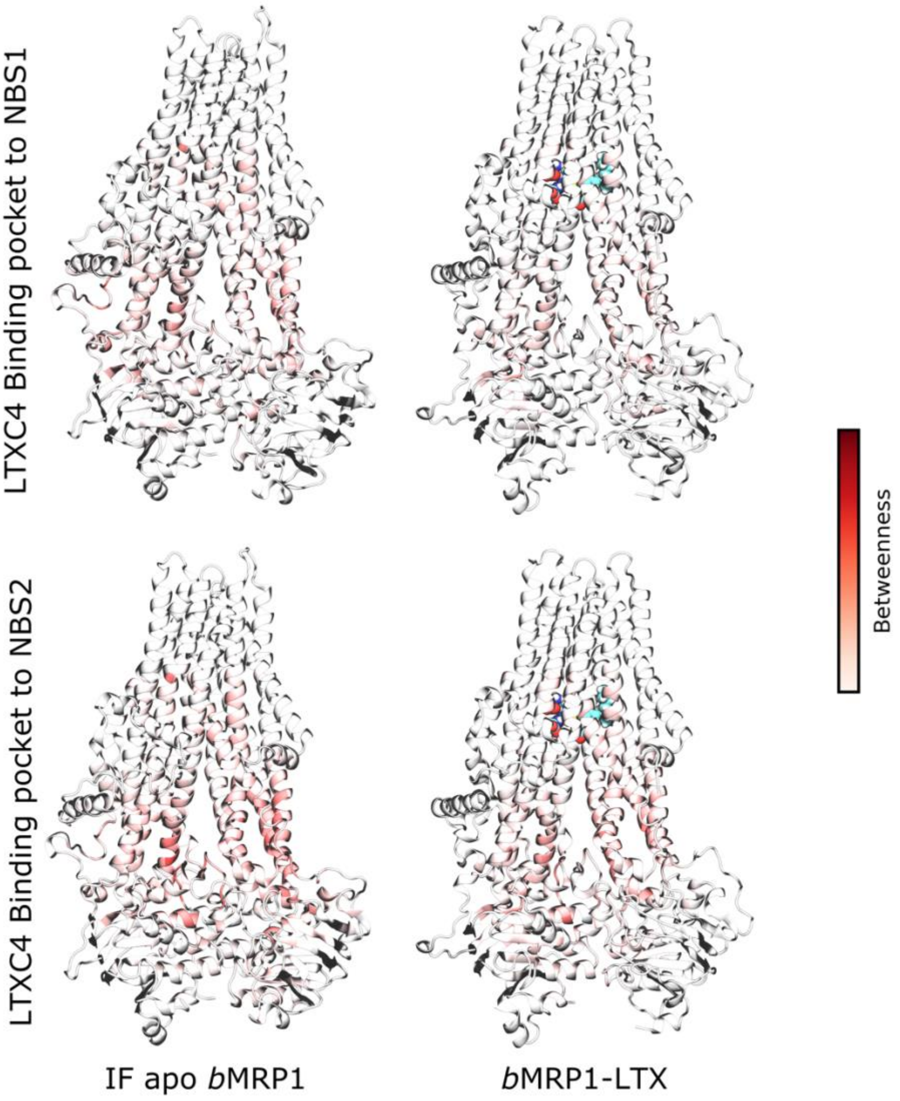

**Supplementary Figure 26. Average normalized information flow through *b*MRP1 protein regarding the allosteric communication from substrate binding pocket to NBS1 and NBS2 for IF apo *b*MRP1 and *b*MRP1-LTX system in POPC:POPE:Chol (2:1:1).**

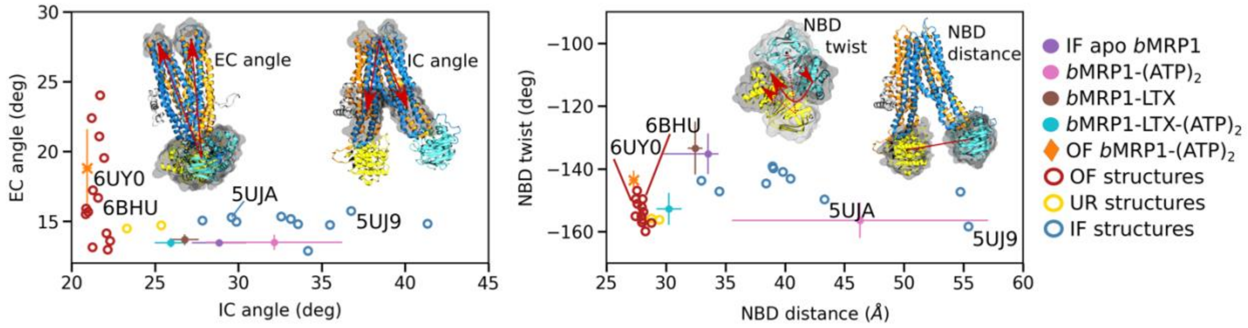

**Supplementary Figure 27. ABC conformational space of trajectories in POPC compared to resolved ABC transporters.**

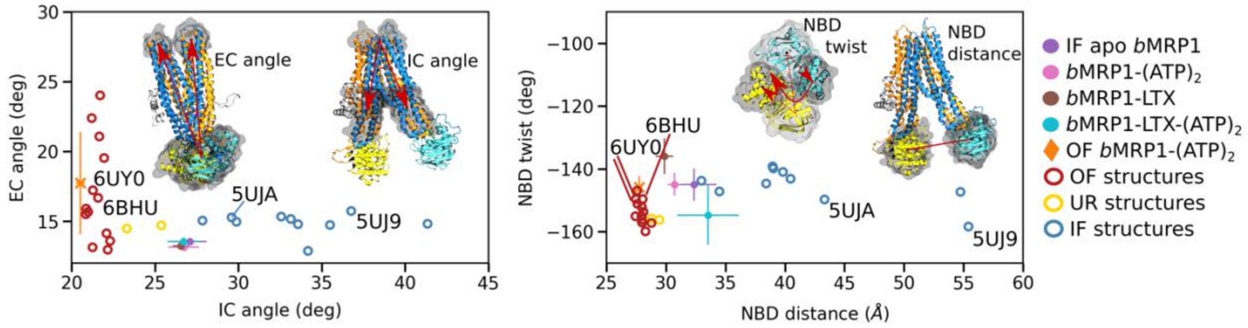

**Supplementary Figure 28. ABC conformational space of trajectories in POPC:Chol (3:1) compared to resolved ABC transporters.**

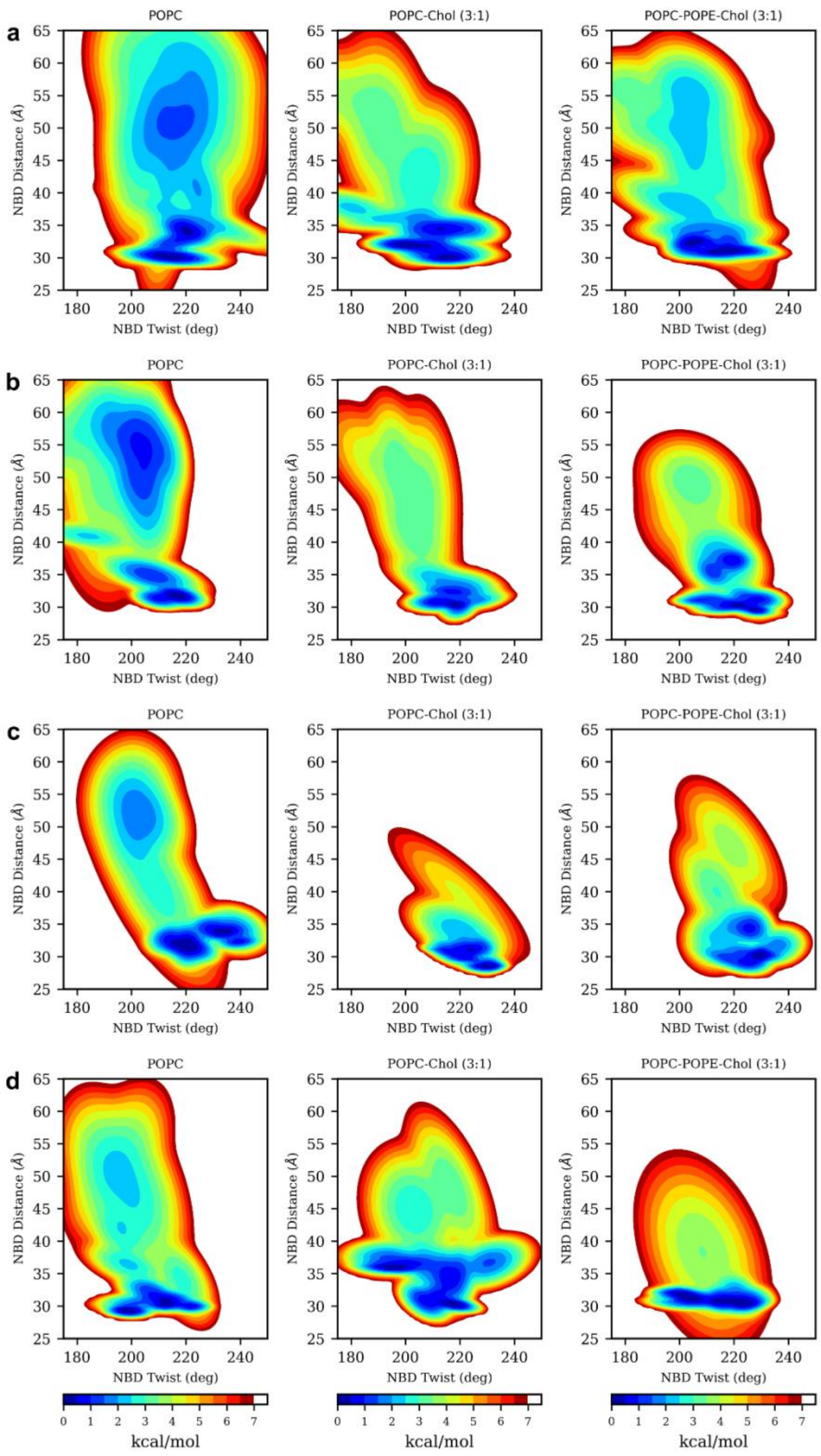

**Supplementary Figure 29. Free energy landscapes according to NBD distance and NBD twist. obtained in different lipid bilayers using the InfleCS approach for IF systems,** embedded in different lipid bilayers**. a)** IF apo *b*MRP1, **b)** *b*MRP1-$(ATP)_2$, **c)** *b*MRP1-LTX, **d)** *b*MRP1-LTX-$(ATP)_2$ and **e)** OF *b*MRP1-$(ATP)_2$.

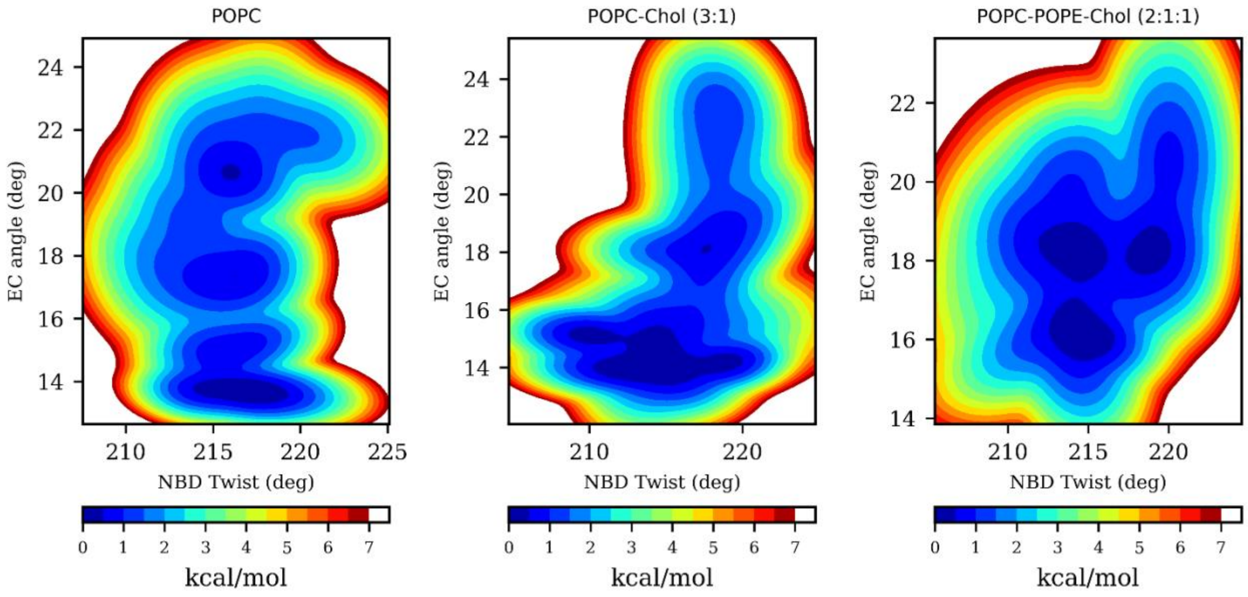

**Supplementary Figure 30. Free energy landscapes according to NBD twist and EC angle obtained in different lipid bilayers using the InfleCS approach for OF *b*MRP1-(ATP)$_2$ systems.**

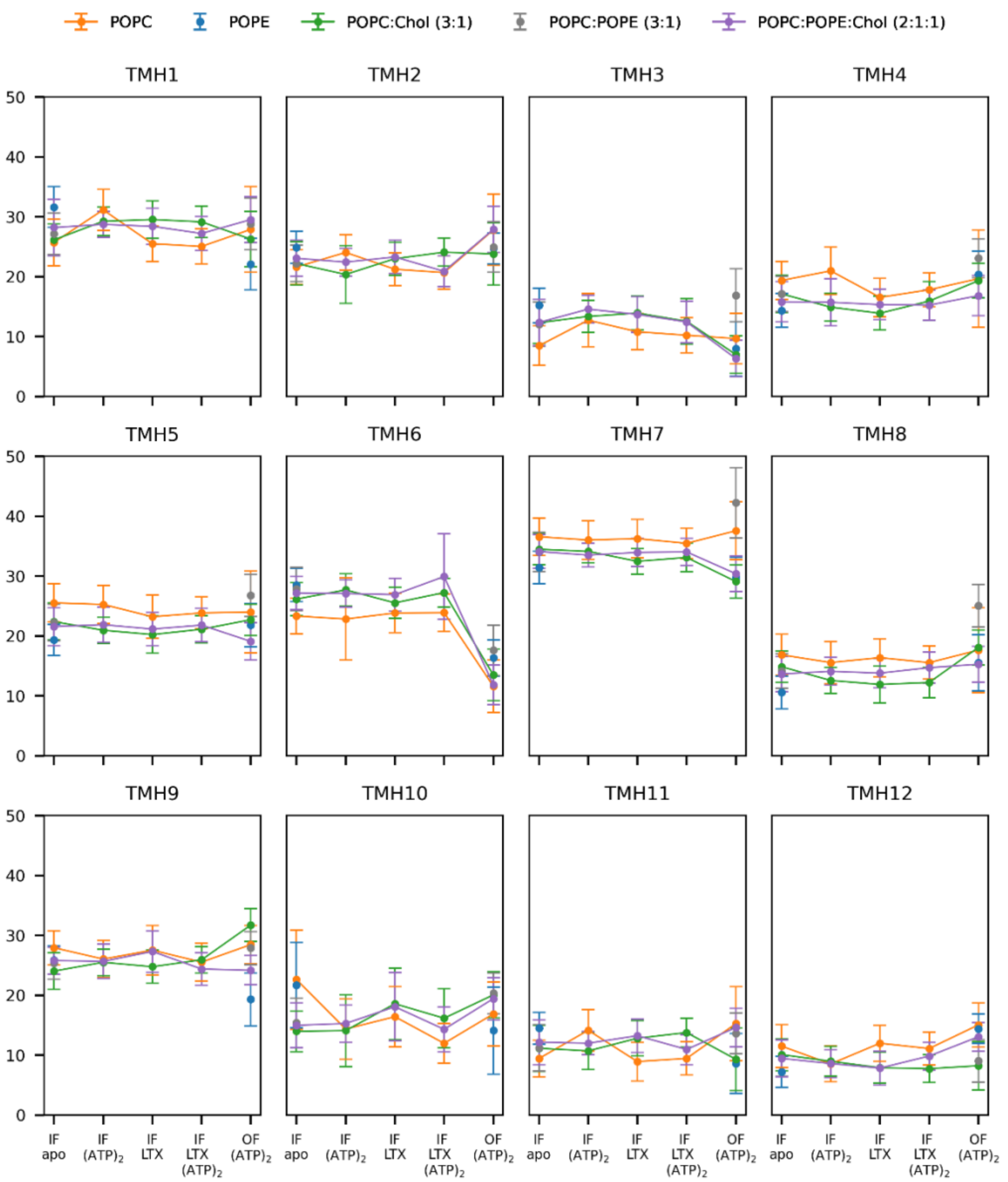

**Supplementary Figure 31. Tilt angles with respect to membrane normal axis of different transmembrane helices.** Since $TMD_0$ was not modelled in the present study, the conventional TMH1 to TMH12 labelling for type I ABC transporter was used.

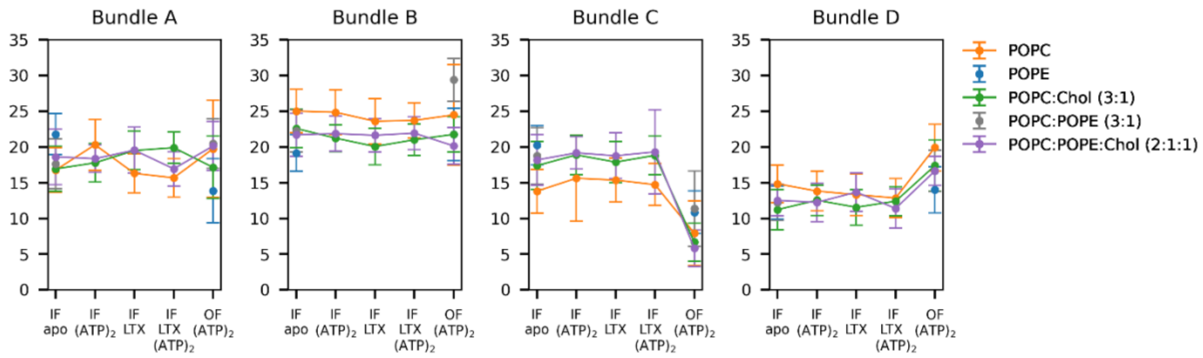

**Supplementary Figure 32. Tilt angles of bundles A, B, C and D which consist in (TMH1, 2, 10 and 11), (TMH4, 5, 7 and 8), (TMH 3 and 6) and (TMH9 and 12), respectively (see main text).**

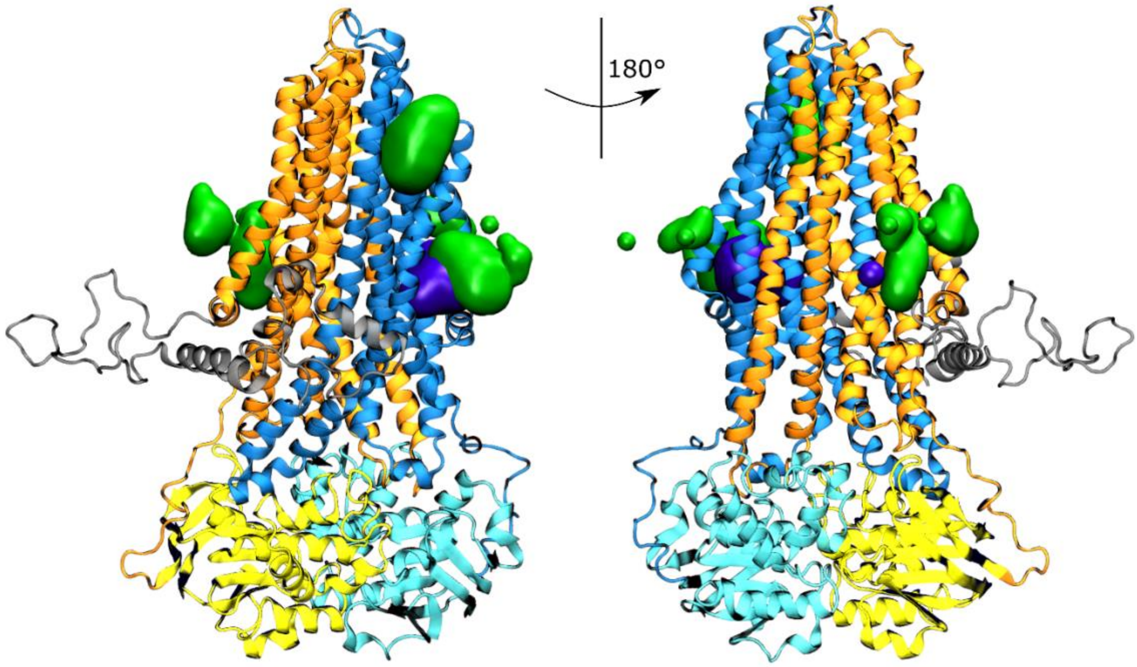

**Supplementary Figure 33.** Calculated binding hotspots obtained from cholesterol defined by presence likelihood higher than 50%. Cryo-EM resolved cholesterols are coloured violet, other cholesterols green.

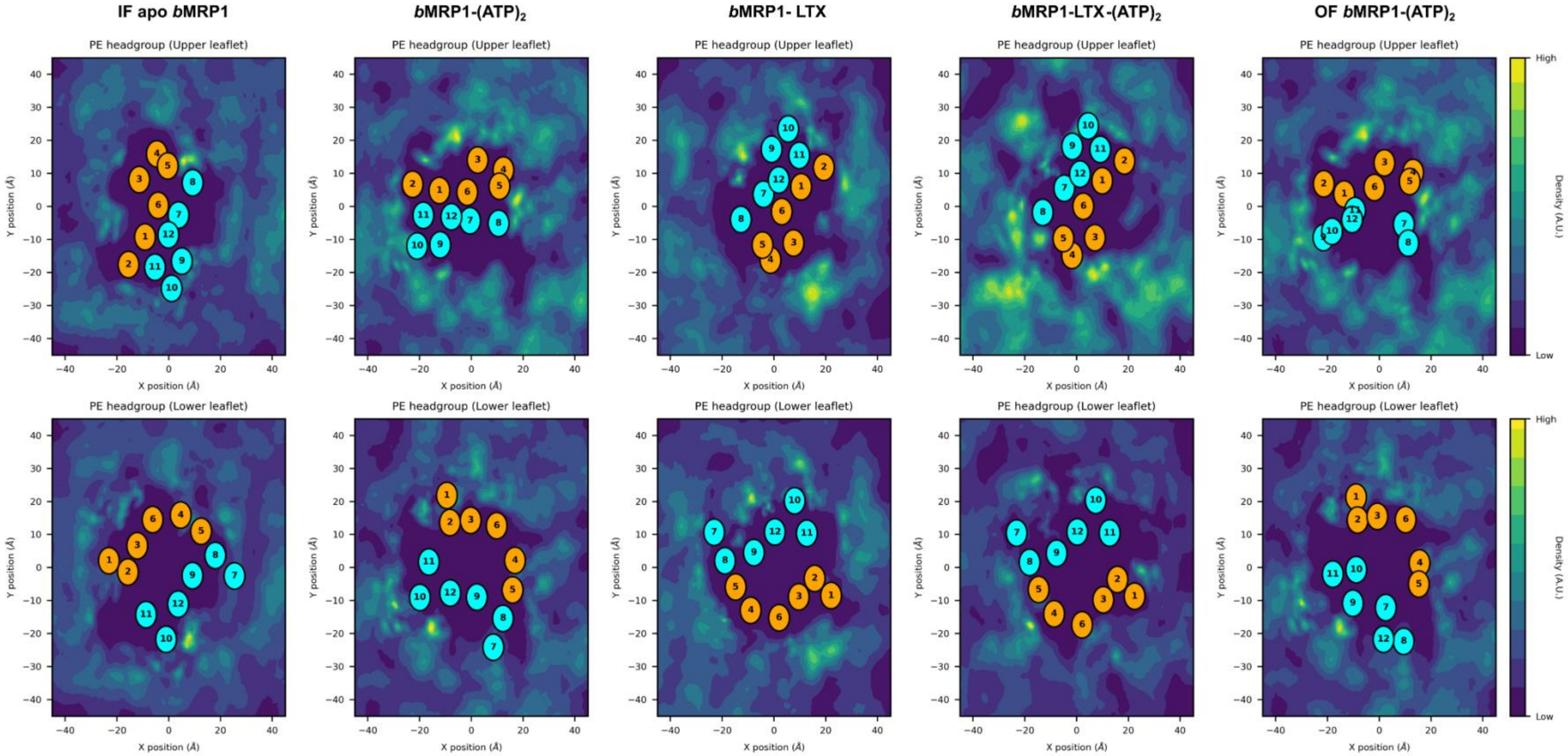

**Supplementary Figure 34. PE-lipid distribution in POPC:POPE:Chol (2:1:1) lipid bilayer.**

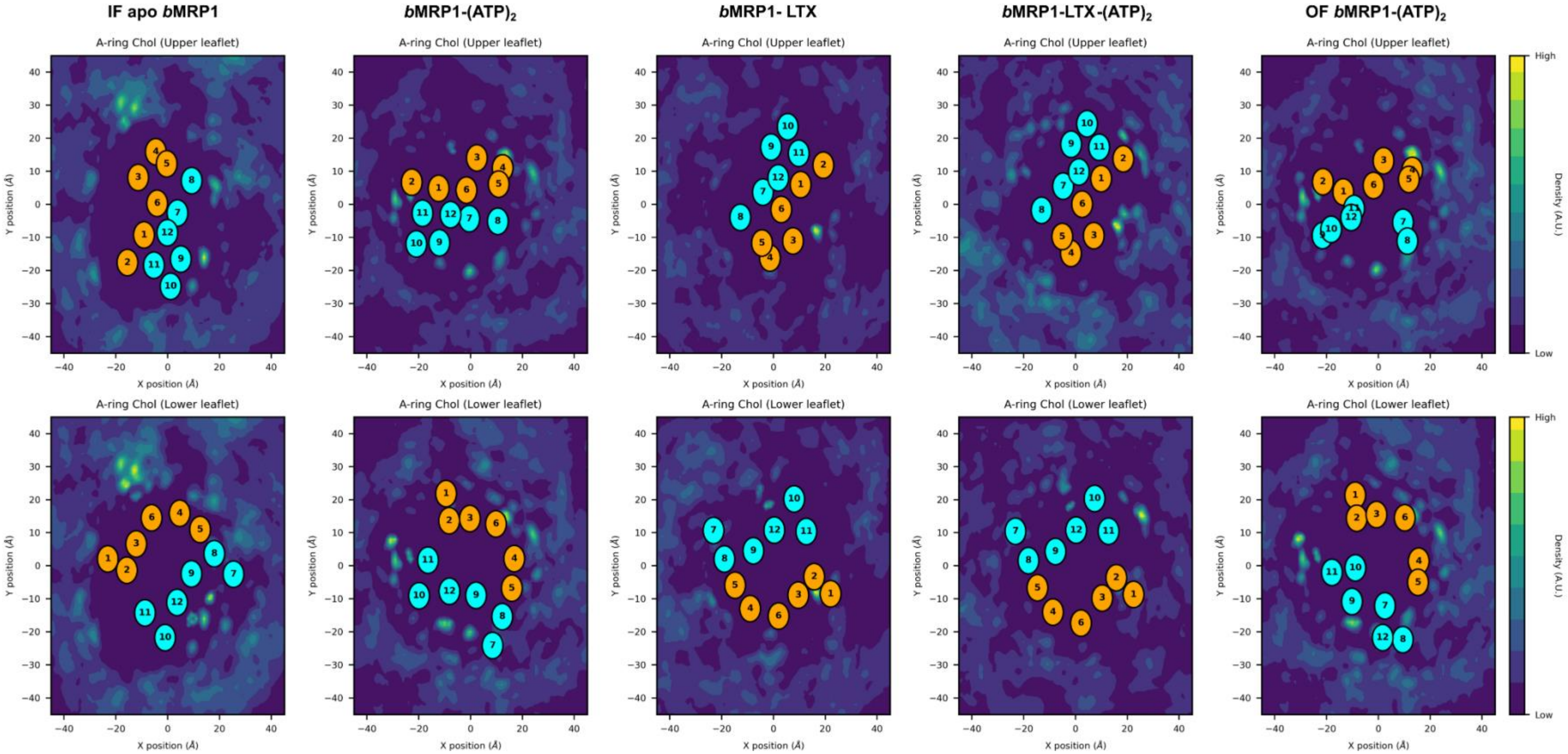

**Supplementary Figure 35. Cholesterol distribution in POPC:POPE:Chol (2:1:1) lipid bilayer.**

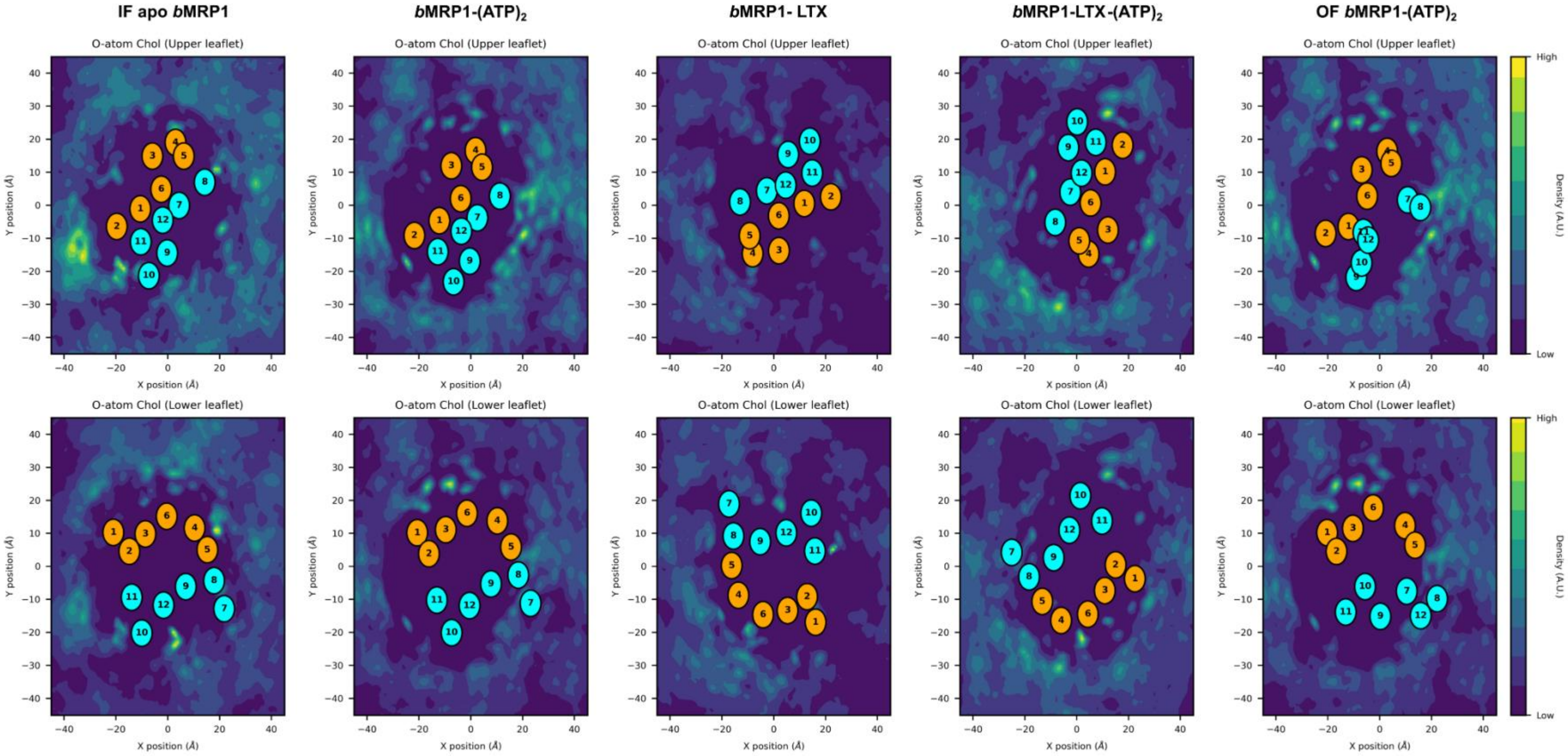

**Supplementary Figure 36. Cholesterol distribution in POPC:Chol (3:1) lipid bilayer.**

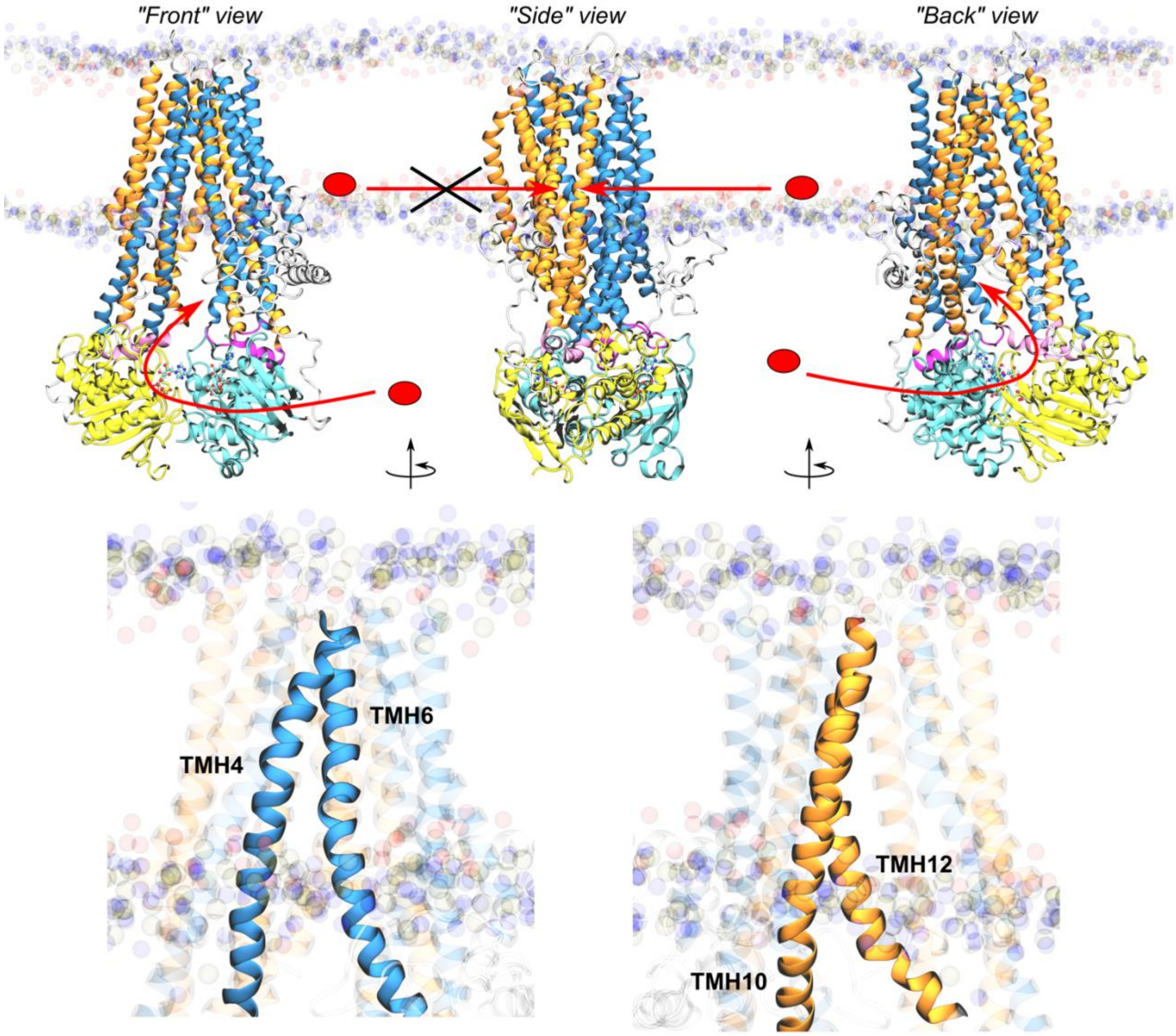

**Supplementary Figure 37. Putative substrate access to *b*MRP1 binding pocket.** We here propose different possible substrate access to *b*MRP1 binding pocket depending on substrate lipid bilayer partitioning. Amphiphilic compounds including charged molecules were shown to possibly partition within the high-density polar head region of lipid bilayer membrane. Our MD simulations suggest that access to TMD binding pocket may be possible between TMH4 and TMH6 while access between TMH10 and TMH12 might less likely.

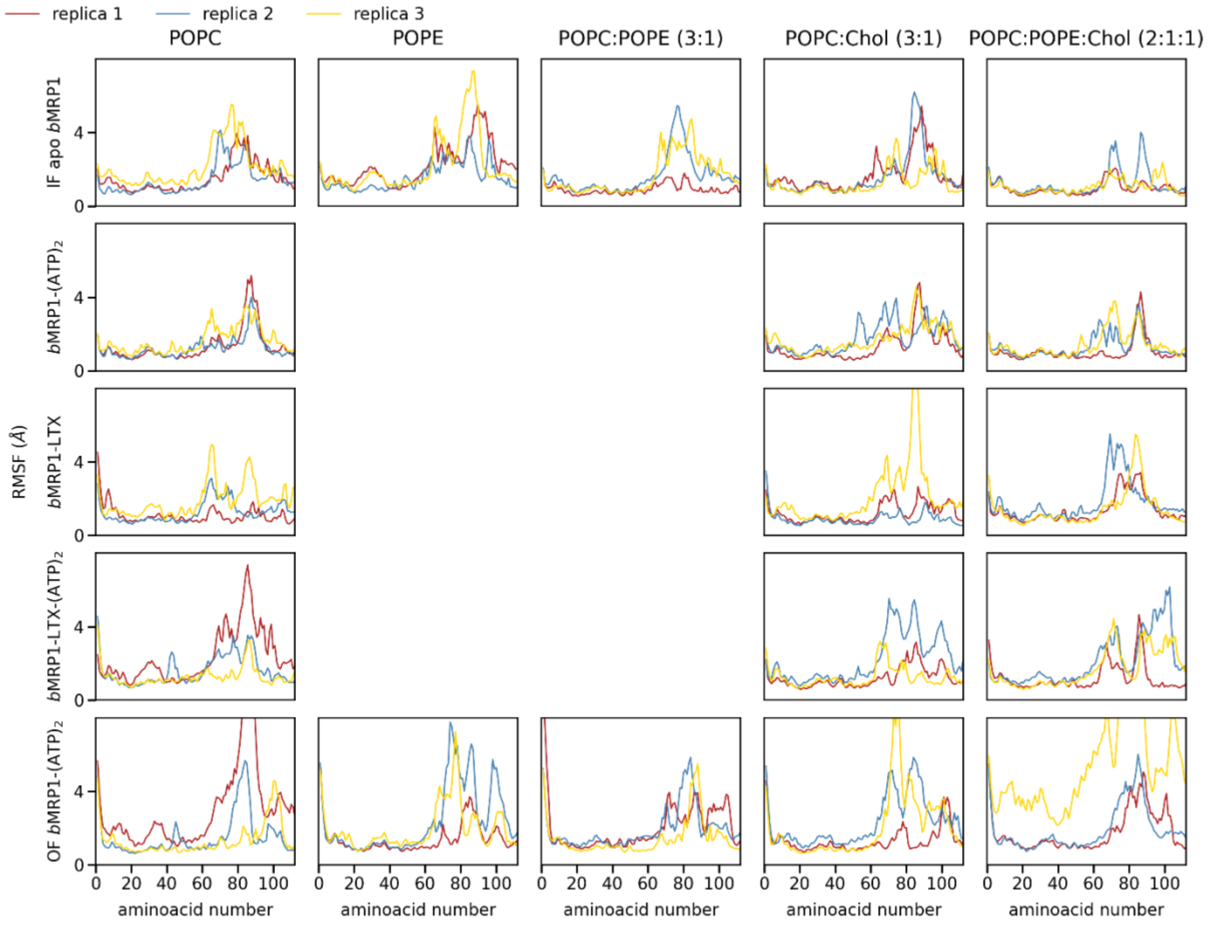

**Supplementary Figure 38. Root-mean square fluctuation of $L_0$.** The part which was modelled using the sequence and not the loop from the IF model (around 80-95) is the most flexible part in the IF models, as well.

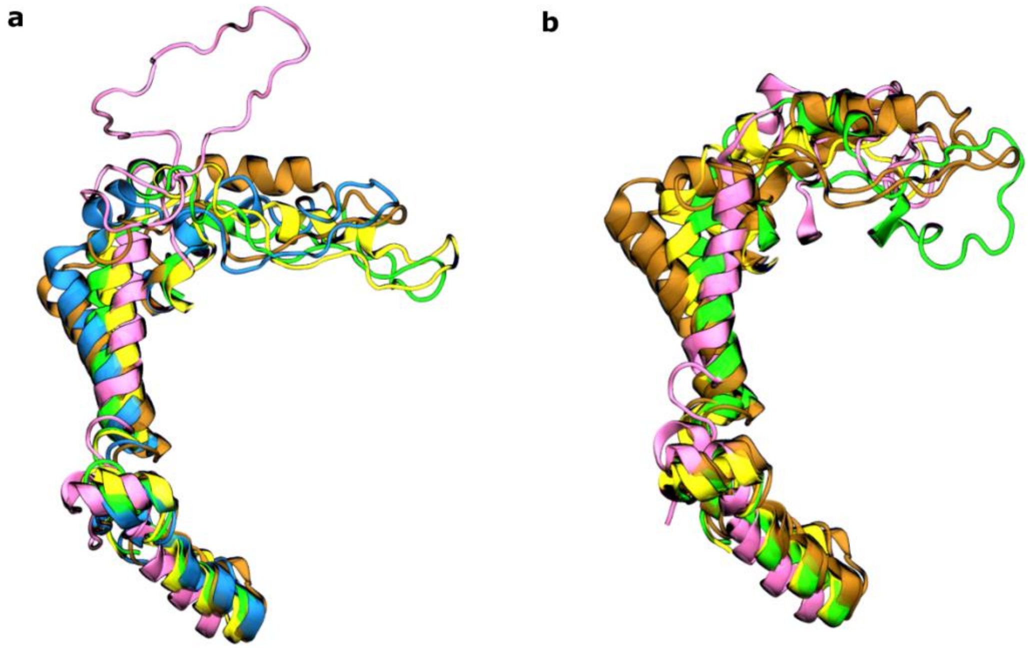

**Supplementary Figure 39. $L_0$ structures. a)** Starting frames converge to the same conformation shown by **b)** the final frames. Top view., aligned on TH1-3,6,10-11 (bundle A and C). IF apo is blue, IF ATP-bound ochre, IF LTX-bound yellow, IF LTX-ATP-bound green, and OF ATP-bound mauve/pink.

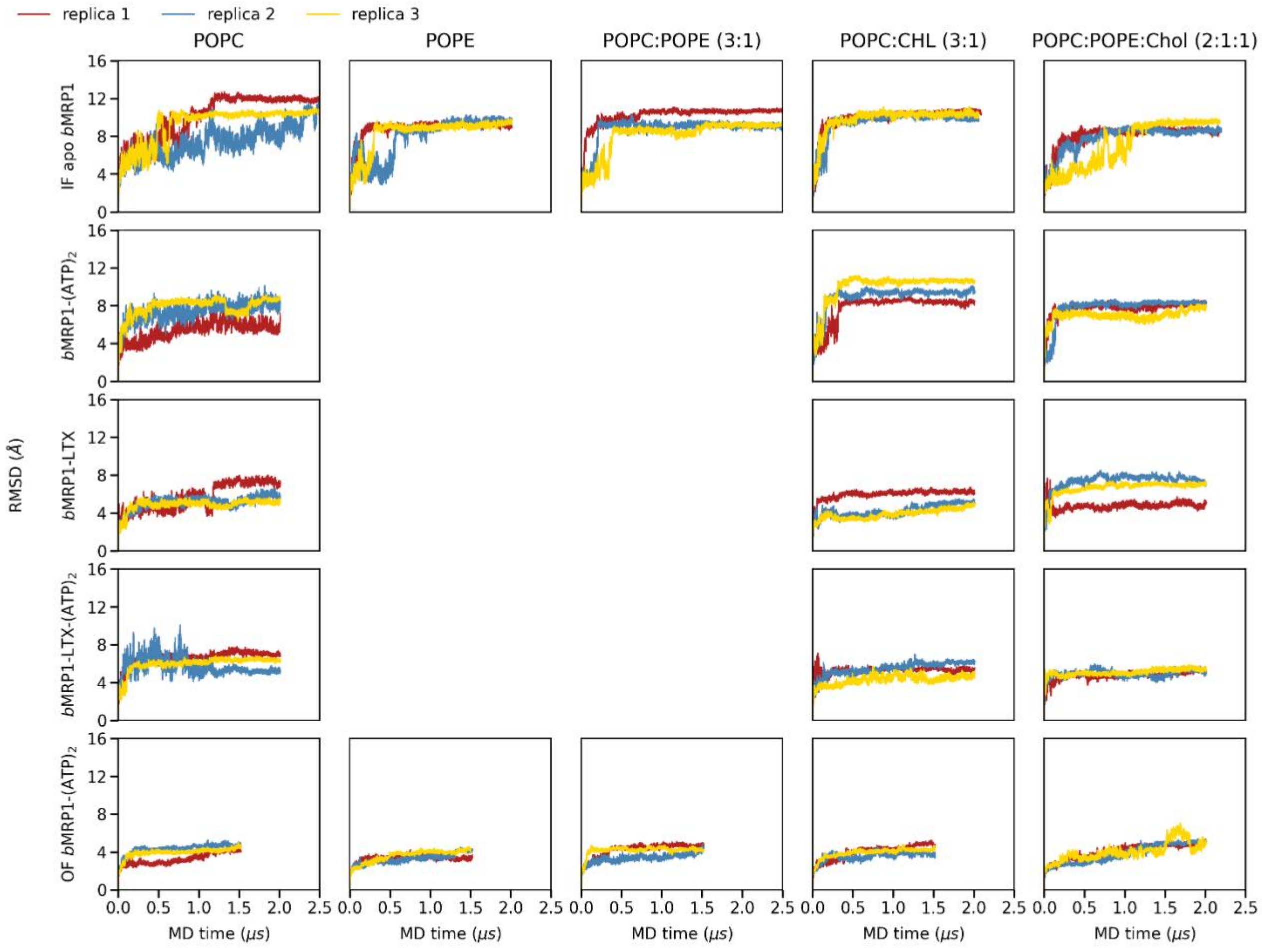

**Supplementary Figure 40. Root-mean-square deviation (RMSD) along MD simulations calculated for all the systems and for all different membrane compositions.** Replica 1, 2 and 3 are coloured red, blue and yellow, respectively.

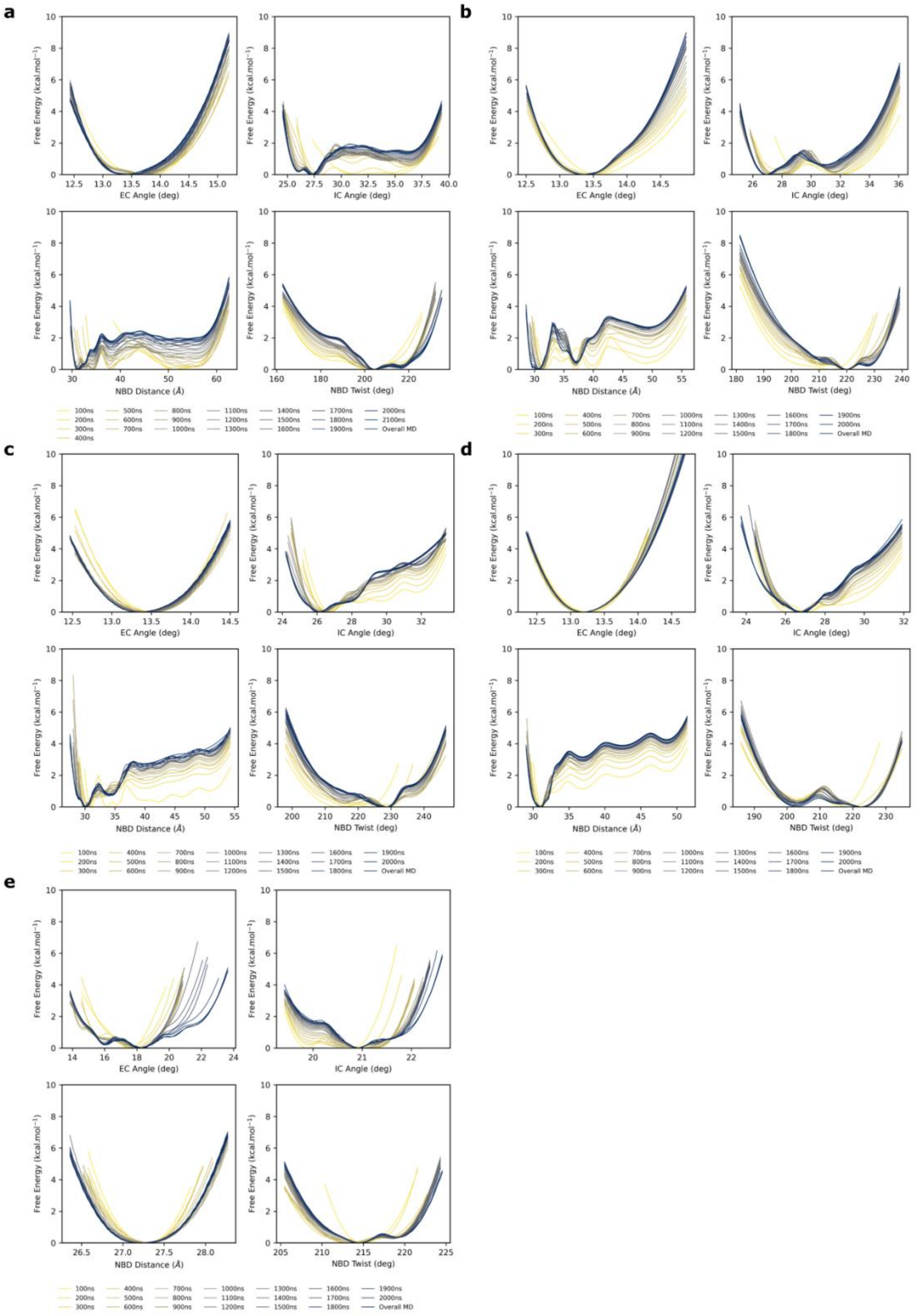

**Supplementary Figure 41. Convergence of InfleCS calculations in POPC:POPE:Chol (2:1:1).** **a)** IF apo *b*MRP1, **b)** *b*MRP1-(ATP)$_2$, **c)** *b*MRP1-LTX, **d)** *b*MRP1-LTX-(ATP)$_2$ and **e)** OF *b*MRP1-(ATP)$_2$.

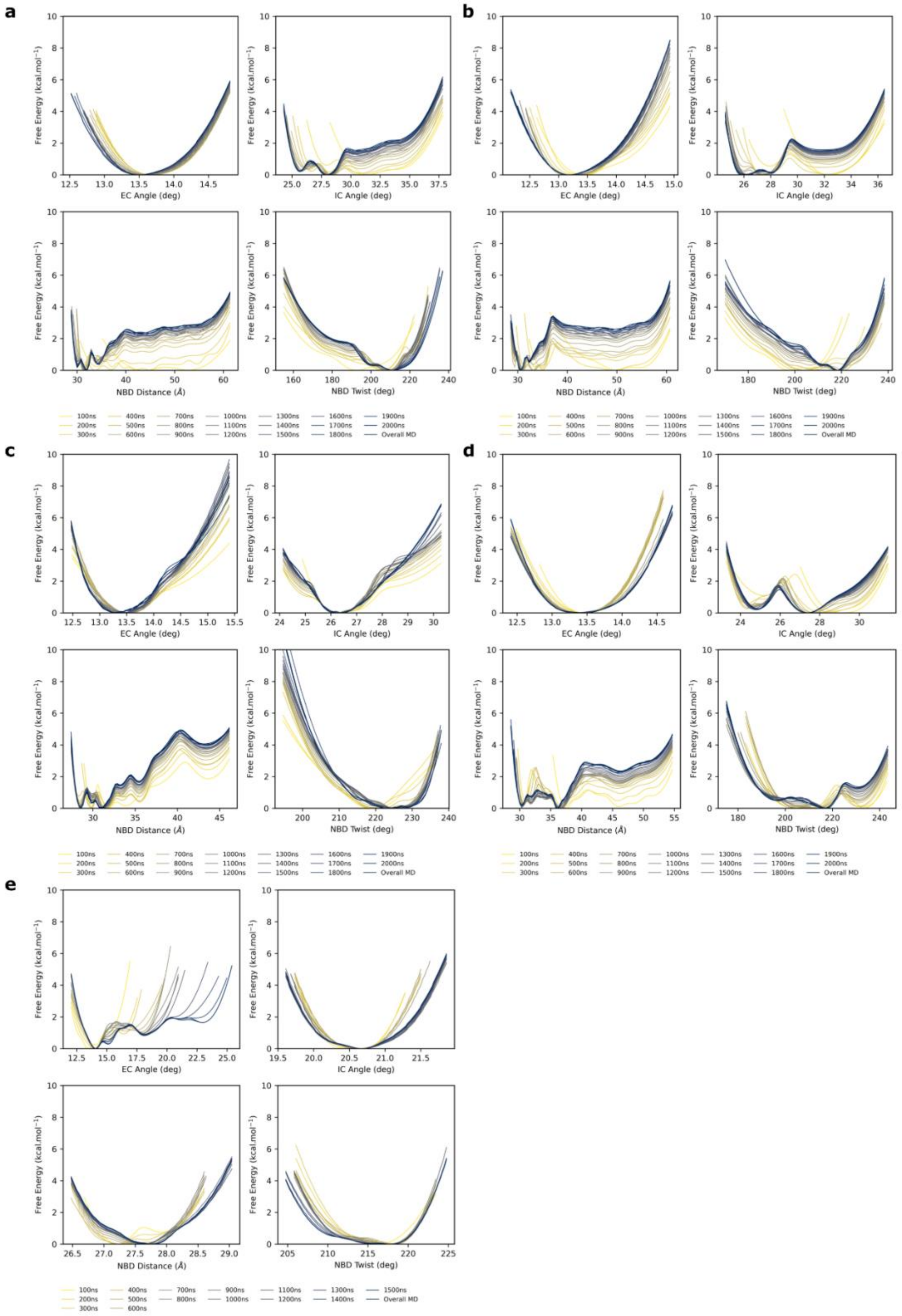

**Supplementary Figure 42. Convergence of InfleCS calculations in POPC:Chol (3:1). a)** IF apo *b*MRP1, **b)** *b*MRP1-$(ATP)_2$, **c)** *b*MRP1-LTX, **d)** *b*MRP1-LTX-$(ATP)_2$ and **e)** OF *b*MRP1-$(ATP)_2$.

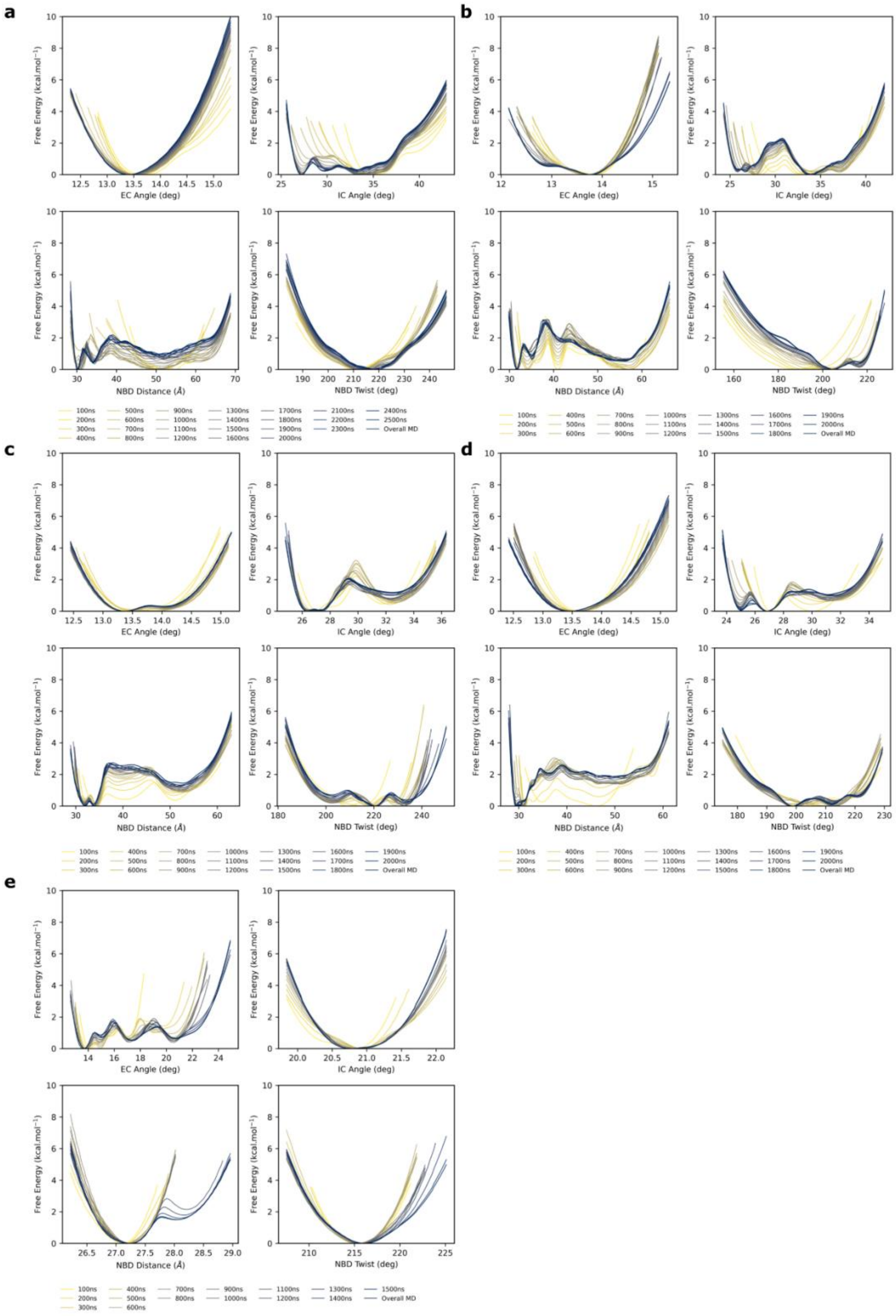

**Supplementary Figure 43. Convergence of InfleCS calculations in pure POPC. a)** IF apo *b*MRP1, **b)** *b*MRP1-$(ATP)_2$, **c)** *b*MRP1-LTX, **d)** *b*MRP1-LTX-$(ATP)_2$ and **e)** OF *b*MRP1-$(ATP)_2$.

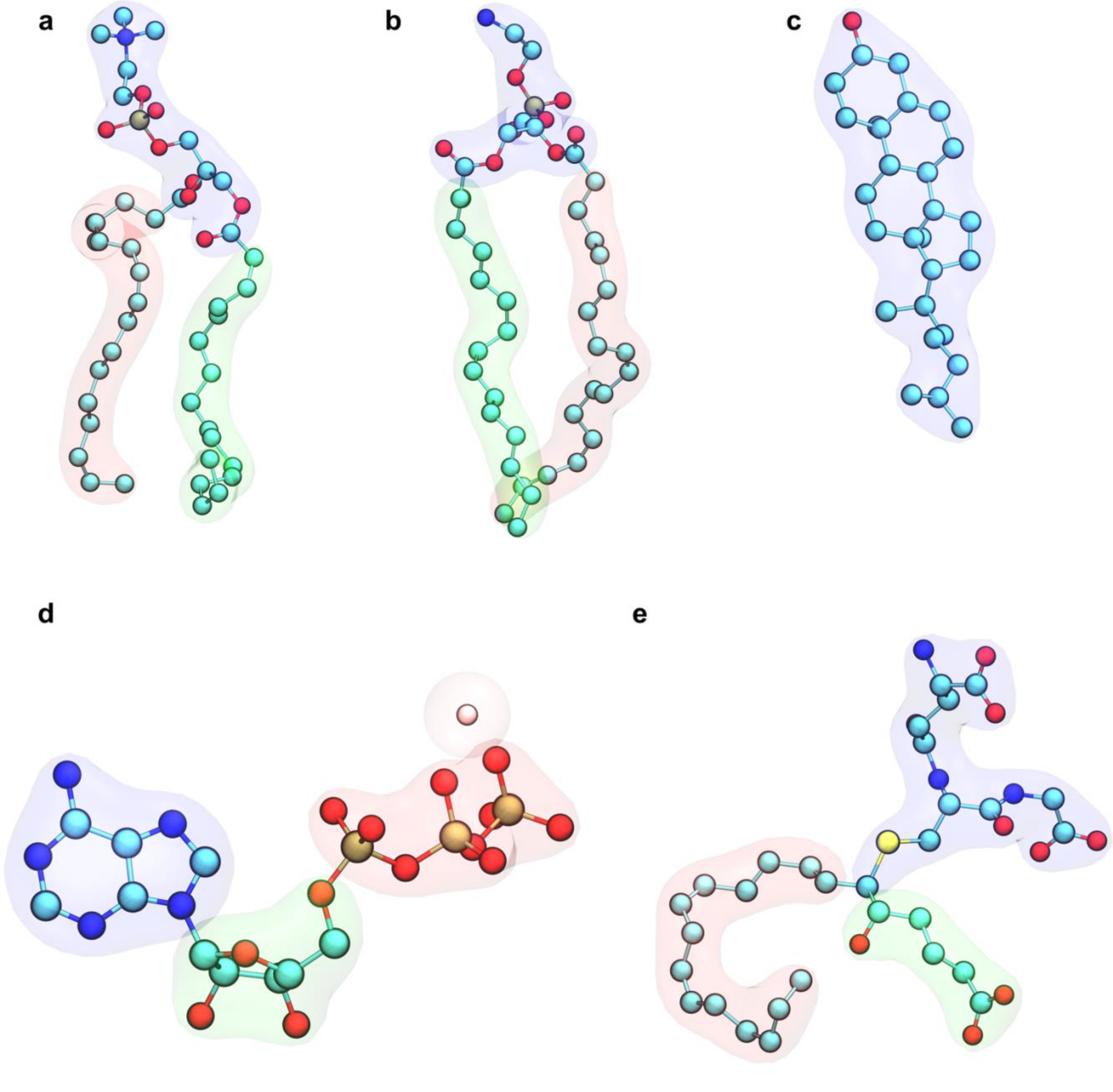

**Supplementary Figure 44. Atom selections for co-factor nodes for allosteric pathway calculations.** Each cofactor as split into fragments (depicted in blue, green, red and pink shades, accordingly) that were used to defined nodes for allosteric pathway calculations. PC and PE lipids were split into three nodes corresponding to polar head and lipid tails. Cholesterol and $Mg^{2+}$ were considered as single node each. ATP molecule was split into three fragments (purine and sugar moieties as well as phosphate tail). LTX was split into three fragments (namely, glutathione moiety, aliphatic chain and 6-ketohexanoate moiety).

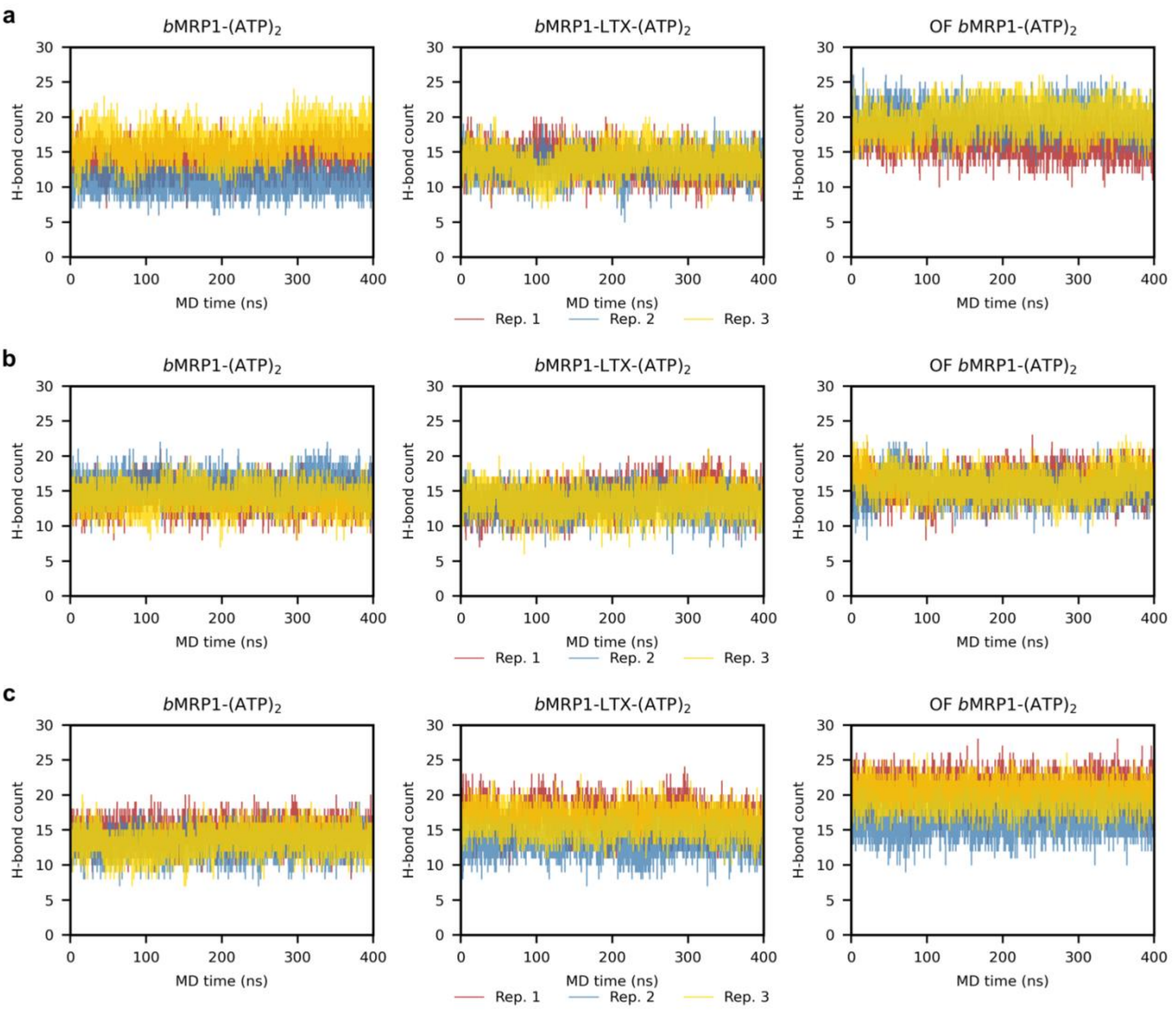

**Supplementary Figure 45. Number of H-bonds calculated between nucleotides and *b*MRP1 protein over MD simulations.** H-bond interactions were counted between nucleotides and protein residues over MD simulations performed on *b*MRP1-$(ATP)_2$, *b*MRP1-LTX-$(ATP)_2$ and OF *b*MRP1-$(ATP)_2$ systems embedded in **a)** POPC:POPE:Chol (2:1:1), **b)** POPC:Chol (3:1) and **c)** POPC. Distance and angle cutoffs were set 3.5 Å and 120°, respectively.

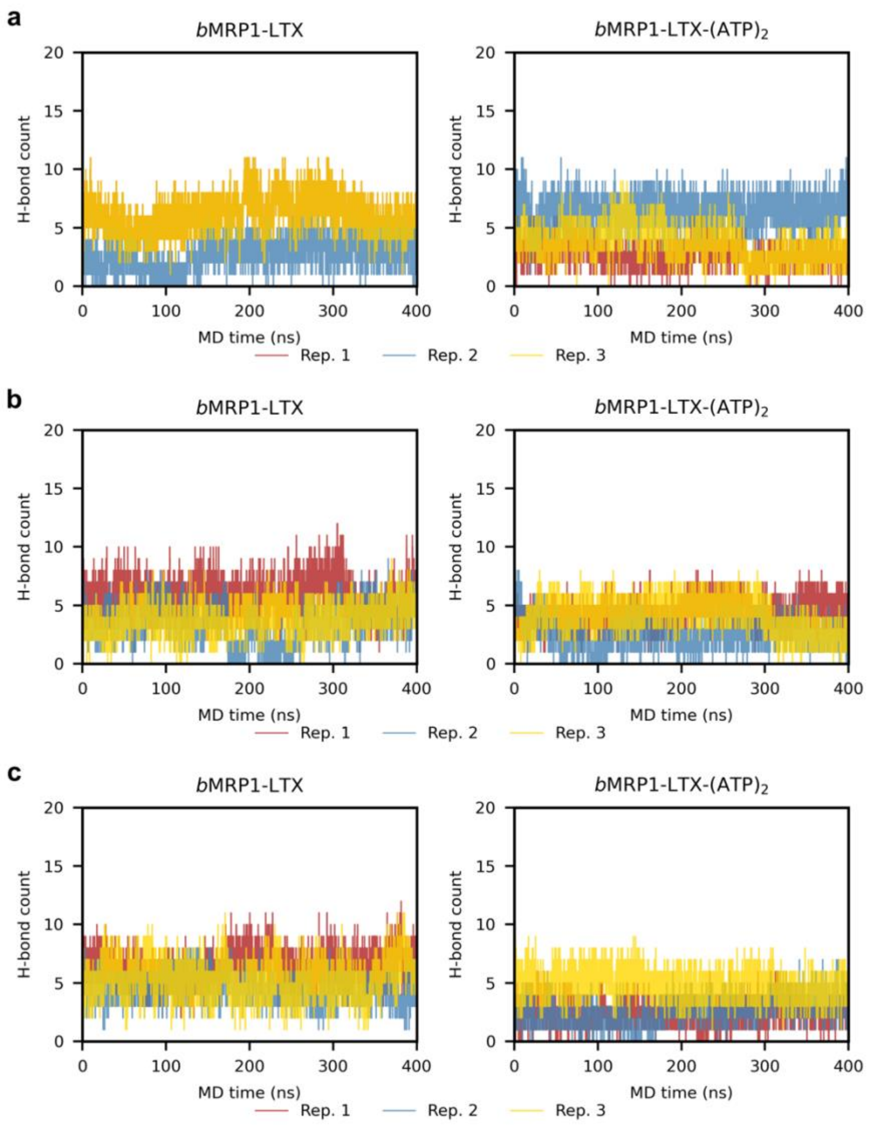

**Supplementary Figure 46. Number of H-bonds calculated between LTX and *b*MRP1 protein over MD simulations.** H-bond interactions were counted between LTX and proteins over MD simulations performed on *b*MRP1-LTX and *b*MRP1-LTX-$(ATP)_2$ systems embedded in **a)** POPC:POPE:Chol (2:1:1), **b)** POPC:Chol (3:1) and **c)** POPC. Distance and angle cutoffs were set 3.5 Å and 120°, respectively.

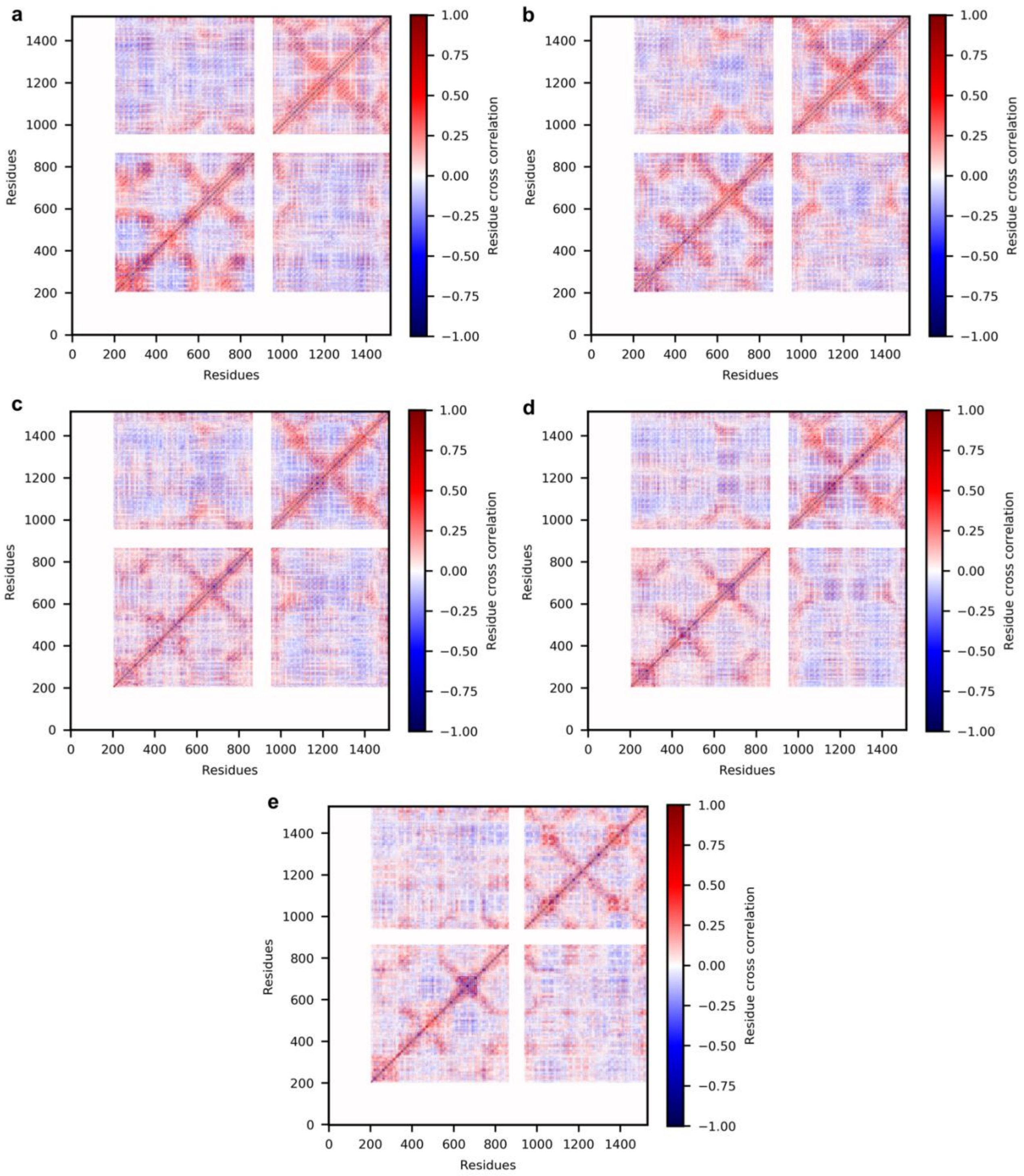

**Supplementary Figure 47. Overall dynamic cross-correlation matrices from MD simulations performed in POPC. a)** IF apo *b*MRP1, **b)** *b*MRP1-(ATP)$_2$, **c)** *b*MRP1-LTX, **d)** *b*MRP1-LTX-(ATP)$_2$ and **e)** OF *b*MRP1-(ATP)$_2$.

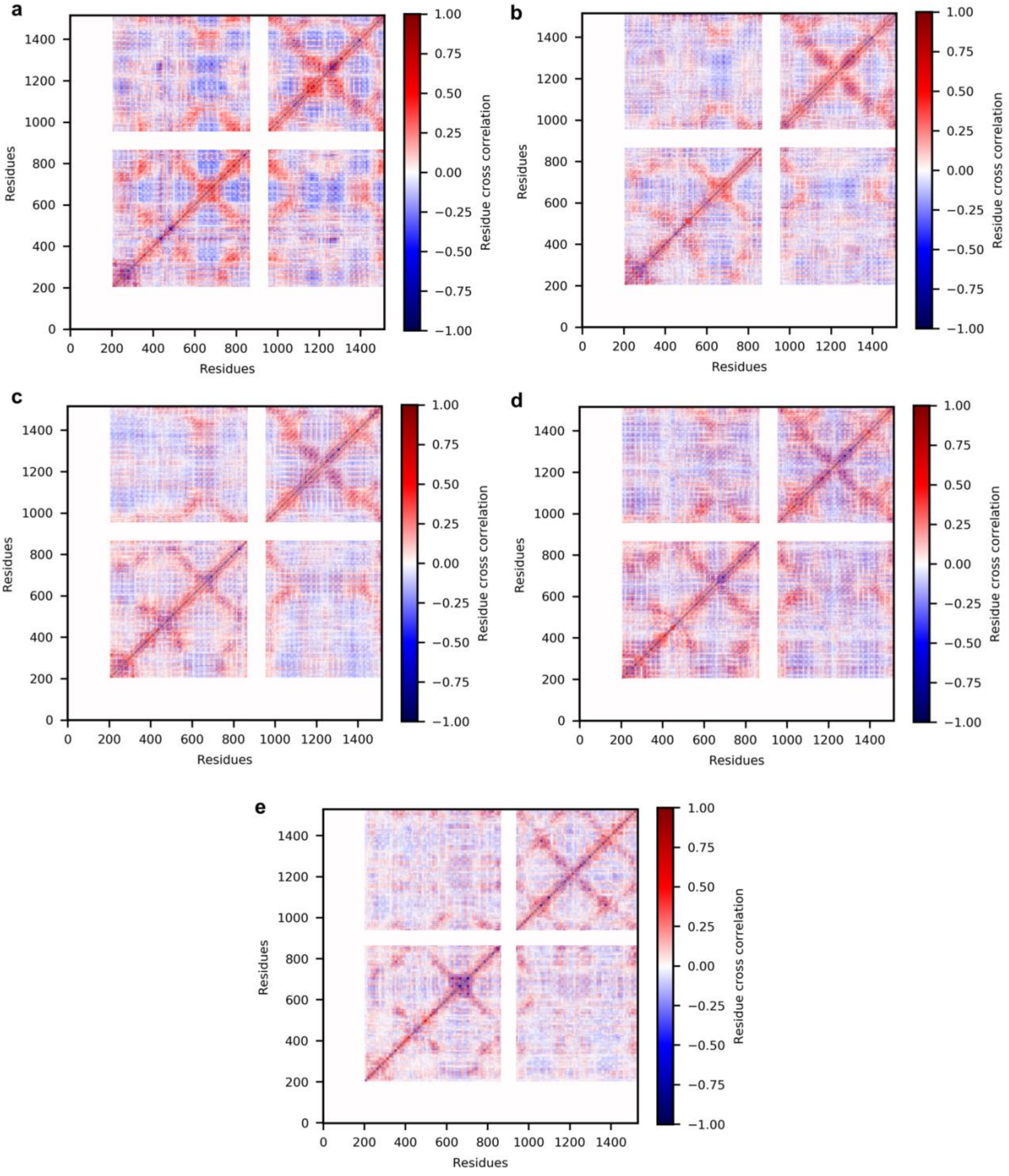

**Supplementary Figure 48. Overall dynamic cross-correlation matrices from MD simulations performed in POPC:Chol (3:1). a)** IF apo *b*MRP1, **b)** *b*MRP1-$(ATP)_2$, **c)** *b*MRP1-LTX, **d)** *b*MRP1-LTX-$(ATP)_2$ and **e)** OF *b*MRP1-$(ATP)_2$.

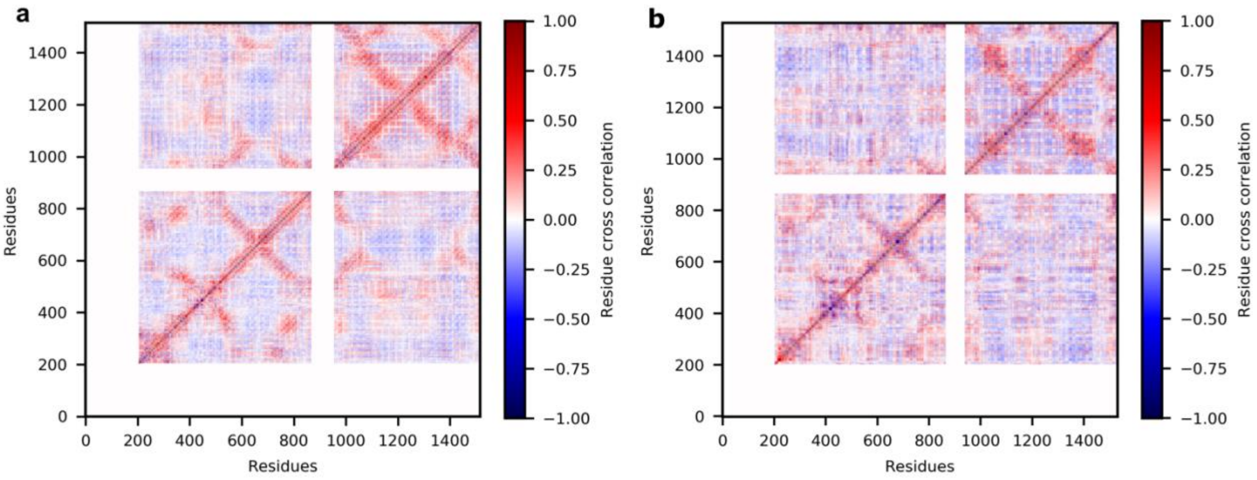

**Supplementary Figure 49. Overall dynamic cross-correlation matrices from MD simulations performed in POPC:POPE (3:1). a)** IF apo *b*MRP1 and **b)** OF *b*MRP1-(ATP)$_2$.

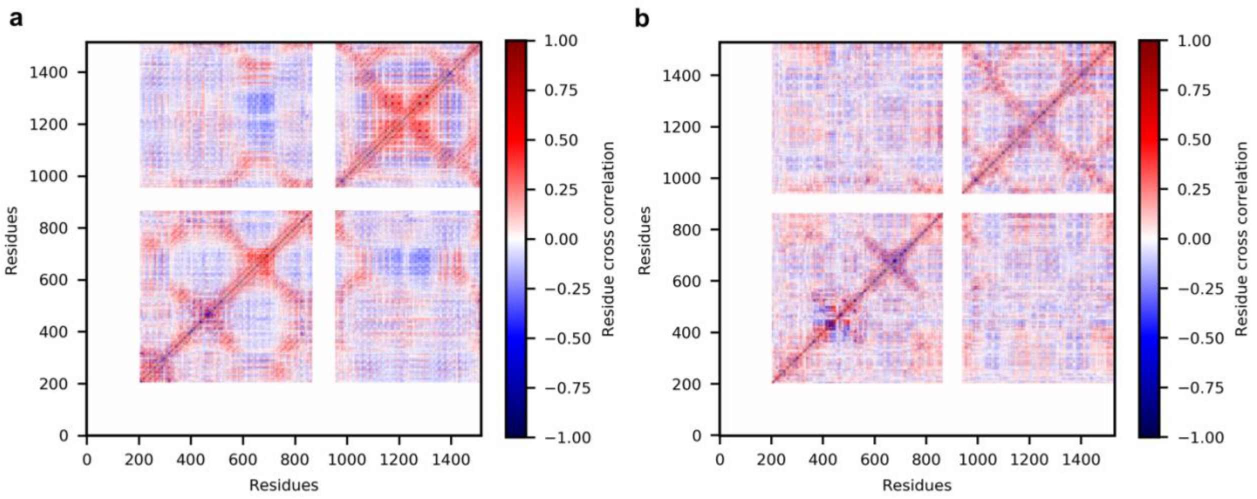

**Supplementary Figure 50. Overall dynamic cross-correlation matrices from MD simulations performed in POPE. a)** IF apo *b*MRP1 and **b)** OF *b*MRP1-(ATP)$_2$.

### Supplementary Movies

**Supplementary Movie 1. First principal component obtained from MD simulations performed on IF apo *b*MRP1 embedded in POPC:POPE:Chol (2:1:1). PCA were performed considering only the "so-called" ABC core, *i.e.*, TMHs and NBDs.**

**Supplementary Movie 2. First principal component obtained from MD simulations performed on IF *b*MRP1-(ATP)$_2$ embedded in POPC:POPE:Chol (2:1:1). PCA were performed considering only the "so-called" ABC core, *i.e.*, TMHs and NBDs.**

**Supplementary Movie 3. First principal component obtained from MD simulations performed on IF *b*MRP1-LTX embedded in POPC:POPE:Chol (2:1:1). PCA were performed considering only the "so-called" ABC core, *i.e.*, TMHs and NBDs.**

**Supplementary Movie 4. First principal component obtained from MD simulations performed on IF *b*MRP1-LTX-(ATP)$_2$ embedded in POPC:POPE:Chol (2:1:1). PCA were performed considering only the "so-called" ABC core, *i.e.*, TMHs and NBDs.**

**Supplementary Movie 5. First principal component obtained from MD simulations performed on OF *b*MRP1-(ATP)$_2$ embedded in POPC:POPE:Chol (2:1:1). PCA were performed considering only the "so-called" ABC core, *i.e.*, TMHs and NBDs.**
